# Infectious subgenomic amplicon strategies for Japanese encephalitis and West Nile viruses

**DOI:** 10.1101/2024.08.09.607374

**Authors:** Prince Pal Singh, Nguyen Phuong Khanh Le, Uladzimir Karniychuk

## Abstract

Classical methods for constructing infectious cDNA clones of flaviviruses are often hindered by instability and toxicity. The Infectious-Subgenomic-Amplicons (ISA) method is an advancement which utilizes overlapping DNA fragments representing viral genomic sequence and in-cell recombination to bypass bacterial plasmid assembly. However, the ISA method has limitations due to the toxicity of some ISA DNA fragments in bacteria during synthetic production. We validated modified ISA strategies for producing toxic ISA Japanese encephalitis virus (JEV) and West Nile virus (WNV) DNA fragments. Three approaches were explored including subdividing toxic DNA fragments into two sub-fragments for synthetic clonal production, using a low-copy bacterial plasmid, and subdividing the toxic DNA fragments into four short overlapping sub-fragments, each up to 1.8 kb. The latter novel approach in ISA applications enabled the synthesis of entirely bacteria-free ISA DNA fragments. Our results demonstrate that subdividing toxic fragments into sub-fragments smaller than 1.8 kb for synthesis is the efficient strategy, circumventing the need for bacterial plasmids and ensuring rapid production of synthetic flaviviruses. This method also shortens the production timeline. We also compared the efficacy of JEV and WNV ISA in zinc finger antiviral protein 1 (ZAP) wild-type and knockout cells and found that knockout cells may be more effective for ISA rescue of flaviviruses, including attenuated strains for live attenuated vaccines. The validated modified ISA strategies provide an efficient approach for producing synthetic JEV and WNV. This will enable rapid research during outbreaks of emerging flaviviruses by facilitating the quick generation of new virus variants.

## INTRODUCTION

Flaviviruses are single-stranded RNA viruses transmitted mainly by arthropods, as well as sexually and transplacentally. The genus Flavivirus (family *Flaviviridae*) comprises viruses which cause human disease, including dengue virus (DENV), Japanese encephalitis virus (JEV), and West Nile virus (WNV). These flaviviruses infect millions of people each year, causing symptoms from mild fevers to fatal hemorrhagic and neurological diseases [1]. The rising number of global flavivirus infections, frequent outbreaks, and expanding geographical range pose significant public health concerns [1]. Flaviviruses also pose constant threats to livestock industries [2]. There are no approved human vaccines for certain flaviviruses, such as Zika virus and WNV. Moreover, no licensed antivirals exist against flaviviruses.

There are four DENV serotypes which cause illnesses ranging from mild dengue fever to severe life-threatening dengue. Dengue virus is the major cause of hospitalization and death in children in Southeast Asian countries. Annually, DENV results in approximately 390 million infections, 100 million clinically apparent cases, and 500,000 severe cases, with at least 2.5 billion people at risk [1,3]. Over the past 70 years, infection rates have steadily increased, making DENV the most prevalent arthropod-borne viral disease worldwide. By comparison, only sporadic DENV epidemics were documented before the Second World War [1,4]. Between 2010 and 2022, more than 33,000 locally acquired dengue cases have been reported in the USA. Since 2014, there have been fewer than 1,000 cases per year, but 2022 was the first time in a decade that cases reached four digits. In 2023, Florida officials issued a warning about dengue as the cases continued to increase.

Japanese encephalitis virus causes encephalitis in humans in the Asia-Pacific region [5], with 68,000 annual cases and 15,000 deaths [6]. JEV epidemics were originally described in Japan in the 19^th^ century, and the virus was first recovered in 1935 from an infected human in Tokyo [1]. The enzootic cycle of JEV is between water birds and *Culex* mosquitoes, with pigs also serving as an amplifying host [1]. Humans are considered incidental dead-end hosts and generally do not produce viremia sufficient to infect mosquitoes. Though many human infections are asymptomatic, around 20-30% of clinical infections are fatal, and 30-50% of the survivors develop life-long neurological sequelae [5]. Human transplacental and fetal JEV infections are also reported [7]. The geographic range of emerging JEV keeps expanding from the South of Russia to Australia, Japan, Eastern China, India, and South-East Asia. Also, severe travel-related cases are reported around the globe [8]. There is a concern that JEV can be introduced into North America given a large population of susceptible avian species and amplifying hosts—pigs, wild boars; susceptible *Culex* mosquitoes are also ubiquitous [9–11]. In addition to morbidity and mortality in humans, JEV causes economically significant clinical disease in pigs, mostly abortions and stillbirth [11,12]. Japanese encephalitis virus keeps expanding—the 2022 JEV outbreak in Australia with infections in swine herds, zoonotic transmission, and human deaths was caused by genotype IV which was not associated with outbreaks before [13].

West Nile virus (WNV) is classified by the National Institute of Allergy and Infectious Diseases as a “Category B Priority” pathogen, indicating its high threat level. West Nile virus cycles between *Culex* mosquitoes and birds, but it can also infect humans, horses, and other mammals. The virus is widespread, found in Africa, the Middle East, West Asia, Australia, the Americas, and Europe. The first major European outbreak occurred in Romania in 1996, with the largest in 2018 affecting 12 countries. In the U.S., WNV emerged in 1999, rapidly spreading and causing significant morbidity and annual outbreaks, notably in 2002, 2003, and 2012 [1,14].

West Nile virus is neuroinvasive, leading to central nervous system damage, including meningitis and encephalitis, which can be fatal or cause long-term disabilities such as a poliomyelitis-like syndrome. The virus’s spread has increased due to expanding mosquito habitats and changing bird migration patterns caused by climate change. Currently, it is the most common insect-borne zoonotic virus in the U.S., with no approved human vaccines or antivirals. From 1999 to 2010, blood supply screening estimated 2 to 4 million total infections in the U.S. [15]. In 2022, Centers for Disease Control and Prevention reported 1,126 cases and 93 deaths in the U.S., and 2,406 cases and 1,599 neuroinvasive diseases in 2023, underscoring growing cases and inadequate environmental controls.

Reverse genetics, defined as the generation of viruses entirely from complementary DNA (cDNA) [16], has revolutionized virology research, including research of flaviviruses. This powerful technique enables the experimental editing of viral genomes to investigate effects of point mutations, host-pathogen interactions, virulence factors, mechanisms of immune evasion, and the development of reporter viruses, vaccines, and antiviral testing [17,18]. The classical method involves constructing an infectious DNA clone. This clone is typically a bacterial DNA plasmid or two plasmids containing the entire cDNA of the viral genome and a suitable promoter, such as the T7 promoter [17,19]. In this approach, viral RNA is obtained from biological specimens, and a double-stranded cDNA copy of the RNA virus genome is generated and incorporated into plasmids, which are then amplified in bacteria. After plasmid purification, infectious viruses are produced either by direct transfection of permissive cells or by transfection of genomic RNA obtained through *in vitro* transcription [17,19].

Despite advances in molecular techniques, classical methods for constructing infectious cDNA clones are time-consuming, unpredictable, and labor-intensive and frequently associated with undesirable mutations or unstable and toxic clones in bacteria [17,19]. Flavivirus infectious DNA clones are particularly challenging due to the instability and toxicity of some viral sequences in bacterial hosts [17–23]. To mitigate these difficulties, bacteria-free approaches have been developed, including long PCR, circular polymerase extension reaction, Gibson assembly, and Infectious-Subgenomic-Amplicons (ISA) [17].

The ISA method represents a significant advancement in reverse genetics for flaviviruses [24]. It allows to bypass the problematic step of assembling and amplifying the full-length flavivirus genome in bacterial plasmids and bacterial cells. Instead, the ISA process relies on in-cell recombination, which simplifies and shortens the procedure where overlapping PCR-amplified ISA DNA fragments, flanked with a pCMV promoter and HDR/SV40pA sequences, are directly transfected into mammalian cells, insect cells, or *in vivo* in mice [17,25]. For example, in the most recent study using the classical ISA the authors rescued DENV; in contrast cloning the full-length DENV genome in bacteria failed due to toxicity [19].

While ISA is termed a “bacteria-free” approach, it is not completely free of bacteria and associated toxicity issues. Specifically, to our knowledge all sixteen published studies utilizing ISA for flavivirus rescue have used PCR amplification of fragments from pre-existing full-length flavivirus DNA infectious clones, infected biological specimens, or synthetic overlapping DNA ISA fragments. These later synthetic DNA fragments, provided by commercial suppliers, are assembled, inserted into bacterial plasmids, and amplified in bacteria [19,24–38]. Although this approach avoids constructing unstable and toxic plasmids containing sequences of the entire viral genome, some synthetic ISA DNA fragments still may pose bacterial toxicity, slowing or halting their commercial production. Indeed, in several of these papers, along with commercially generated ISA DNA fragments, the authors PCR-amplified some ISA DNA fragments directly from historically preexisting entire-genome-length flavivirus infectious DNA clones or from infected biological specimens to rescue yellow fever virus, DENV, and JEV [24,34]. While the reason for applying this mixed, not entirely synthetic reverse genetics approach was not explicitly explained in these publications, it is likely due to the toxicity of ISA DNA fragments preventing commercial generation of fragments for fully synthetic flaviviruses.

In our studies, we encountered similar technical difficulties with the toxicity of flavivirus ISA DNA fragments, which impeded commercial suppliers’ ability to provide the required fragments. Given the importance of generating entirely synthetic flavivirus stocks for certain experimental objectives, in this study we applied and validated modified ISA strategies. These approaches allowed us to generate JEV and WNV using a completely synthetic method for toxic ISA flavivirus fragments.

## MATERIALS AND METHODS

Wild-type and CCCH-type zinc finger antiviral protein (ZAP) knockout cells, *in silico* permutation of the viral genome, ISA, DENV, JEV, and WNV stock generation and titration, comparative ISA, mouse experiments, RT-qPCR, and statistics are described in supplemental materials and methods.

## RESULTS

### Classical Infectious-Subgenomic-Amplicons method for dengue virus

As described before [19], transfection of three overlapping ISA DNA fragments generated via *synthetic clonal genes* with medium copy number bacterial plasmids (**Table S1**) and amplification in bacteria (**Fig 1**) consistently rescued infectious DENV in all five well replicates. The virus induced a cytopathic effect (CPE) in all transfected wells at passages 0 and 1 (**Fig 2A**), as well as in flasks during subsequent passaging and virus stock generation. Sanger sequencing confirmed correct sequences in the DENV stock. The DENV stock produced in VERO-ZAP-KO cells exhibited high infectious titers when titrated on both VERO-ZAP-WT and VERO-ZAP-KO cells (**Fig 2B**), along with high viral RNA loads (**Fig 2C**) and E protein expression (**Fig 2D**).

**Figure 1.**
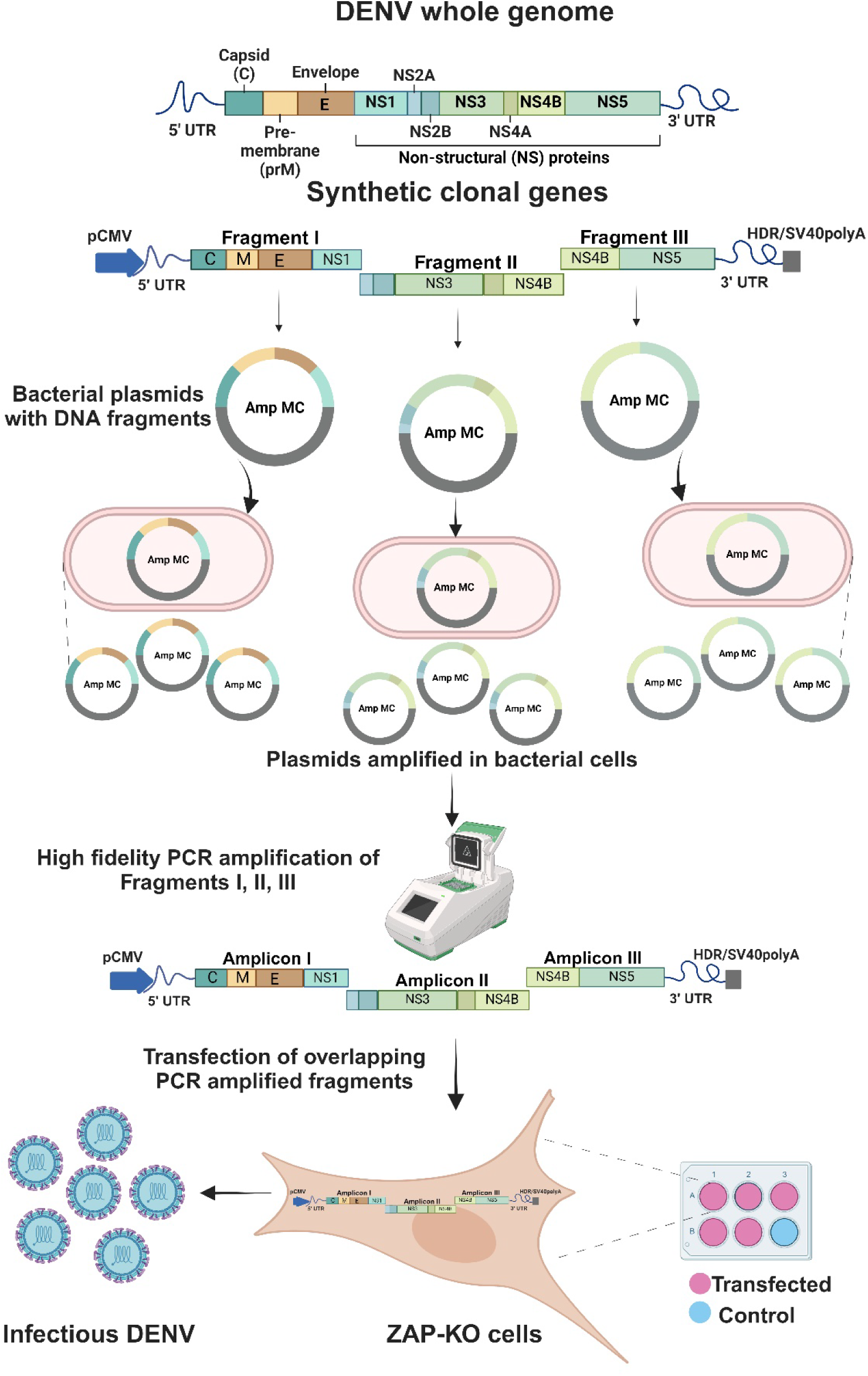
Classical ISA method for dengue virus.

**Figure 2.**
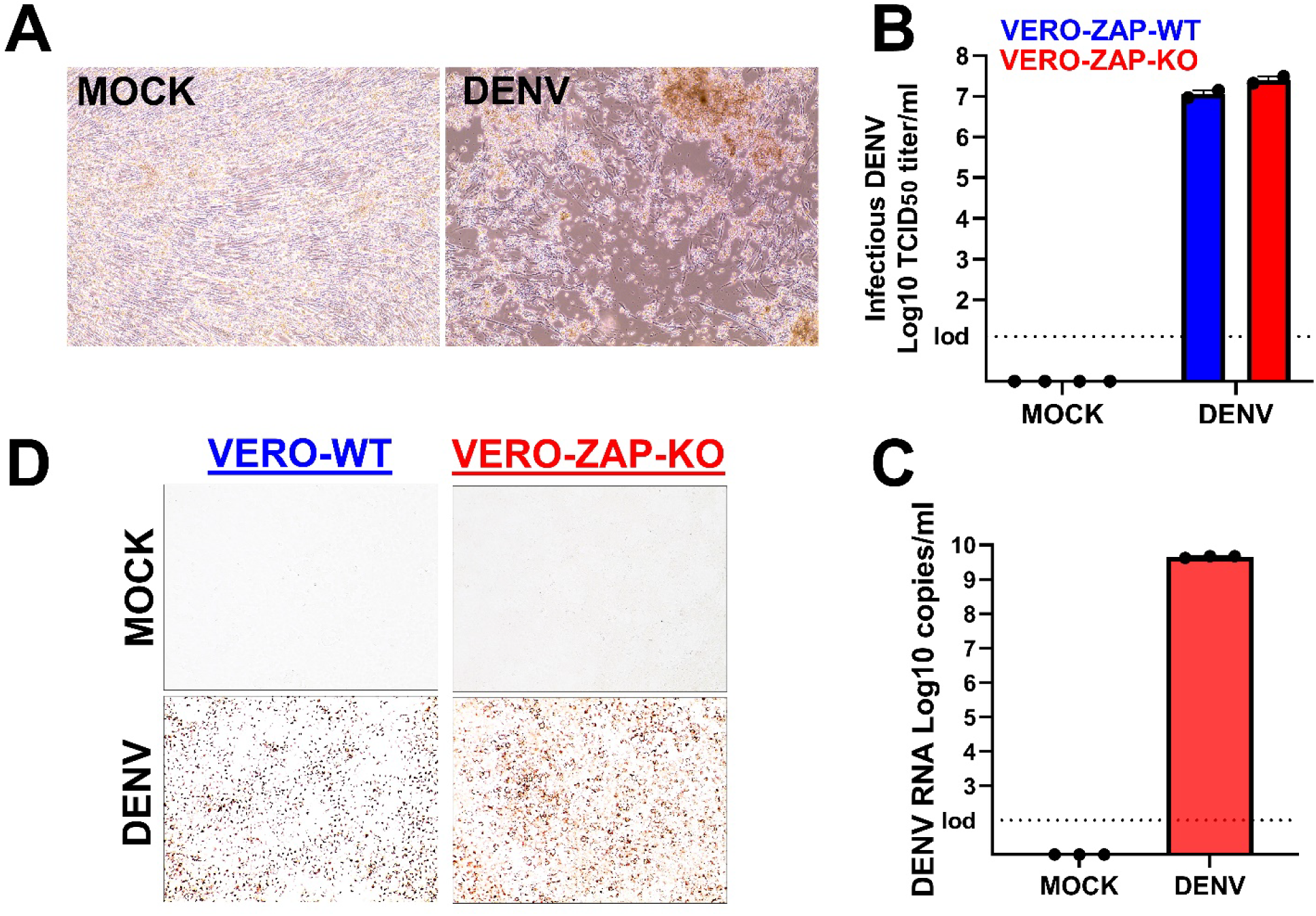
Dengue virus ISA and stock generation. (**A**) Cytopathic effect in DENV ISA- transfected ZAP-KO BHK+VERO cells, passage 0 day 7 after transfection. All five replicate wells showed CPE at passages 0 and 1; the representative images are shown. (**B**) Infectious TCID_50_ titers of the DENV stock produced in ZAP-KO cells. The stock was titrated in VERO-ZAP-WT or VERO-ZAP-KO cells. (**C**) DENV RNA loads of the stock determined by virus-specific RT-qPCR. (**D**) Images of cells positive for the DENV E protein (red staining) at 7 days after infection with MOI 0.01. Magnification x200. The experiment was done in 4 replicates. Figures represent the general patterns in all replicates.

Altogether, as in the previous study [19], we confirmed that classical ISA (**Fig 1**) with three overlapping DNA fragments generated via *synthetic clonal genes* with bacterial plasmids is a highly efficient and robust technique for type 2 DENV reverse genetics, enabling the rescue of synthetic recombinant viruses. The properties of ISA-derived DENV have been extensively characterized in prior research [19]; thus, we further focused on the more challenging ISA strategies for JEV and WNV.

### Infectious-Subgenomic-Amplicons method to overcome bacterial toxicity caused by Japanese encephalitis virus sequences

In contrast to DENV, classical ISA failed to rescue the JEV SA14-14-2 wild-type strain (JEV- WT). The JEV Fragment I exhibited bacterial toxicity across three different medium and low copy bacterial plasmids (**Table S1, Fig 3**). To overcome this, we divided the toxic Fragment I sequence (4,550 bp) into two sub-fragments, Fragment-I-A and Fragment-I-B, each 2,315-2,316 bp. These sub-fragments were produced via *synthetic clonal genes* using medium copy number bacterial plasmids and amplified in bacteria (**Table S1, Fig 3, File S1**). Transfection of four overlapping ISA DNA fragments generated via *synthetic clonal genes* (**Fig 3**) consistently rescued infectious JEV-WT in all five replicates. The virus induced CPE in all transfected wells at passages 0 and 1 (**Fig 4A**), as well as in flasks during subsequent passaging and virus stock generation. Sanger sequencing confirmed correct sequences in the JEV-WT stock. The JEV-WT stock produced in VERO-ZAP-KO cells exhibited high infectious titers when titrated on both VERO-ZAP-WT and VERO-ZAP-KO cells (**Fig 4B**), along with high viral RNA loads (**Fig 4B**) and E protein expression (**Fig 4C**).

**Figure 3.**
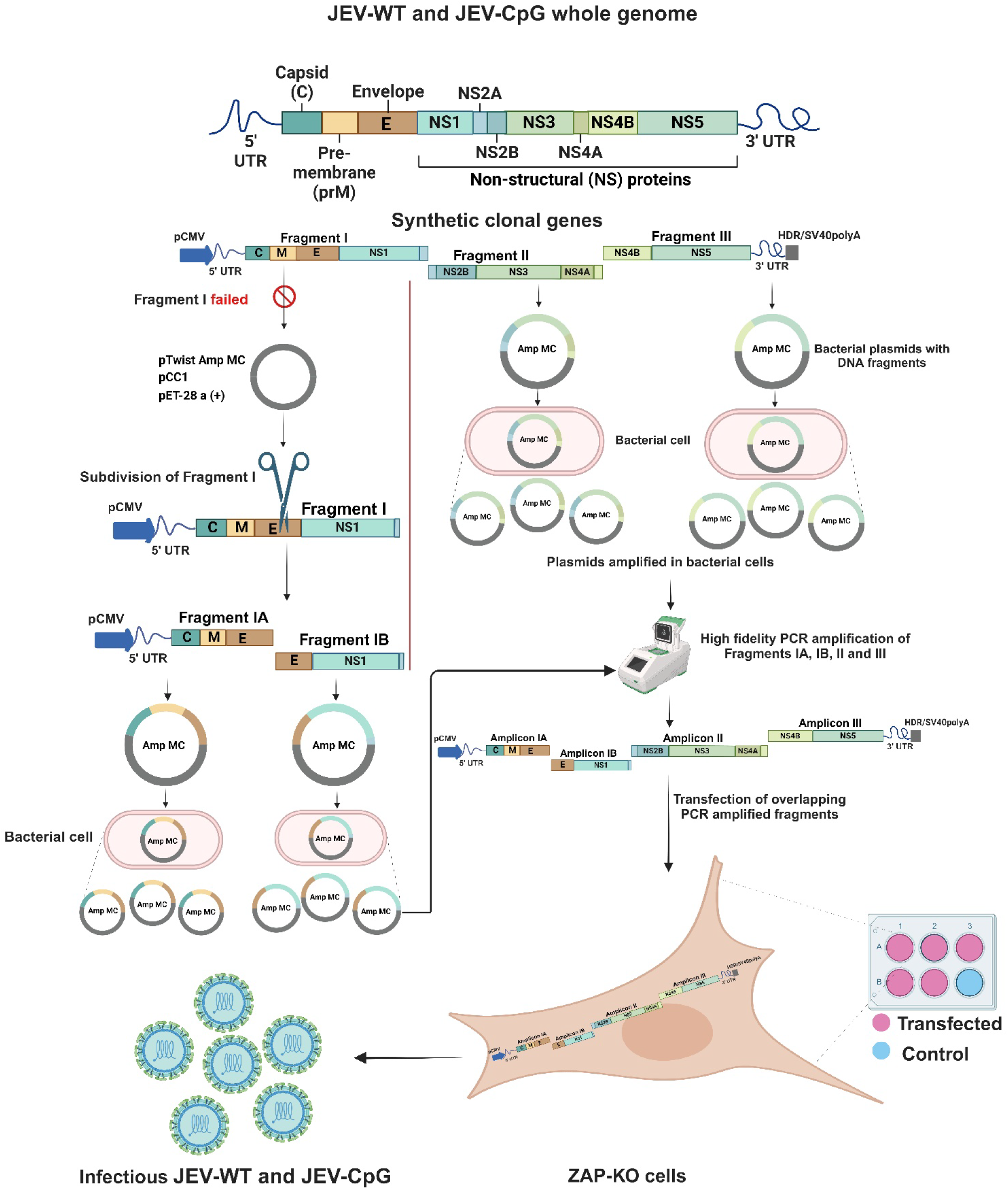
The adapted ISA method for Japanese encephalitis virus. To overcome toxicity, we divided the toxic Fragment I into two sub-fragments (**Table S1**, **File S1**).

**Figure 4.**
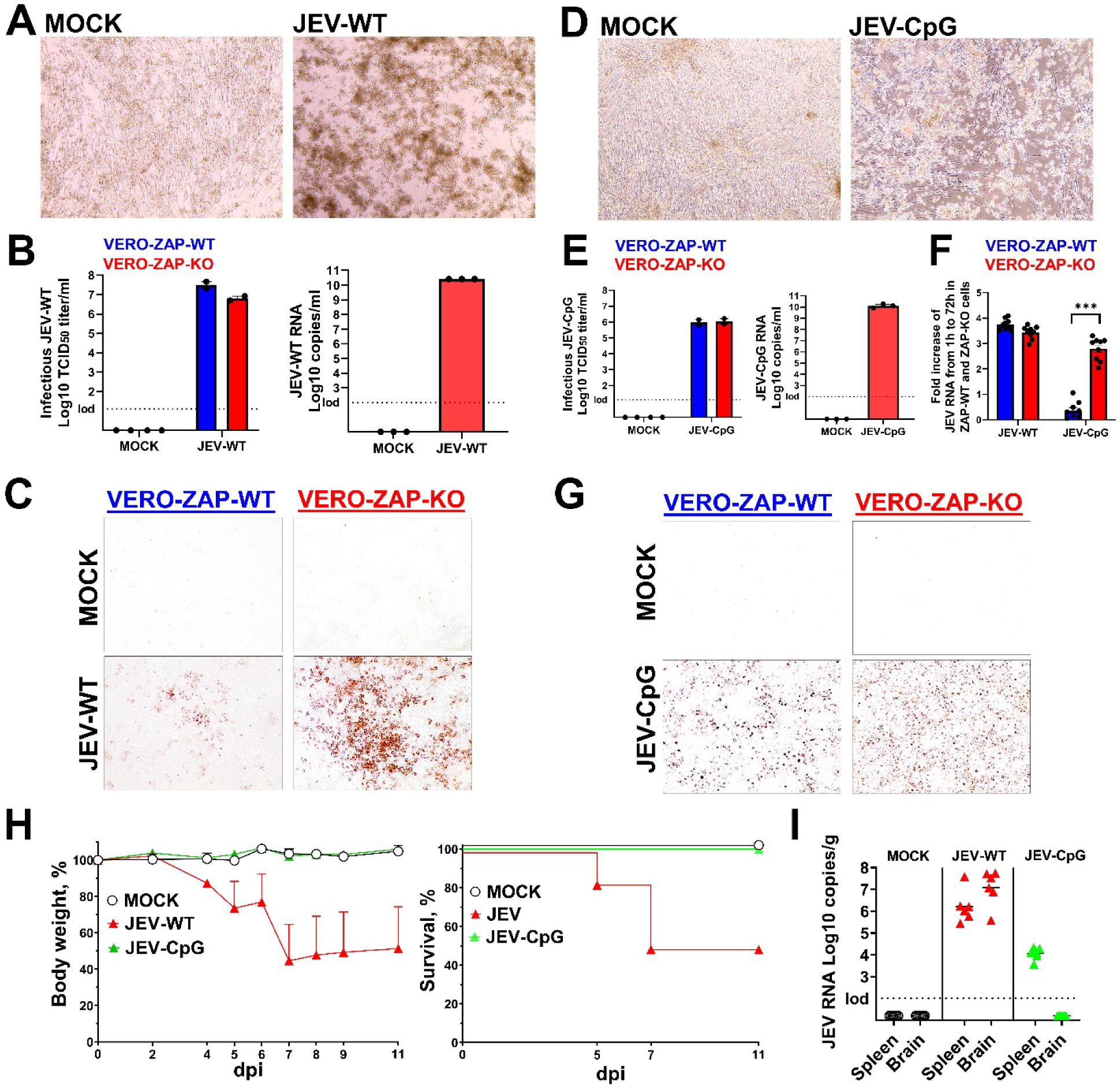
JEV ISA and stock generation. Cytopathic effect in JEV-WT (**A**) and JEV-CpG (**D**) ISA-transfected ZAP-KO BHK+VERO cells, passage 0, day 6 after transfection. All five replicates showed CPE; the representative images are shown. Infectious TCID_50_ titers of the JEV-WT (**B**) and JEV-CpG (**E**) stocks produced in ZAP-KO cells. The stock was titrated in VERO-ZAP-WT or VERO-ZAP-KO cells. JEV RNA loads of the stock determined by virus-specific RT-qPCR. Images of cells positive for the JEV-WT (**C**) and JEV-CpG (**G**) E protein (red staining) at 3 days after infection, MOI 0.01. Magnification x200. The experiment was done in 3 biological replicates. Figures represent the general staining patterns in replicates. (**F**) The fold increase of JEV-WT or JEV-CpG RNA loads in cell supernatants collected at 1h and 72h after inoculation. VERO-WT and VERO-ZAP-KO cells were inoculated with MOI 0.01. Virus inoculums were removed and replaced with media. Supernatants were collected at 1h (to normalize for leftover inoculum RNA) and at 72h. Three biological x 3 technical replicates. **\*\*\****P*=0.0017; unpaired t-test. (**H**) Mean body weight and survival in MOCK *Ifnar1^-/-^* mice and *Ifnar1^-/-^* mice injected with the ISA-derived JEV- WT or JWV-CpG. The following clinical scoring was used: 0—no visible abnormalities; 1—mild ataxia or tremors; 2—obvious ataxia or tremors; 3—depression, hunching, reluctance to walk, and falling to the side when walking; 4—close to moribund but still somewhat responsive; 5— paralysis; 6—loos of 20% baseline body weight; 7—found dead. Scores 4, 5, or 6 were used for the endpoint and mouse euthanasia. In body weight a standard error of the mean (SEM) is shown. (**I**) JEV loads in mouse spleen and brain sampled at euthanasia due to severe clinical signs or at the end of the study—at 12 days after injection.

Enrichment of CpG dinucleotides in viral genomes is an emerging approach for live attenuated vaccines and oncolytics [26–29,39]. We generated CpG-enriched JEV (JEV-CpG) (**Table S1**, **Table S2**, **File S2**) using the same ISA strategy as for JEV-WT (**Fig 3**). The Fragment I sequence was subdivided into two sub-fragments, Fragment-I-A and Fragment-I-B, produced using *synthetic clonal genes* in medium copy number bacterial plasmids. Transfection of four overlapping ISA DNA fragments successfully rescued infectious JEV-CpG in all five replicates, with the virus inducing CPE in all transfected wells at passages 0 and 1 (**Fig 4D**). Sanger sequencing confirmed the correct sequences in the JEV-CpG stock. The JEV-CpG stock produced in VERO-ZAP-KO cells showed high infectious titers on both VERO-ZAP-WT and VERO-ZAP-KO cells (**Fig 4E**), along with high viral RNA loads (**Fig 4E**) and E protein expression (**Fig 4G**). Consistent with previous enrichment studies in Zika virus [27,28], the CpG-enriched JEV variant exhibited an attenuated phenotype in VERO-ZAP-WT cells (**Fig 4F**).

Next, we conducted a mouse study to identify whether JEV-WT and JEV-CpG obtained with the synthetic ISA approach causes typical infection phenotypes *in vivo*. The SA14-14-2 JEV strain does not cause infection in conventional mice; thus, we used *Ifnar1^-/-^* mice deficient for the type I IFN receptor function. The wild SA14-14-2 JEV strain causes clinical signs and lethality in *Ifnar1^-/-^* mice [40]. Five-week-old mice, six mice per group, were injected subcutaneously with virus-free media (MOCK) or ISA-derived JEV-WT or JEV-CpG. Mice in the MOCK group did not show clinical signs (**Fig 4H**); one MOCK mouse lost weight after manipulation and injection but recovered. Mice in the JEV-WT group showed a high 50% mortality due to paralysis and loss of 20% baseline body weight (**Fig 4H**). Another 50% of mice in the JEV-WT group also showed clinical signs—ataxia, depression and hunching but survived. Consistent with previous enrichment studies in Zika virus [27,28], the JEV-CpG variant showed an attenuated phenotype with no mortality or disease in mice (**Fig 4H**).

We tested spleen and brain from MOCK and JEV groups with the virus-specific RT-qPCR assay. As expected, JEV RNA was not detected in the MOCK group (**Fig 4I**). All samples from the JEV-WT group showed high viral RNA loads—10^5.58-7.52^ RNA copies per g of the brain, and 10^5.44-7.60^ RNA copies per g of the spleen (**Fig 4I**). In contrast, all brain samples from the JEV- CpG group were negative in RT-qPCR (**Fig 4I**). JEV-CpG RNA loads in the spleen were considerably lower (*P* < 0.0001, t-test) than JEV-WT loads (**Fig 4I**).

Infection phenotypes in our study with the ISA-derived JEV-WT were comparable to the infection phenotypes in the previous study where *Ifnar1^-/-^*mice injected with the wild (not synthetic) SA14-14-2 JEV strain also showed mortality, clinical signs, and virus in the spleen [40]. In contrast to our study however, the authors did not find wild SA14-14-2 JEV in the brain. The discrepancy can be explained by the different sampling time—at 12 days after injection in our study, and 4 days in the previous study [40]. Also, in the previous study, mice injected with 10^6.7^ plaque forming units (PFU) showed 100% mortality, while mice in our study (10^7^ TCID_50_/mouse) showed 50% mortality and 100% morbidity. The difference may be due to different methods to quantify viral loads for injection (PFU versus TCID_50_ assay), the route of injection—intraperitoneal previously and subcutaneous in our study, and potentially reduced viral population complexity in synthetically rescued JEV compared to wild viruses.

Altogether, dividing the toxic JEV fragment into two sub-fragments provides a highly efficient and robust ISA approach for reverse genetics. This method overcomes the bacterial toxicity caused by JEV sequences and enables the successful rescue of synthetic recombinant JEV.

### Infectious-Subgenomic-Amplicons methods to overcome bacterial toxicity caused by West Nile virus sequences

During the rescue of the wild-type WNV NY99 strain (WNV-WT), Fragment I exhibited bacterial toxicity when using a medium copy bacterial plasmid (**Fig 5**). To address this, a low copy plasmid was utilized (**Fig 5**), which enabled the successful rescue of Fragment I after several attempts over a three-month period.

**Figure 5.**
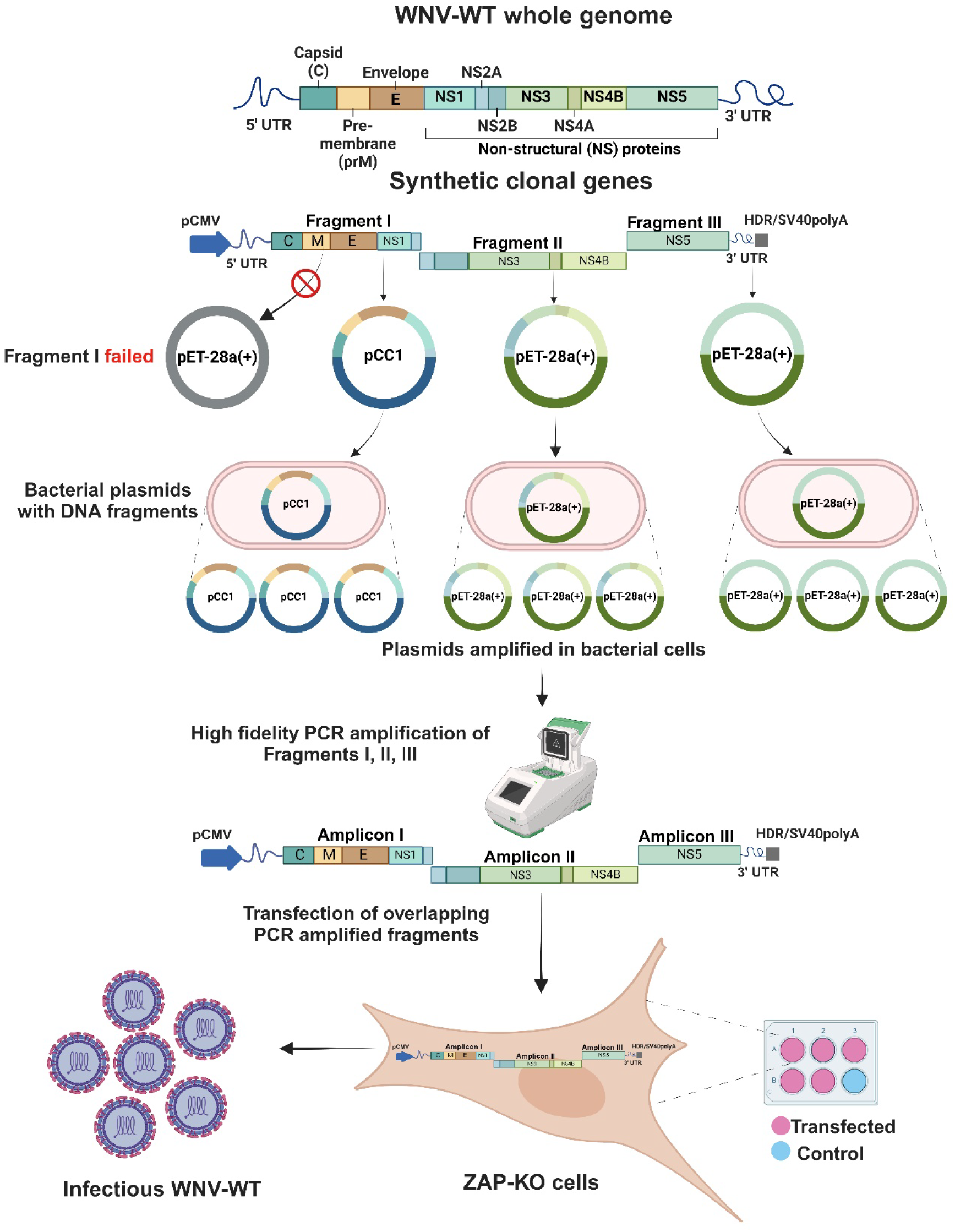
The adapted ISA method for the wild-type WNV variant (WNV-WT). To overcome toxicity, we generated Fragment I in the low copy pCC1 bacterial plasmid (**Table S1**, **File S1**).

As observed with JEV (**Fig 3**), classical ISA failed to rescue the WNV NY99 strain which was permutated (WNV-Per) to generate the permutated control virus (see materials and methods; **File S2**, **Table S2**) across three different medium and low copy bacterial plasmids (**Table S1, Fig 6**). To overcome this issue, the toxic Fragment I sequence (4,358 bp) was divided into three sub-fragments: Fragment-I-A (1,198 bp), Fragment-I-B (1,126 bp), and Fragment-I-c (2,179 bp) (**Table S1, File S1**). The first two fragments were made shorter to facilitate bacteria-free *synthesis of DNA* (available only for fragments shorter than 1.8 kb) based on the assumption that the 5’ flanking CMV promoter sequence contributed to the toxicity in bacteria. The longer Fragment-I-c was used for *synthetic clonal genes* with medium copy plasmid and bacterial amplification (**Table S1, File S1**). However, due to continued toxicity and failure, Fragment-I-c was further subdivided into overlapping Fragment-I-C (1,162 bp) and Fragment-I-D (1,089 bp), both small enough for bacteria-free *synthesis of DNA*. This troubleshooting led to the six overlapping ISA fragments for WNV-Per (**Fig 6, Table S1, File S1**).

**Figure 6.**
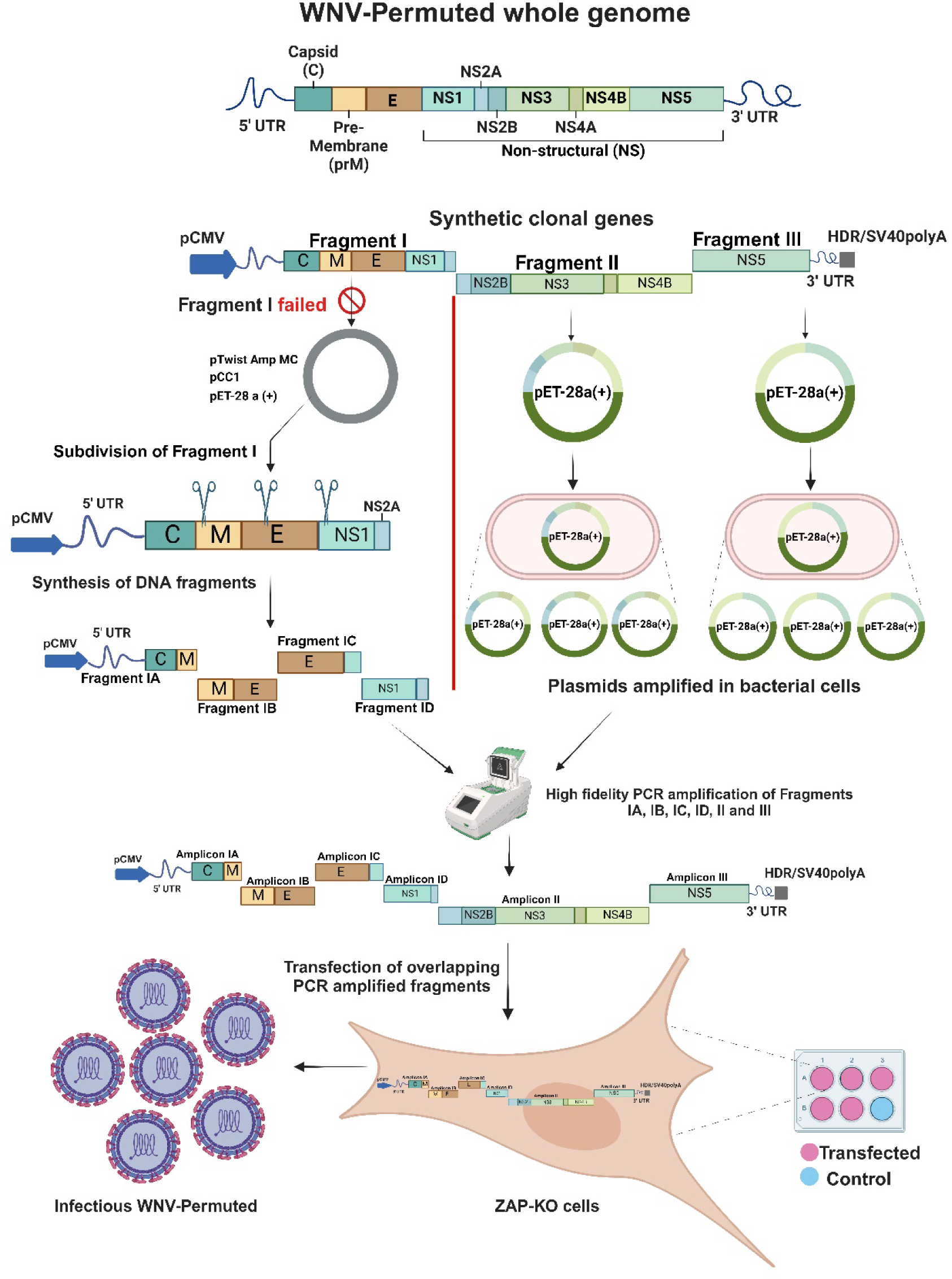
The adapted ISA method for the permuted WNV variant (WNV-Per). To overcome toxicity, we subdivide the toxic Fragment I into four sub-fragments for bacteria-free *synthesis of DNA* production (**Table S1**, **File S1**).

Transfection of the three overlapping ISA DNA fragments for WNV-WT and six overlapping ISA DNA fragments for WNV-Per consistently rescued the corresponding infectious WNV variants in all five replicates. Sanger sequencing confirmed correct sequences in WNV stocks. Viruses induced CPE in all transfected wells at passages 0 and 1 (**Fig 7A, B**), as well as in flasks during subsequent passaging and virus stock generation. Both WNV-WT and WNV-Per stocks produced in VERO-ZAP-KO cells had high infectious titers when titrated on both VERO-ZAP-WT and VERO-ZAP-KO cells (**Fig 7C, D**), along with high viral RNA loads (**Fig 7C, D**) and E protein expression (**Fig 7E, F**).

**Figure 7.**
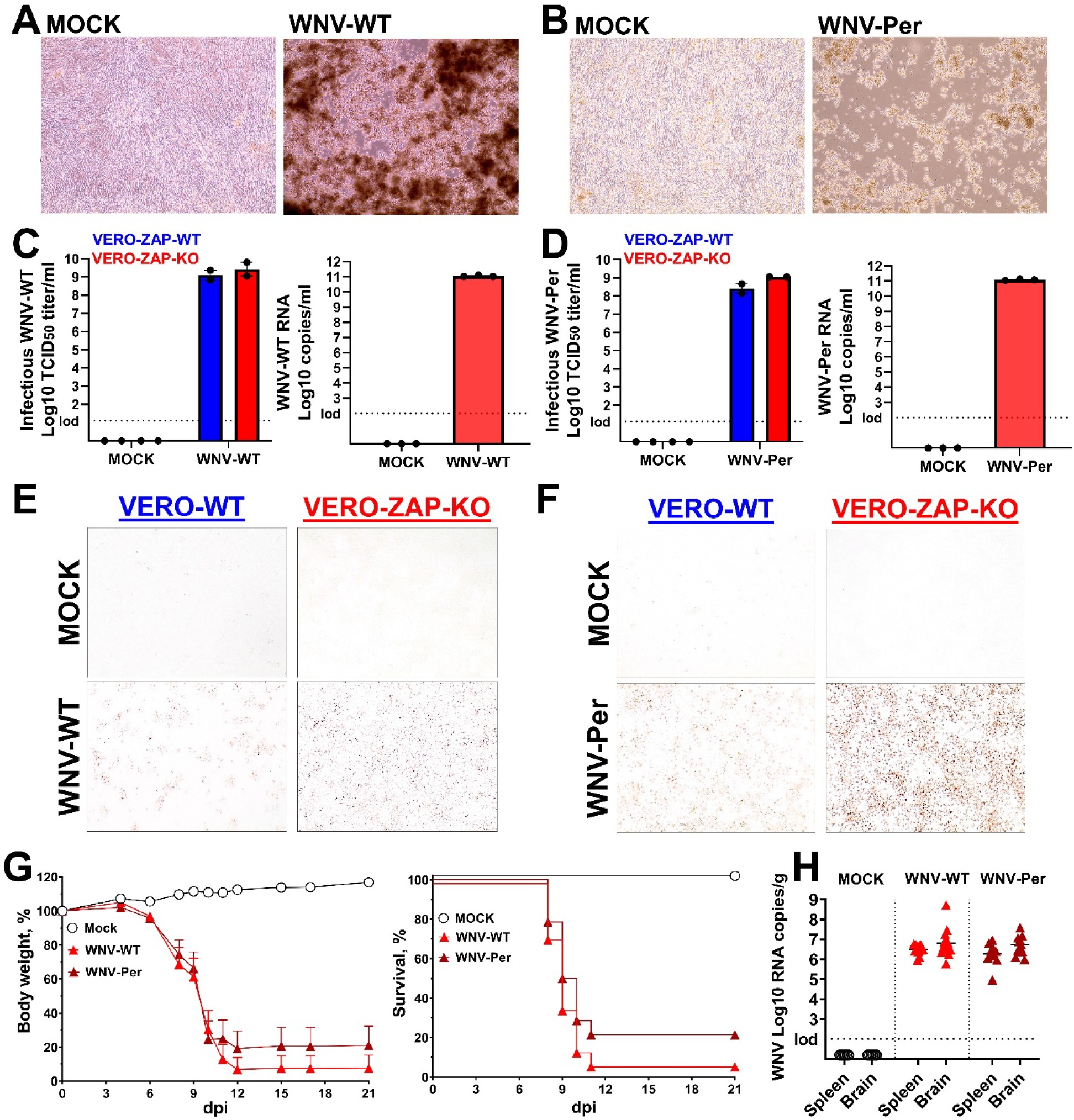
WNV ISA and stock generation. Cytopathic effect in WNV-WT (**A**) and WNV-Per (**B**) ISA-transfected ZAP-KO BHK+VERO cells, passage 0, day 6 after transfection. All five replicates for each WNV variant showed CPE; representative images are shown. Infectious TCID_50_ titers and RT-qPCR RNA loads of the WNV-WT (**C**) and WNV-Per (**D**) stocks produced in ZAP- KO cells. For TCID_50_ virus stocks were titrated in VERO-ZAP-WT or VERO-ZAP-KO cells. Images of cells positive for the WNV-WT (**E**) and WNV-Per (**F**) E protein (red staining) at 3 days after infection, MOI 0.01. Magnification x200. The experiment was done in 3 biological replicates. Figures represent the general patterns in all replicates. (**G**) Mean body weight and survival in MOCK C57BL/6J mice and C57BL/6J mice injected with the ISA-derived WNV-WT and WNV- Per. The following clinical scoring was used: 0—no visible abnormalities; 1—mild ataxia or tremors; 2—obvious ataxia or tremors; 3—depression, hunching, reluctance to walk, and falling to the side when walking; 4—close to moribund but still somewhat responsive; 5—paralysis; 6— loos of 20% baseline body weight; 7—found dead. Scores 4, 5, or 6 were used for the endpoint and mouse euthanasia. In body weight a standard error of the mean (SEM) is shown. (**H**) WNV loads in mouse spleen and brain sampled at euthanasia due to severe clinical signs or at the end of the study.

Next, we conducted a mouse study to identify whether WNV-WT and WNV-Per variants obtained with the synthetic ISA approach cause typical infection phenotypes *in vivo*. We used a well-established C57BL/6J mouse model for severe WNV infection [41–43]. Six-week-old C57BL/6J mice (14 mice per group) were MOCK infected or injected intraperitoneally (IP) with WNV-WT or WNV-Per variants. Mice in the MOCK group did not show clinical signs, weight loss, or mortality (**Fig 7G**). Mice in the WNV-WT and WNV-Per groups showed high mortality—93% and 79% (**Fig 7G**), ataxia, depression, paralysis, loss of 20% body weight, and death.

We tested spleen and brain from MOCK and WNV groups with the virus-specific RT-qPCR assay. As expected, WNV RNA was not detected in the MOCK group (**Fig 7H**). In contrast, all samples from the WNV-WT and WNV-Per groups showed high WNV RNA loads—10^4.98-6.97^ RNA copies per g of the spleen, and 10^5.78-8.72^ RNA copies per g of the brain (**Fig 7H**). The difference in viral RNA loads between WNV-WT and WNV-Per groups was not statistically significant (*P* < 0.05), which is expected because permutation in the WNV-Per variant does not affect encoded proteins and does not considerably affect genome coding parameters (**Table S2**).

Infection phenotypes in our study with the ISA-derived WNV variants were comparable to the infection phenotypes in the previous study where C57BL/6J mice injected with the wild (not synthetic) WNV NY99 strain also showed mortality, clinical signs, and virus in the spleen and brain [41–43].

### Comparative Infectious-Subgenomic-Amplicons in ZAP wild-type and ZAP knockout cells

The zinc finger antiviral protein 1 (ZAP) exhibits broad-spectrum antiviral activity. Recent studies, including our own, have demonstrated that flaviviruses such as Zika virus, JEV, and WNV are susceptible to ZAP antiviral effects in VERO cells [44–46]. Here, we compared the efficiency of JEV and WNV ISA in wild-type (VERO-ZAP-WT) and ZAP knockout (VERO-ZAP-KO) cells. We selected JEV-WT, JEV-CpG, and WNV-Per variants, using four and six ISA fragments (**Figs 3, 6**) for comparative ISA. Infectious-Subgenomic-Amplicons for each variant were performed concurrently, as described in the supplemental materials and methods, with DNA amplicons for transfection into VERO-ZAP-WT or VERO-ZAP-KO cells derived from the same PCR reactions.

JEV-WT induced CPE in all five replicates of both ZAP-WT and ZAP-KO cells during passage 0, with more pronounced CPE in ZAP-KO cells (**Fig 8A**). JEV-CpG also induced CPE in all five replicates of both ZAP-WT and ZAP-KO cells during passage 0, with more pronounced CPE in ZAP-KO cells (**Fig 8A**). WNV-Per caused equally strong CPE in both ZAP-WT and ZAP-KO cells (**Fig 8A**). During passage 1, JEV-WT, JEV-CpG, and WNV-Per infections in ZAP-KO cells were associated with higher E protein expression compared to ZAP-WT cells (\*\*\**P* < 0.0001; **Figs 8B, D, F**). Consequently, JEV-CpG and WNV-Per TCID_50_ infectious titers in supernatants were significantly higher in ZAP-KO cells than in ZAP-WT cells (*P* = 0.014-0.017) (**Figs 8C, E, J**).

**Figure 8.**
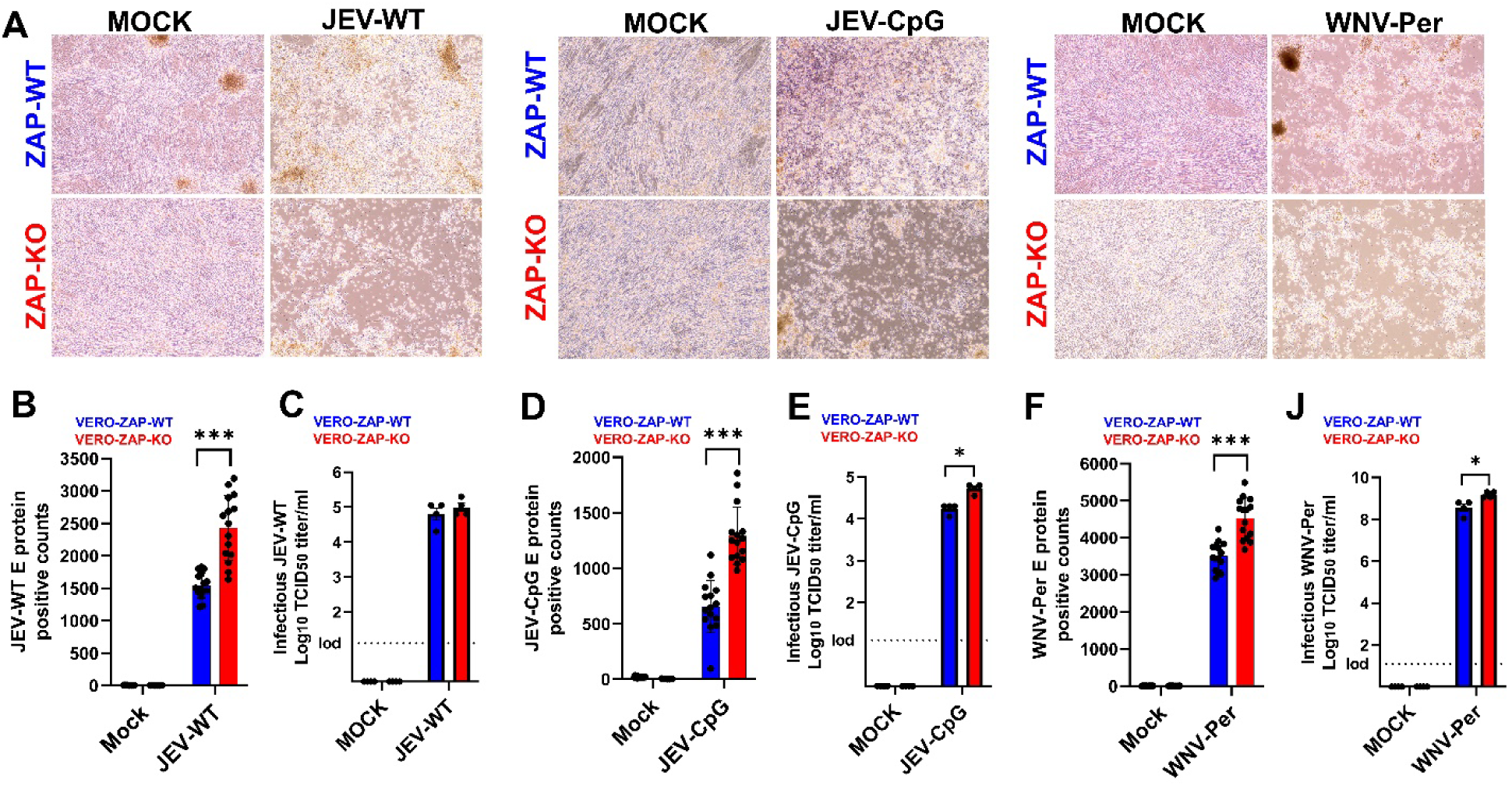
Comparative ISA in ZAP-WT and ZAP-KO cells. (**A**) Cytopathic effect in JEV-WT, JEV-CpG, and WNV-Per ISA-transfected ZAP-WT and ZAP-KO BHK+VERO cells, passage 0, day 5 (JEV-WT and JEV-CpG) and day 4 (WNV-Per) after transfection. All five replicates for JEV-WT, JEV-CpG, and WNV-Per showed CPE; the representative images are shown. The digital quantification of JEV-WT (**B**), JEV-CpG (**D**), and WNV-Per (**F**) E protein-positive cells in all 5- well replicates at day 4 (JEV) and day 2 (WNV) during passage 1 of comparative ISA. For technical replicates, three random images were obtained from each well. Examples of JEV-WT, JEV-CpG, and WNV-Per E protein IHC are in Figures 4C**, G,** and **7F**. **\*\*\****P*<0.0001, unpaired t- test. JEV-WT (**C**), JEV-CpG (**E**), and WNV-Per (**J**) infectious TCID_50_ titers in supernatants collected at day 4 (JEV) and day 2 (WNV-Per) during passage 1 of comparative ISA. Data from 4-well replicates are shown. **\****P*<0.014-0.017, unpaired t-test.

Overall, both wild-type and ZAP knockout cells demonstrated 100% ISA efficiency, successfully rescuing JEV and WNV in all five replicates. However, ISA in ZAP-KO cells resulted in the considerably higher number of virus-positive cells and elevated infectious virus loads.

## DISCUSSION

The goal of this study was to validate modified ISA strategies for producing toxic ISA flavivirus DNA fragments synthetically, overcoming bacterial toxicity and eliminating the need to amplify toxic fragments from biological specimens or other preexisting sources. These ISA strategies enable the generation of entirely synthetic JEV and WNV stocks. The strategies involved: (i) Subdividing the problematic toxic DNA fragment into two sub-fragments for subsequent *synthetic clonal gene* production with bacterial plasmids and bacteria. (ii) Using a low copy bacterial plasmid. (iii) Subdividing the toxic DNA fragment into four short overlapping sub-fragments, each up to 1.8 kb, for subsequent bacteria-free *synthesis of DNA* production. To our knowledge, the latter strategy had never been applied for ISA.

As expected, and previously demonstrated for the DENV 2 serotype [19], the classical ISA approach of subdividing the flavivirus genome into three overlapping fragments of approximately 4 kb each works well for DENV. These fragments were synthesized quickly and without technical difficulties by a commercial supplier using a medium copy bacterial plasmid (**Table S1**).

In contrast, this classical three-fragment ISA approach failed when we attempted to rescue JEV- WT Fragment I using three different medium and low copy bacterial plasmids (**Table S1**). The solution was to divide the toxic fragment into two sub-fragments. We encountered similar technical difficulties producing WNV-WT Fragment I. Here, replacing the medium copy plasmid with a low copy plasmid resolved the issue, allowing the rescue of the entirely synthetic WNV- WT variant, but this took several attempts and over three months.

More challenging difficulties arose when we attempted to rescue the permuted WNV-Per variant. Similar to JEV ISA, we failed to produce Fragment I using three different medium and low copy bacterial plasmids. Subdividing the toxic Fragment I into three sub-fragments (A, B, and c) did not work because the sub-fragment Fragment I-c was also toxic to bacteria (**Table S1**).

Subsequently, we divided the highly toxic WNV-Per Fragment I into four sub-fragments, each smaller than 1.8 kb. This allowed the *synthesis of DNA* commercial production process that is bacteria-free (**Table S1**).

While various approaches have been employed to overcome technical difficulties in constructing cDNA molecules representing the full-length flavivirus genomes in bacterial vectors, this process remains challenging, laborious, and time-consuming. As demonstrated by our difficulties during the rescue of synthetic JEV and WNV (**Table S1**), these challenges are also present in the classical ISA approach. We propose that subdividing toxic fragments into sub-fragments smaller than 1.8 kb suitable for the *synthesis of DNA* process is currently the most rapid and robust strategy to overcome the need for bacterial plasmids and bacteria and ensure the fast rescue of entirely synthetic flaviviruses.

This strategy significantly shortens the time required to produce recombinant flaviviruses. For example, the *synthesis of DNA* process by a commercial supplier (**Table S1**) has turnaround time 2-4 days. Additionally, the synthesis of DNA is cost-effective with current 0.07 USD per bp; for example, 4.3 kb DNA fragment for WNV ISA ordered via *synthetic clonal genes* process with bacterial plasmids and amplification in bacteria costs around 840 USD with 3-4 weeks turnaround time, while the same fragment divided into four sub-fragments and ordered via bacteria-free *synthesis of DNA* costs 360 USD with 2-4 days turnaround time. During production, the provider flanks DNA fragments with short 22 bp adapters (**File S1**), facilitating efficient PCR amplification of ordered ISA fragments from the provided construct. The provider also applies NGS to verify correct sequences, reducing both time and costs for virus rescue. We anticipate that ongoing improvements in DNA synthesis technology and associated cost reductions will provide even faster and more cost-effective bacteria-free fragments for ISA, enabling the even more rapid production of entirely synthetic flaviviruses.

The zinc finger CCCH-type antiviral protein 1, also known as ZAP, ZC3HAV1, or PARP13, is a cellular protein with broad antiviral activity. ZAP binds viral RNA and evokes antiviral activity by mediating viral RNA degradation and translational inhibition [47]. It also interacts with various cellular proteins that may act as co-factors, enhancing its antiviral effects [48].

Furthermore, ZAP can augment other antiviral systems [44,49]. Recent studies, including our own, have demonstrated that flaviviruses such as Zika virus, JEV, and WNV are susceptible to ZAP antiviral effects in VERO cells [44–46]. Thus, we also compared the efficacy of JEV-WT.

JEV-CpG, and WNV-Per ISA in ZAP-WT and ZAP-KO cells. Both cell types showed 100% efficiency, with viruses rescued in all five replicates. However, the increased number of virus-positive cells and infectious titers in ZAP-KO cells suggests that these cells may be more effective for rescuing modified flaviviruses, such as attenuated variants for live attenuated vaccines, including CpG-enriched vaccines. This is significant because CpG dinucleotide enrichment is a novel vaccine approach. Increasing CpG dinucleotides in viral RNA, while preserving the original amino acid composition, may weaken the virus, preventing disease but potentially inducing strong antiviral immunity. However, CpG-enriched vaccine candidates are attenuated both *in vitro* and *in vivo*, which can complicate their rescue and the generation of high-titer vaccine stocks [27,28]. Using VERO-ZAP-KO cells could aid in developing CpG-enriched vaccine candidates, and it is advantageous that VERO cells are already used as a GMP line for producing many vaccines.

In conclusion, the ISA method is a robust and straightforward approach to reverse genetics for JEV and WNV. In this study, we validated a modified ISA strategy that involves subdividing the toxic DNA fragment into four short overlapping sub-fragments, each up to 1.8 kb, for subsequent bacteria-free *DNA synthesis* production. To our knowledge, this strategy had never been applied to ISA before, enabling the rapid generation of flaviviruses using a completely synthetic method for toxic ISA flavivirus fragments. The strategy can be particularly beneficial for research on emerging flaviviruses where a rapid response is needed to rescue new virus variants during outbreaks.

## ETHICS APPROVAL STATEMENT

The animal studies were approved by The Ohio State University Institutional Biosafety Committee (e-Protocol # 2022R00000111) and The Institutional Animal Care and Use Committee (e-Protocol #2023A00000095).

## AUTHOR CONTRIBUTIONS

Conceptualization: PPS, NPKL, UK. Investigation: PPS, NPKL, UK. Data analysis: PPS, NPKL, UK. Funding: UK. Writing—original draft preparation: PPS, NPKL, UK. Writing—review and editing: PPS, NPKL, UK.

## DATA AVAILABILITY

All data are available in the manuscript or in the supplemental data.

## SUPPORTING INFORMATION CAPTIONS

### Supplemental materials and methods

**Table S1** ISA fragments and primers.

**Table S2** JEV and WNV genomic parameters.

**File S1** ISA fragments.

**File S2** JEV-CpG and WNV-Permuted reference sequences.

## SUPPLEMENTAL MATERIALS AND METHODS

### Cells

We maintained wild-type VERO E6 (VERO-ZAP-WT) and BHK-21 (BHK-ZAP-WT) cells, along with CCCH-type zinc finger antiviral protein (ZAP) knockout (ZAP-KO) derivatives (VERO-ZAP-KO and BHK-ZAP-KO), in DMEM (Fisher, MA, USA; #11-965-118) supplemented with 10% heat-inactivated fetal bovine serum (FBS, Fisher, MA, USA; #A5256801), 1x Penicillin-Streptomycin (Fisher, MA, USA; #15140122), and 2.67 mM sodium bicarbonate (Fisher, MA, USA; # 25080094) at +37°C in a 5% CO_2_ humidified incubator.

To generate the VERO-ZAP-KO cell line, we previously used the guide RNA (gRNA) GTCTCTGGCAGTACTTGCGA targeting the first exon of the *Chlorocebus sabaeus* (Gene ID: 103226990) ZC3HAV1 gene, necessary for expressing all ZAP isoforms. A non-targeting gRNA (ACGGAGGCTAAGCGTCGCAA) served as the control [1,2]. These gRNAs were transiently transfected into cells using GenCrispr NLS-Cas9-NLS Nuclease (GenScript). Transfected cells were then seeded in 96-well plates through limiting dilution to generate isogenic single clones.

After expansion, the clones were genotyped via Sanger sequencing using primers ATCGCTGGGCTGGACTAACG and GCAGAGAAGGGAGTGGCTGAA to identify location of indels on *Chlorocebus sabaeus* genome: -2 bp deletion, NW_023666072.1: 4677550- 4677551. The knockout subclone confirmed by genotyping was further validated by western blot for ZAP expression in our previous studies [1,2]. Negative control cells were not isolated into single clones but were validated by Sanger sequencing to confirm the absence of indels.

The same strategy was used to generate BHK-ZAP-KO cells with gRNA GGAGATCGTAGGCCAGACCGGGG targeting the first exon of the *Mesocricetus auratus* (Gene ID: 110343172) Zc3hav1 gene and the same non-targeting control gRNA. After expansion, the clones were genotyped via Sanger sequencing with primers GCTGCTTCATCACCAAGATCC and GGTCGGGAAGACCCTTTCAC to identify location of indels on *Mesocricetus auratus* genome: -29 bp deletion, NW_024429200.1: 3564916-3564944. We confirmed the absence of mycoplasma contamination in all cells using the LookOut Mycoplasma PCR Detection Kit (Millipore Sigma, MA, USA; #MP0035).

### *In silico* CpG enrichment and permutation in viral genomes

For CpG dinucleotide enrichment in the JEV genome, specifically in the region encoding the E protein (**File S2**), we used the SSE 1.3 software package [3], as we previously described [4,5]. The genomic parameters of the wild-type and CpG-enriched JEV variants are listed in **Table S2**.

Certain experiments involving viral genomic mutations need permuted or scrambled virus control variants. Permutation can potentially impact the efficiency of reverse genetics methods. To simulate this scenario for ISA, we generated a permuted control for WNV. For permutation of WNV sequences we selected genomic regions encoding E and NS1 (**File S2**) because these proteins are important for flavivirus pathogenesis, modulation of the infection cycle, viral RNA replication, and host immune evasion. Moreover, we previously permuted these genomic regions in the Zika virus genome; this permutation did not affect Zika infection phenotypes *in vitro* and in mice [4]. We used an SSE 1.3 software package [3], specifically the CDLR randomizing in the SSE 1.3 [3,6–8]. During randomization we sought to avoid areas of the genome containing RNA elements that are required for replication or translation of the virus genome, such as cis- replicating elements, gene start or gene-end signals; we also avoided regions with prominent secondary structures. The CDLR randomizes the order of codons within the sequence while maintaining encoded proteins, dinucleotide frequencies, and coding parameters (**Table S2**).

### Infectious-Subgenomic-Amplicons (ISA) and virus stock generation

The following reference sequences were used for *de novo* synthesis of flaviviruses: DENV type 2 strain 16681 [GenBank: U87411.1], JEV strain SA14-14-2 [GenBank: MK585066.1], and WNV strain NY99 [GenBank: DQ211652.1]. To rescue DENV, JEV, and WNV, initially, we used three overlapping (70-80 nt overlap) ISA DNA fragments (3,371-4,550 bp length) covering the entire viral genome (**Table S1**; **File S1**); three overlapping DNA fragments is common ISA strategy for flaviviruses [9,10]; thus, we consider it as a classical approach. The first and last fragments for each virus are flanked with pCMV promoter and HDR/SV40pA sequences (**File S1**).

Initially all DNA fragments for ISA were ordered through GeneScript (NJ, USA) or Twist Biosciences (CA, USA) *synthetic clonal genes* services where DNA fragments are synthesized, inserted into the bacterial plasmids (**Table S1**), amplified in bacteria, extracted, sequenced, and plasmids with inserted fragments are delivered as lyophilized DNA with known concentration. Some flavivirus ISA DNA fragments were toxic to bacteria, in both medium copy and low copy plasmids (**Table S1**), and after multiple attempts *synthetic clonal genes* approach failed (see results for details). To overcome this flavivirus DNA sequence toxicity to bacteria, we split toxic clonal genes into smaller overlapping fragments below 1.8 kb length. These smaller size fragments allowed *synthesis of DNA* services from Twist Biosciences were DNA fragments, flanked with 22 bp adapters (**File S1**), are rapidly (2-4 days service) synthesized without using bacterial plasmids and bacteria. Twist Biosciences and GeneScript confirmed the correct sequences of DNA fragments by next-generation-sequencing (NGS).

Overlapping clonal genes or synthetic DNA ISA fragments were amplified with high fidelity Invitrogen Platinum PCR SuperMix High Fidelity (Fisher, MA, USA; #12-532-016) (primers are in **Table S1**), mixed in equimolar concentration to obtain the final 1µg of DNA for transfection, and transfected into BHK-ZAP-KO cell monolayers (6-well plates, five transfection replicates, one mock-transfection control well) with Lipofectamine 3000 (Fisher, MA, USA; #L3000015) in 640 µl of OptiMEM (Fisher, MA, USA; #11-058-021) for 12 h at +37°C, 5% CO_2_. Afterward, OptiMEM media was removed and replaced with 3 ml of DMEM with 10% of FBS containing 4x10^5^ VERO-ZAP-KO cells on the top of BHK-ZAP-KO cells and incubated plates for additional 6-7 days (passage 0). For passage 1, 400 µl of media from each well of passage 0 was transferred to wells with VERO-ZAP-KO cells and incubated for 4-7 days. During passages 0 and 1, cells were monitored for cytopathic effect (CPE). Generated flaviviruses were additionally passaged 2-3 times on VERO-ZAP-KO cells to generate virus stocks; flasks were monitored for CPE. Cell culture media from the final stocks were centrifuged (12,000 g, 20 min, +4°C), aliquoted, and frozen (-80°C). Viral titers were quantified in duplicate in VERO-ZAP-WT and VERO-ZAP-KO cells with endpoint dilution assay described below. The absence of mycoplasma contamination in all virus stocks and cell cultures was confirmed using PCR Detection Kit (Millipore Sigma, MA, USA; #MP0035).

To validate virus stock sequences, RNA was extracted from virus stock culture supernatants as described below. PCR was performed using SuperScript™ IV One-Step RT-PCR System (Fisher, MA, USA; #12-594-025) as per manufacturer’s instructions with the primers targeting sequences encoding E protein: DENV (Forward 5- GACAGCTGTCACTCCTTCAAT GAC-3 and Reverse 5- CATGCTCTTCCCCTGAGTGAG G-3); JEV-WT (Forward 5- GTTTTAATTGTCTGGGAATGGGC-3 and Reverse 5-GCCAACTTTGTATTTGATGTTTTC-3); JEV-CpG (Forward 5-CAGTTTTAATTGTCTGGGAATGGG-3 and Reverse 5- GTCAGCATGCACATTGGTCG-3); WNV-WT (Forward 5- CTATTGCTTTTGGTGGCCCCAG-3 and Reverse 5-GTAGTTGGTCCATGGACAAA-3); WNV-Per (Forward 5-CTATTGCTTTTGGTGGCCCCAG-3 and Reverse 5- GTAGTTGGTCCATGGACAAA-3). PCR products were run on a 1% agarose gel, and DNA was purified and concentrated using DNA Clean & Concentrator-5 (Zymo Research, CA, USA; # D4013) as per manufacturer’s instructions. DNA was sequenced by Sanger sequencing and sequences were aligned with corresponding ISA sequences (**File S1**). Sanger sequencing confirmed correct sequences in DENV, JEV, and WNV stocks.

### Comparative ISA

To compare ISA efficiency on ZAP-WT and ZAP-KO cells, JEV and WNV fragments were transfected into ZAP-WT or ZAP-KO cells as above. These experiments were conducted simultaneously; also, the DNA amplicons for transfection in ZAP-WT or ZAP-KO cells were obtained from the same PCR reactions. During passage 0, ZAP-WT and ZAP-KO cells were observed for 5 (JEV variants) and 4 (WNV variants) days to compare CPE; the time was shorter for WNV because it causes more aggressive infection and CPE. During the 1st passage, CPE was observed, supernatants were collected on days 2-4 (2 for WNV variants; 4 for JEV variants) from each well, and infectious virus titers were quantified with the end-point dilution assay as described below. The shorter passage duration in the comparative ISA than in the initial ISA used for virus rescue were applied to preserve cell monolayers for quantitative immunohistochemistry (IHC) described below. The quantitative difference between ZAP-WT and ZAP-KO conditions was represented by TCID_50_ infectious titers in cell culture supernatants from four replicates. Additionally, cellular monolayers from each transfected and mock-transfected well (five replicates) were fixed and stained for viral E protein via IHC as described in the infectious virus titration below. To quantify and compare IHC staining between ZAP-WT and ZAP-KO conditions, three images (magnification x200) were randomly acquired in all five well replicates with the same microscopic settings. Afterward, each image was processed in ImageJ (NIH) (https://imagej.net/imaging/particle-analysis) with Automatic Particle counting [2].

### Mouse experiments

The animal studies were approved by The Ohio State University Institutional Biosafety Committee (e-Protocol # 2022R00000111) and IACUC (e-Protocol #2023A00000095). Mice were euthanized by deeply anesthetizing them with isoflurane, followed by cervical dislocation.

We conducted mouse experiments to identify whether ISA-rescued synthetic JEV and WNV cause expected infection phenotypes which were previously described in mice infected with wild or field isolates. The wild SA14-14-2 JEV strain does not cause clinical signs in conventional mice; thus, we used *Ifnar1^-/-^*mice (The Jackson Laboratory, USA; Strain #:028288). The SA14- 14-2 JEV strain causes clinical signs or lethality in infected *Ifnar1^-/-^* mice [11]. After one week of acclimatization, five-week-old mice (6 mice per group) were injected subcutaneously with the virus-free media (MOCK) or ISA-derived JEV-WT or JEV-CpG variants (**Table S1**). We standardized mouse injection doses by normalization of viral RNA loads (determined by RT-qPCR, see below) in the JEV-WT and JEV-CpG inoculums which resulted in 10^10^ RNA copies/mouse for both variants (JEV-WT – 10^7^ TCID_50_/mouse; JEV-CpG – 10^6.3^ TCID_50_/mouse; titers quantified on VERO-ZAP-WT cells, see below). Afterward, mice were monitored for clinical signs and body weight. Mice were sacrificed when exhibiting greater than 20% weight loss or severe clinical signs. We sampled spleen and brain from deceased mice, or from survived mice at the end of the study to quantify JEV loads.

The wild NY99 WNV strain causes severe infection in conventional C57BL/6J mice; thus, we used the well-characterized C57BL/6J mouse lethal model [12–14]. Five-week-old C57BL/6J mice (Strain #:000664; 14 mice per group) were ordered from The Jackson Laboratory, acclimatized for a week, and injected intraperitoneally (IP) with the virus-free media (MOCK), ISA-derived WNV wild-type (WNV-WT), or ISA-derived WNV permuted (WNV-Per) variant (**Table S1**). We standardized mouse injection doses by normalization of viral RNA loads (determined by RT-qPCR, see below) in the WNV-WT and WNV-Per inoculums which resulted in 10^7^ RNA copies/mouse for both variants (WNV-WT - 10^5^ TCID_50_/mouse; WNV-Permuted -10^5.6^ TCID_50_/mouse; titers quantified on VERO-ZAP-WT cells, see below). Afterward, mice were monitored for clinical signs and body weight changes. Mice were sacrificed when exhibiting 20% or greater weight loss or severe clinical signs, or at the end of the study to sample spleen and brain from mice.

### Infectious virus titration

To quantify infectious loads in DENV, JEV and WNV stocks, the endpoint dilution assay was used to determine the 50% tissue culture infectious dose (TCID_50_), with two replicates, as we had described before [4,15–19]. Viral stocks were serially diluted in four replicates in media and 50 μl of each dilution was added to confluent VERO-ZAP-WT or VERO-ZAP-KO cells cultured in 96- well plates. Dilutions were made in DMEM media with 1% FBS. After 2 h at 37°C, 150 μl of media was added. The cells were incubated for 5-7 days (5 days for WNV; 7 days for DENV and JEV) at 37°C, 5% CO_2_. Afterward, plates were washed, dried, formalin fixed, and stained with specific antibodies (Ab) [4,15–19]. For DENV and WNV IHC, we used primary anti-pan flavivirus envelop (E) protein monoclonal Ab D1-4G2-4-15 (ATCC, VA, USA; #HB-112) followed by Goat anti-Mouse IgG HRP Ab (Abcam, MA, USA; #ab97023). For JEV, we used JEV E polyclonal Ab (GeneTex, CA, USA; #GTX125867;) followed by Goat Anti-Rabbit IgG H&L (HRP) Ab (Abcam, MA, USA; #ab97051) [4,15–19]. Infectious titers were calculated by the Spearman-Kärber formula and expressed in log_10_ TCID_50_ per ml. Media from mock-inoculated cells were used as negative controls.

### RNA extraction and virus-specific reverse transcriptase quantitative polymerase chain reaction assays (RT-qPCR)

The RNA from cell culture supernatants was extracted with QIAamp Viral RNA Mini Kit (QIAGEN) according to the manufacturer’s instructions. The RNA from mouse tissues was extracted with PureLink RNA Mini Kit (Invitrogen) according to the manufacturer’s instructions. All RT-qPCR reactions were conducted on the QuantStudio 3 real-time PCR system (Applied Biosystems) and analyzed using QuantStudio Design & Analysis Software v1.5.2.

For DENV (F: 5′-AAGGACTAGAGGTTAKAGGAGACCC-3′; R: 5′- GGCGYTCTGTGCCTGGAWTGATG-3′; Probe: 56-JOEN/AA CAG CAT A/ZEN/T TGA CGC TGG GAA AGA CC-3IABkFQ), JEV-WT and JEV-CpG (F: 5′-GCCACCCAGGAGGTCCTT- 3′; R: 5′-CCCCAAAACCGCAGGAAT-3′; Probe: 56-FAM-CAAGAGGTG/ZEN/GACGGCC-3IABkFQ), and WNV-WT and WNV-Per (F: 5′-AGTAGTTCGCCTGTGTGAGC-3′; R: 5′- GCCCTCCTGGTTTCTTAGA-3′; Probe: FAM-AATCCTCACAAACACTACTAAGTTTGTCA-TAMRA) we used the previously validated probe-based one-step RT-qPCR assays [20–22]. The Luna Universal Probe One-Step RT-qPCR Kit (NEB) reaction mixture (20 μL) consisted of 10 μL Luna Universal Probe One-Step Reaction Mix, 1 μL Luna WarmStart RT Enzyme Mix, 1 μL (10 μM) of forward and reverse primers, 0.5 μL of the probe, 2.5 μL nuclease-free water and 4 μL of RNA template. The reverse transcription and enzyme activation steps of 10 min at 55°C and 1 min at 95°C were followed by 40 amplification cycles (10 s at 95°C and 30 s at 60°C). A standard curve was used to quantify viral RNA loads as we described [15].

PCR values were corrected for fluid volumes or tissue weights and upon logarithmical transformation expressed as virus RNA genome copies per ml or g. Strict precautions were taken to prevent PCR contamination. All master mix preparations were done in the dedicated PCR cabinet. Aerosol-resistant filter pipette tips and disposable gloves were used. Kit reagent controls were included in every RNA extraction and PCR run.

### Statistics

We used GraphPad 9 PRISM for statistics. For the comparison of the viral RNA loads, infectious titers, and quantitative IHC we used an unpaired t-test. *P*-value < 0.05 was considered statistically significant.

