## supplemental tables and files for "Infectious subgenomic amplicon strategies for Japanese encephalitis and West Nile viruses": File S1 ISA fragments.docx

**DENV wild-type ISA Fragments**

DENV type 2 strain 16681 [GenBank: U87411.1].

Highlighted in red – pCMV promoter sequence.

Highlighted in green – HDR/SV40pA sequence.

>DENV2_16681_**Fragment-I**

**CACCCAACTGATCTTCAGCATCTTCAATATTGGCCATTAGCCATATTATTCATTGGTTATATAGCATAAATCAATATTGGCTATTGGCCATTGCATACGTTGTATCTATATCATAATATGTACATTTATATTGGCTCATGTCCAATATGACCGCCATGTTGGCATTGATTATTGACTAGTTATTAATAGTAATCAATTACGGGGTCATTAGTTCATAGCCCATATATGGAGTTCCGCGTTACATAACTTACGGTAAATGGCCCGCCTGGCTGACCGCCCAACGACCCCCGCCCATTGACGTCAATAATGACGTATGTTCCCATAGTAACGCCAATAGGGACTTTCCATTGACGTCAATGGGTGGAGTATTTACGGTAAACTGCCCACTTGGCAGTACATCAAGTGTATCATATGCCAAGTCCGCCCCCTATTGACGTCAATGACGGTAAATGGCCCGCCTGGCATTATGCCCAGTACATGACCTTACGGGACTTTCCTACTTGGCAGTACATCTACGTATTAGTCATCGCTATTACCATGGTGATGCGGTTTTGGCAGTACACCAATGGGCGTGGATAGCGGTTTGACTCACGGGGATTTCCAAGTCTCCACCCCATTGACGTCAATGGGAGTTTGTTTTGGCACCAAAATCAACGGGACTTTCCAAAATGTCGTAATAACCCCGCCCCGTTGACGCAAATGGGCGGTAGGCGTGTACGGTGGGAGGTCTATATAAGCAGAGCTCGTTTAGTGAACCG**AGTTGTTAGTCTACGTGGACCGACAAAGACAGATTCTTTGAGGGAGCTAAGCTCAACGTAGTTCTAACAGTTTTTTAATTAGAGAGCAGATCTCTGATGAATAACCAACGGAAAAAGGCGAAAAACACGCCTTTCAATATGCTGAAACGCGAGAGAAACCGCGTGTCGACTGTGCAACAGCTGACAAAGAGATTCTCACTTGGAATGCTGCAGGGACGAGGACCATTAAAACTGTTCATGGCCCTGGTGGCGTTCCTTCGTTTCCTAACAATCCCACCAACAGCAGGGATATTGAAGAGATGGGGAACAATTAAAAAATCAAAAGCTATTAATGTTTTGAGAGGGTTCAGGAAAGAGATTGGAAGGATGCTGAACATCTTGAATAGGAGACGCAGATCTGCAGGCATGATCATTATGCTGATTCCAACAGTGATGGCGTTCCATTTAACCACACGTAACGGAGAACCACACATGATCGTCAGCAGACAAGAGAAAGGGAAAAGTCTTCTGTTTAAAACAGAGGATGGCGTGAACATGTGTACCCTCATGGCCATGGACCTTGGTGAATTGTGTGAAGACACAATCACGTACAAGTGTCCCCTTCTCAGGCAGAATGAGCCAGAAGACATAGACTGTTGGTGCAACTCTACGTCCACGTGGGTAACTTATGGGACGTGTACCACCATGGGAGAACATAGAAGAGAAAAAAGATCAGTGGCACTCGTTCCACATGTGGGAATGGGACTGGAGACACGAACTGAAACATGGATGTCATCAGAAGGGGCCTGGAAACATGTCCAGAGAATTGAAACTTGGATCTTGAGACATCCAGGCTTCACCATGATGGCAGCAATCCTGGCATACACCATAGGAACGACACATTTCCAAAGAGCCCTGATTTTCATCTTACTGACAGCTGTCACTCCTTCAATGACAATGCGTTGCATAGGAATGTCAAATAGAGACTTTGTGGAAGGGGTTTCAGGAGGAAGCTGGGTTGACATAGTCTTAGAACATGGAAGCTGTGTGACGACGATGGCAAAAAACAAACCAACATTGGATTTTGAACTGATAAAAACAGAAGCCAAACAGCCTGCCACCCTAAGGAAGTACTGTATAGAGGCAAAGCTAACCAACACAACAACAGAATCTCGCTGCCCAACACAAGGGGAACCCAGCCTAAATGAAGAGCAGGACAAAAGGTTCGTCTGCAAACACTCCATGGTAGACAGAGGATGGGGAAATGGATGTGGACTATTTGGAAAGGGAGGCATTGTGACCTGTGCTATGTTCAGATGCAAAAAGAACATGGAAGGAAAAGTTGTGCAACCAGAAAACTTGGAATACACCATTGTGATAACACCTCACTCAGGGGAAGAGCATGCAGTCGGAAATGACACAGGAAAACATGGCAAGGAAATCAAAATAACACCACAGAGTTCCATCACAGAAGCAGAATTGACAGGTTATGGCACTGTCACAATGGAGTGCTCTCCAAGAACGGGCCTCGACTTCAATGAGATGGTGTTGCTGCAGATGGAAAATAAAGCTTGGCTGGTGCACAGGCAATGGTTCCTAGACCTGCCGTTACCATGGTTGCCCGGAGCGGACACACAAGGGTCAAATTGGATACAGAAAGAGACATTGGTCACTTTCAAAAATCCCCATGCGAAGAAACAGGATGTTGTTGTTTTAGGATCCCAAGAAGGGGCCATGCACACAGCACTTACAGGGGCCACAGAAATCCAAATGTCATCAGGAAACTTACTCTTCACAGGACATCTCAAGTGCAGGCTGAGAATGGACAAGCTACAGCTCAAAGGAATGTCATACTCTATGTGCACAGGAAAGTTTAAAGTTGTGAAGGAAATAGCAGAAACACAACATGGAACAATAGTTATCAGAGTGCAATATGAAGGGGACGGCTCTCCATGCAAGATCCCTTTTGAGATAATGGATTTGGAAAAAAGACATGTCTTAGGTCGCCTGATTACAGTCAACCCAATTGTGACAGAAAAAGATAGCCCAGTCAACATAGAAGCAGAACCTCCATTCGGAGACAGCTACATCATCATAGGAGTAGAGCCGGGACAACTGAAGCTCAACTGGTTTAAGAAAGGAAGTTCTATCGGCCAAATGTTTGAGACAACAATGAGGGGGGCGAAGAGAATGGCCATTTTAGGTGACACAGCCTGGGATTTTGGATCCTTGGGAGGAGTGTTTACATCTATAGGAAAGGCTCTCCACCAAGTCTTTGGAGCAATCTATGGAGCTGCCTTCAGTGGGGTTTCATGGACTATGAAAATCCTCATAGGAGTCATTATCACATGGATAGGAATGAATTCACGCAGCACCTCACTGTCTGTGACACTAGTATTGGTGGGAATTGTGACACTGTATTTGGGAGTCATGGTGCAGGCCGATAGTGGTTGCGTTGTGAGCTGGAAAAACAAAGAACTGAAATGTGGCAGTGGGATTTTCATCACAGACAACGTGCACACATGGACAGAACAATACAAGTTCCAACCAGAATCCCCTTCAAAACTAGCTTCAGCTATCCAGAAAGCCCATGAAGAGGGCATTTGTGGAATCCGCTCAGTAACAAGACTGGAGAATCTGATGTGGAAACAAATAACACCAGAATTGAATCACATTCTATCAGAAAATGAGGTGAAGTTAACTATTATGACAGGAGACATCAAAGGAATCATGCAGGCAGGAAAACGATCTCTGCGGCCTCAGCCCACTGAGCTGAAGTATTCATGGAAAACATGGGGCAAAGCAAAAATGCTCTCTACAGAGTCTCATAACCAGACCTTTCTCATTGATGGCCCCGAAACAGCAGAATGCCCCAACACAAATAGAGCTTGGAATTCGTTGGAAGTTGAAGACTATGGCTTTGGAGTATTCACCACCAATATATGGCTAAAATTGAAAGAAAAACAGGATGTATTCTGCGACTCAAAACTCATGTCAGCGGCCATAAAAGACAACAGAGCCGTCCATGCCGATATGGGTTATTGGATAGAAAGTGCACTCAATGACACATGGAAGATAGAGAAAGCCTCTTTCATTGAAGTTAAAAACTGCCACTGGCCAAAATCACACACCCTCTGGAGCAATGGAGTGCTAGAAAGTGAGATGATAATTCCAAAGAATCTCGCTGGACCAGTGTCTCAACACAACTATAGACCAGGCTACCATACACAAATAACAGGACCATGGCATCTAGGTAAGCTTGAGATGGACTTTGATTTCTGTGATGGAACAACAGTGGTAGTGACTGAGGACTGCGGAAATAGAGGACCCTCTTTGAGAACAACCACTGCCTCTGGAAAACTCATAACAGAATGGTGCTGCCGATCTTGCACATTACCACCGCTAAGATACAGAGGTGAGGATGGGTGCTGGTACGGGATGGAAATCAGACCATTGAAGGAGAAAGAAGAGAATTTGGTCAACTCCTTGGTCACAGCTGGACATGGGCAGGTCGACAACTTTTCACTAGGAGTCTTGGGAATGGCATTGTTCCTGGAGGAAATGCTTAGGACCCGAGTAGGAACGAAACATGCAA

>DENV2_16681_**Fragment-II**

AACTTTTCACTAGGAGTCTTGGGAATGGCATTGTTCCTGGAGGAAATGCTTAGGACCCGAGTAGGAACGAAACATGCAATACTACTAGTTGCAGTTTCTTTTGTGACATTGATCACAGGGAACATGTCCTTTAGAGACCTGGGAAGAGTGATGGTTATGGTAGGCGCCACTATGACGGATGACATAGGTATGGGCGTGACTTATCTTGCCCTACTAGCAGCCTTCAAAGTCAGACCAACTTTTGCAGCTGGACTACTCTTGAGAAAGCTGACCTCCAAGGAATTGATGATGACTACTATAGGAATTGTACTCCTCTCCCAGAGCACCATACCAGAGACCATTCTTGAGTTGACTGATGCGTTAGCCTTAGGCATGATGGTCCTCAAAATGGTGAGAAATATGGAAAAGTATCAATTGGCAGTGACTATCATGGCTATCTTGTGCGTCCCAAACGCAGTGATATTACAAAACGCATGGAAAGTGAGTTGCACAATATTGGCAGTGGTGTCCGTTTCCCCACTGCTCTTAACATCCTCACAGCAAAAAACAGATTGGATACCATTAGCATTGACGATCAAAGGTCTCAATCCAACAGCTATTTTTCTAACAACCCTCTCAAGAACCAGCAAGAAAAGGAGCTGGCCATTAAATGAGGCTATCATGGCAGTCGGGATGGTGAGCATTTTAGCCAGTTCTCTCCTAAAAAATGATATTCCCATGACAGGACCATTAGTGGCTGGAGGGCTCCTCACTGTGTGCTACGTGCTCACTGGACGATCGGCCGATTTGGAACTGGAGAGAGCAGCCGATGTCAAATGGGAAGACCAGGCAGAGATATCAGGAAGCAGTCCAATCCTGTCAATAACAATATCAGAAGATGGTAGCATGTCGATAAAAAATGAAGAGGAAGAACAAACACTGACCATACTCATTAGAACAGGATTGCTGGTGATCTCAGGACTTTTTCCTGTATCAATACCAATCACGGCAGCAGCATGGTACCTGTGGGAAGTGAAGAAACAACGGGCCGGAGTATTGTGGGATGTTCCTTCACCCCCACCCATGGGAAAGGCTGAACTGGAAGATGGAGCCTATAGAATTAAGCAAAAAGGGATTCTTGGATATTCCCAGATCGGAGCCGGAGTTTACAAAGAAGGAACATTCCATACAATGTGGCATGTCACACGTGGCGCTGTTCTAATGCATAAAGGAAAGAGGATTGAACCATCATGGGCGGACGTCAAGAAAGACCTAATATCATATGGAGGAGGCTGGAAGTTAGAAGGAGAATGGAAGGAAGGAGAAGAAGTCCAGGTATTGGCACTGGAGCCTGGAAAAAATCCAAGAGCCGTCCAAACGAAACCTGGTCTTTTCAAAACCAACGCCGGAACAATAGGTGCTGTATCTCTGGACTTTTCTCCTGGAACGTCAGGATCTCCAATTATCGACAAAAAAGGAAAAGTTGTGGGTCTTTATGGTAATGGTGTTGTTACAAGGAGTGGAGCATATGTGAGTGCTATAGCCCAGACTGAAAAAAGCATTGAAGACAACCCAGAGATCGAAGATGACATTTTCCGAAAGAGAAGACTGACCATCATGGACCTCCACCCAGGAGCGGGAAAGACGAAGAGATACCTTCCGGCCATAGTCAGAGAAGCTATAAAACGGGGTTTGAGAACATTAATCTTGGCCCCCACTAGAGTTGTGGCAGCTGAAATGGAGGAAGCCCTTAGAGGACTTCCAATAAGATACCAGACCCCAGCCATCAGAGCTGAGCACACCGGGCGGGAGATTGTGGACCTAATGTGTCATGCCACATTTACCATGAGGCTGCTATCACCAGTTAGAGTGCCAAACTACAACCTGATTATCATGGACGAAGCCCATTTCACAGACCCAGCAAGTATAGCAGCTAGAGGATACATCTCAACTCGAGTGGAGATGGGTGAGGCAGCTGGGATTTTTATGACAGCCACTCCCCCGGGAAGCAGAGACCCATTTCCTCAGAGCAATGCACCAATCATAGATGAAGAAAGAGAAATCCCTGAACGTTCGTGGAATTCCGGACATGAATGGGTCACGGATTTTAAAGGGAAGACTGTTTGGTTCGTTCCAAGTATAAAAGCAGGAAATGATATAGCAGCTTGCCTGAGGAAAAATGGAAAGAAAGTGATACAACTCAGTAGGAAGACCTTTGATTCTGAGTATGTCAAGACTAGAACCAATGATTGGGACTTCGTGGTTACAACTGACATTTCAGAAATGGGTGCCAATTTCAAGGCTGAGAGGGTTATAGACCCCAGACGCTGCATGAAACCAGTCATACTAACAGATGGTGAAGAGCGGGTGATTCTGGCAGGACCTATGCCAGTGACCCACTCTAGTGCAGCACAAAGAAGAGGGAGAATAGGAAGAAATCCAAAAAATGAGAATGACCAGTACATATACATGGGGGAACCTCTGGAAAATGATGAAGACTGTGCACACTGGAAAGAAGCTAAAATGCTCCTAGATAACATCAACACGCCAGAAGGAATCATTCCTAGCATGTTCGAACCAGAGCGTGAAAAGGTGGATGCCATTGATGGCGAATACCGCTTGAGAGGAGAAGCAAGGAAAACCTTTGTAGACTTAATGAGAAGAGGAGACCTACCAGTCTGGTTGGCCTACAGAGTGGCAGCTGAAGGCATCAACTACGCAGACAGAAGGTGGTGTTTTGATGGAGTCAAGAACAACCAAATCCTAGAAGAAAACGTGGAAGTTGAAATCTGGACAAAAGAAGGGGAAAGGAAGAAATTGAAACCCAGATGGTTGGATGCTAGGATCTATTCTGACCCACTGGCGCTAAAAGAATTTAAGGAATTTGCAGCCGGAAGAAAGTCTCTGACCCTGAACCTAATCACAGAAATGGGTAGGCTCCCAACCTTCATGACTCAGAAGGCAAGAGACGCACTGGACAACTTAGCAGTGCTGCACACGGCTGAGGCAGGTGGAAGGGCGTACAACCATGCTCTCAGTGAACTGCCGGAGACCCTGGAGACATTGCTTTTACTGACACTTCTGGCTACAGTCACGGGAGGGATCTTTTTATTCTTGATGAGCGGAAGGGGCATAGGGAAGATGACCCTGGGAATGTGCTGCATAATCACGGCTAGCATCCTCCTATGGTACGCACAAATACAGCCACACTGGATAGCAGCTTCAATAATACTGGAGTTTTTTCTCATAGTTTTGCTTATTCCAGAACCTGAAAAACAGAGAACACCCCAAGACAACCAACTGACCTACGTTGTCATAGCCATCCTCACAGTGGTGGCCGCAACCATGGCAAACGAGATGGGTTTCCTAGAAAAAACGAAGAAAGATCTCGGATTGGGAAGCATTGCAACCCAGCAACCCGAGAGCAACATCCTGGACATAGATCTACGTCCTGCATCAGCATGGACGCTGTATGCCGTGGCCACAACATTTGTTACACCAATGTTGAGACATAGCATTGAAAATTCCTCAGTGAATGTGTCCCTAACAGCTATAGCCAACCAAGCCACAGTGTTAATGGGTCTCGGGAAAGGATGGCCATTGTCAAAGATGGACATCGGAGTTCCCCTTCTCGCCATTGGATGCTACTCACAAGTCAACCCCATAACTCTCACAGCAGCTCTTTTCTTATTGGTAGCACATTATGCCATCATAGGGCCAGGACTCCAAGCAAAAGCAACCAGAGAAGCTCAG

>DENV2_16681_**Fragment-III**

CAGCTCTTTTCTTATTGGTAGCACATTATGCCATCATAGGGCCAGGACTCCAAGCAAAAGCAACCAGAGAAGCTCAGAAAAGAGCAGCGGCGGGCATCATGAAAAACCCAACTGTCGATGGAATAACAGTGATTGACCTAGATCCAATACCTTATGATCCAAAGTTTGAAAAGCAGTTGGGACAAGTAATGCTCCTAGTCCTCTGCGTGACTCAAGTATTGATGATGAGGACTACATGGGCTCTGTGTGAGGCTTTAACCTTAGCTACCGGGCCCATCTCCACATTGTGGGAAGGAAATCCAGGGAGGTTTTGGAACACTACCATTGCGGTGTCAATGGCTAACATTTTTAGAGGGAGTTACTTGGCCGGAGCTGGACTTCTCTTTTCTATTATGAAGAACACAACCAACACAAGAAGGGGAACTGGCAACATAGGAGAGACGCTTGGAGAGAAATGGAAAAGCCGATTGAACGCATTGGGAAAAAGTGAATTCCAGATCTACAAGAAAAGTGGAATCCAGGAAGTGGATAGAACCTTAGCAAAAGAAGGCATTAAAAGAGGAGAAACGGACCATCACGCTGTGTCGCGAGGCTCAGCAAAACTGAGATGGTTCGTTGAGAGAAACATGGTCACACCAGAAGGGAAAGTAGTGGACCTCGGTTGTGGCAGAGGAGGCTGGTCATACTATTGTGGAGGACTAAAGAATGTAAGAGAAGTCAAAGGCCTAACAAAAGGAGGACCAGGACACGAAGAACCCATCCCCATGTCAACATATGGGTGGAATCTAGTGCGTCTTCAAAGTGGAGTTGACGTTTTCTTCATCCCGCCAGAAAAGTGTGACACATTATTGTGTGACATAGGGGAGTCATCACCAAATCCCACAGTGGAAGCAGGACGAACACTCAGAGTCCTTAACTTAGTAGAAAATTGGTTGAACAACAACACTCAATTTTGCATAAAGGTTCTCAACCCATATATGCCCTCAGTCATAGAAAAAATGGAAGCACTACAAAGGAAATATGGAGGAGCCTTAGTGAGGAATCCACTCTCACGAAACTCCACACATGAGATGTACTGGGTATCCAATGCTTCCGGGAACATAGTGTCATCAGTGAACATGATTTCAAGGATGTTGATCAACAGATTTACAATGAGATACAAGAAAGCCACTTACGAGCCGGATGTTGACCTCGGAAGCGGAACCCGTAACATCGGGATTGAAAGTGAGATACCAAACCTAGATATAATTGGGAAAAGAATAGAAAAAATAAAGCAAGAGCATGAAACATCATGGCACTATGACCAAGACCACCCATACAAAACGTGGGCATACCATGGTAGCTATGAAACAAAACAGACTGGATCAGCATCATCCATGGTCAACGGAGTGGTCAGGCTGCTGACAAAACCTTGGGACGTCGTCCCCATGGTGACACAGATGGCAATGACAGACACGACTCCATTTGGACAACAGCGCGTTTTTAAAGAGAAAGTGGACACGAGAACCCAAGAACCGAAAGAAGGCACGAAGAAACTAATGAAAATAACAGCAGAGTGGCTTTGGAAAGAATTAGGGAAGAAAAAGACACCCAGGATGTGCACCAGAGAAGAATTCACAAGAAAGGTGAGAAGCAATGCAGCCTTGGGGGCCATATTCACTGATGAGAACAAGTGGAAGTCGGCACGTGAGGCTGTTGAAGATAGTAGGTTTTGGGAGCTGGTTGACAAGGAAAGGAATCTCCATCTTGAAGGAAAGTGTGAAACATGTGTGTACAACATGATGGGAAAAAGAGAGAAGAAGCTAGGGGAATTCGGCAAGGCAAAAGGCAGCAGAGCCATATGGTACATGTGGCTTGGAGCACGCTTCTTAGAGTTTGAAGCCCTAGGATTCTTAAATGAAGATCACTGGTTCTCCAGAGAGAACTCCCTGAGTGGAGTGGAAGGAGAAGGGCTGCACAAGCTAGGTTACATTCTAAGAGACGTGAGCAAGAAAGAGGGAGGAGCAATGTATGCCGATGACACCGCAGGATGGGATACAAGAATCACACTAGAAGACCTAAAAAATGAAGAAATGGTAACAAACCACATGGAAGGAGAACACAAGAAACTAGCCGAGGCCATTTTCAAACTAACGTACCAAAACAAGGTGGTGCGTGTGCAAAGACCAACACCAAGAGGCACAGTAATGGACATCATATCGAGAAGAGACCAAAGAGGTAGTGGACAAGTTGGCACCTATGGACTCAATACTTTCACCAATATGGAAGCCCAACTAATCAGACAGATGGAGGGAGAAGGAGTCTTTAAAAGCATTCAGCACCTAACAATCACAGAAGAAATCGCTGTGCAAAACTGGTTAGCAAGAGTGGGGCGCGAAAGGTTATCAAGAATGGCCATCAGTGGAGATGATTGTGTTGTGAAACCTTTAGATGACAGGTTCGCAAGCGCTTTAACAGCTCTAAATGACATGGGAAAGATTAGGAAAGACATACAACAATGGGAACCTTCAAGAGGATGGAATGATTGGACACAAGTGCCCTTCTGTTCACACCATTTCCATGAGTTAATCATGAAAGACGGTCGCGTACTCGTTGTTCCATGTAGAAACCAAGATGAACTGATTGGCAGAGCCCGAATCTCCCAAGGAGCAGGGTGGTCTTTGCGGGAGACGGCCTGTTTGGGGAAGTCTTACGCCCAAATGTGGAGCTTGATGTACTTCCACAGACGCGACCTCAGGCTGGCGGCAAATGCTATTTGCTCGGCAGTACCATCACATTGGGTTCCAACAAGTCGAACAACCTGGTCCATACATGCTAAACATGAATGGATGACAACGGAAGACATGCTGACAGTCTGGAACAGGGTGTGGATTCAAGAAAACCCATGGATGGAAGACAAAACTCCAGTGGAATCATGGGAGGAAATCCCATACTTGGGGAAAAGAGAAGACCAATGGTGCGGCTCATTGATTGGGTTAACAAGCAGGGCCACCTGGGCAAAGAACATCCAAGCAGCAATAAATCAAGTTAGATCCCTTATAGGCAATGAAGAATACACAGATTACATGCCATCCATGAAAAGATTCAGAAGAGAAGAGGAAGAAGCAGGAGTTCTGTGGTAGAAAGCAAAACTAACATGAAACAAGGCTAGAAGTCAGGTCGGATTAAGCCATAGTACGGAAAAAACTATGCTACCTGTGAGCCCCGTCCAAGGACGTTAAAAGAAGTCAGGCCATCATAAATGCCATAGCTTGAGTAAACTATGCAGCCTGTAGCTCCACCTGAGAAGGTGTAAAAAATCCGGGAGGCCACAAACCATGGAAGCTGTACGCATGGCGTAGTGGACTAGCGGTTAGAGGAGACCCCTCCCTTACAAATCGCAGCAACAATGGGGGCCCAAGGCGAGATGAAGCTGTAGTCTCGCTGGAAGGACTAGAGGTTAGAGGAGACCCCCCCGAAACAAAAAACAGCATATTGACGCTGGGAAAGACCAGAGATCCTGCTGTCTCCTCAGCATCATTCCAGGCACAGAACGCCAGAAAATGGAATGGTGCTGTTGAATCAACAGGTTCT**GGCCGGCATGGTCCCAGCCTCCTCGCTGGCGCCGGCTGGGCAACATTCCGAGGGGACCGTCCCCTCGGTAATGGCGAATGGGACTCGCGACAGACATGATAAGATACATTGATGAGTTTGGACAAACCACAACTAGAATGCAGTGAAAAAAATGCTTTATTTGTGAAATTAAGCGCTGGCATTGACCCTGAG**

**JEV wild-type (JEV-WT) ISA Fragments**

JEV strain SA14-14-2 [GenBank: MK585066.1]

Highlighted in red – pCMV promoter sequence.

Highlighted in green – HDR/SV40pA sequence.

>wt-SA14-14-2_**Fragment-I**

The rescue of this fragment failed due to its toxicity to bacteria. The alternative ISA strategy is described below.

**CACCCAACTGATCTTCAGCATCTTCAATATTGGCCATTAGCCATATTATTCATTGGTTATATAGCATAAATCAATATTGGCTATTGGCCATTGCATACGTTGTATCTATATCATAATATGTACATTTATATTGGCTCATGTCCAATATGACCGCCATGTTGGCATTGATTATTGACTAGTTATTAATAGTAATCAATTACGGGGTCATTAGTTCATAGCCCATATATGGAGTTCCGCGTTACATAACTTACGGTAAATGGCCCGCCTGGCTGACCGCCCAACGACCCCCGCCCATTGACGTCAATAATGACGTATGTTCCCATAGTAACGCCAATAGGGACTTTCCATTGACGTCAATGGGTGGAGTATTTACGGTAAACTGCCCACTTGGCAGTACATCAAGTGTATCATATGCCAAGTCCGCCCCCTATTGACGTCAATGACGGTAAATGGCCCGCCTGGCATTATGCCCAGTACATGACCTTACGGGACTTTCCTACTTGGCAGTACATCTACGTATTAGTCATCGCTATTACCATGGTGATGCGGTTTTGGCAGTACACCAATGGGCGTGGATAGCGGTTTGACTCACGGGGATTTCCAAGTCTCCACCCCATTGACGTCAATGGGAGTTTGTTTTGGCACCAAAATCAACGGGACTTTCCAAAATGTCGTAATAACCCCGCCCCGTTGACGCAAATGGGCGGTAGGCGTGTACGGTGGGAGGTCTATATAAGCAGAGCTCGTTTAGTGAACCG**AGAAGTTTATCTGTGTGAACTTCTTGGCTTAGTATCGTAGAGAAGAATCGAGAGATTAATGCAGTTTAAACAGTTTTTTAGAACGGAAGATAACCATGACTAAAAAACCAGGAGGGCCCGGTAAAAACCGGGCTATCAATATGCTGAAACGCGGCCTACCCCGCGTATTCCCACTAGTGGGAGTGAAGAGGGTAGTAATGAGCTTGTTGGACGGCAGAGGGCCAGTACGTTTCGTGCTGGCTCTTATCACGTTCTTCAAGTTTACAGCATTAGCCCCGACCAAGGCGCTTTCAGGCCGATGGAAAGCAGTGGAAAAGAGTGTGGCAATGAAACATCTTACTAGTTTCAAACGAGAACTTGGAACACTCATTGACGCCGTGAACAAGCGGGGCAGAAAGCAAAACAAAAGAGGAGGAAATGAAGGCTCAATCATGTGGCTCGCGAGCTTGGCAGTTGTCATAGCTTGTGCAGGAGCCATGAAGTTGTCGAATTTCCAGGGGAAGCTTTTGATGACCATTAACAACACGGACATTGCAGACGTTATCGTGATTCCCACCTCAAAAGGAGAGAACAGATGCTGGGTCCGGGCAATCGACGTCGGCTACATGTGTGAGGACACTATCACGTACGAATGTCCTAAGCTTACCATGGGCAATGATCCAGAGGATGTGGATTGCTGGTGTGACAACCAAGAAGTCTACGTCCAATATGGACGGTGCACGCGGACCAGGCATTCCAAGCGAAGCAGGAGATCCGTGTCGGTCCAAACACATGGGGAGAGTTCACTAGTGAATAAAAAAGAGGCTTGGCTGGATTCAACGAAAGCCACACGATATCTCATGAAAACTGAGAACTGGATCATAAGGAATCCTGGCTATGCTTTTCTGGCGGCGGTACTTGGCTGGATGCTTGGCAGTAACAACGGTCAACGCGTGGTATTTACCATCCTCCTGCTGTTGGTCGCTCCGGCTTACAGTTTTAATTGTCTGGGAATGGGCAATCGTGACTTCATAGAAGGAGCCAGTGGAGCCACTTGGGTGGACTTGGTGCTAGAAGGAGACAGCTGCTTGACAATCATGGCAAACAACAAACCAACATTGGACGTCCGCATGATTAACATCGAAGCTAGCCAACTTGCTGAGGTCAGAAGTTACTGCTATCATGCTTCAGTCACTGACATCTCGACGGTGGCTCGGTGCCCCACGACTGGAGAAGCCCACAACGAGAAGCGAGCTGATAGTAGCTATGTGTGCAAACAAGGCTTCACTGACCGTGGGTGGGGCAACGGATGTGGATTTTTCGGGAAGGGAAGCATTGACACATGTGCAAAATTCTCCTGCACCAGTAAAGCGATTGGGAGAACAATCCAGCCAGAAAACATCAAATACAAAGTTGGCATTTTTGTGCATGGAACCACCACTTCGGAAAACCATGGGAATTATTCAGCGCAAGTTGGGGCGTCCCAGGCGGCAAAGTTTACAGTAACACCCAATGCTCCTTCGGTAGCCCTCAAACTTGGTGACTACGGAGAAGTCACACTGGACTGTGAGCCAAGGAGTGGACTGAACACTGAAGCGTTTTACGTCATGACCGTGGGGTCAAAGTCATTTCTGGTCCATAGGGAGTGGTTTCATGACCTCGCTCTCCCCTGGACGTCCCCTTCGAGCACAGCGTGGAGAAACAGAGAACTCCTCATGGAATTTGAAGGGGCGCACGCCACAAAACAGTCCGTTGTTGCTCTTGGGTCACAGGAAGGAGGCCTCCATCATGCGTTGGCAGGAGCCATCGTGGTGGAGTACTCAAGCTCAGTGATGTTAACATCAGGCCACCTGAAATGTAGGCTGAAAATGGACAAACTGGCTCTGAAAGGCACAACCTATGGCATGTGTACAGAAAAATTCTCGTTCGCGAAAAATCCGGTGGACACTGGTCACGGAACAGTTGTCATTGAACTCTCCTACTCTGGGAGTGATGGCCCCTGCAAAATTCCGATTGTTTCCGTTGCGAGCCTCAATGACATGACCCCCGTTGGGCGGCTGGTGACAGTGAACCCCTTCGTCGCGACTTCCAGTGCCAACTCAAAGGTGCTGGTCGAGATGGAACCCCCCTTCGGAGACTCCTACATCGTAGTTGGAAGGGGAGCCAAGCAGATCAACCACCATTGGCACAAAGCTGGAAGCACGCTGGGCAAGGCCTTTTCAACAACTTTGAAGGGAGCTCAAAGACTGGCAGCGTTGGGCGACACAGCCTGGGACTTTGGCTCTATTGGAGGGGTCTTCAACTCCATAGGAAGAGCCGTTCACCAAGTGTTTGGTGGTGCCTTCAGAACACTCTTTGGGGGAATGTCTTGGATCACACAAGGGCTAATGGGTGCCCTACTGCTCTGGATGGGCGTCAACGCACGAGACCGATCAATTGCTTTGGCCTTCTTAGCCACAGGAGGTGTGCTCGTGTTCTTAGCGACCAATGTGCATGCTGACACTGGATGTGCCATTGACATCACAAGAAAAGAGATGAGATGTGGAAGTGGCATCTTCGTGCACAACGACGTGGAAGCCTGGGTGGATAGGTATAAATATTTGCCAGAAACGCCCAGATCCCTAGCGAAGATCGTCCACAAAGCGCACAAGGAAGGCGTGTGCGGAGTCAGATCTGTCACTAGACTGGAGCACCAAATGTGGGAAGCCGTAAGGGACGAATTGAACGTCCTGCTCAAAGAGAATGCAGTGGACCTCAGTGTGGTTGTGAACAAGCCCGTGGGAAGATATCGCTCAGCCCCTAAACGCCTATCCATGACGCAAGAGAAGTTTGAAATGGGCTGGAAAGCATGGGGAAAAAGCATCCTCTTTGCCCCGGAATTGGCTAACTCCACATTTGTCGTAGATGGACCTGAGACAAAGGAATGCCCTGATGAGCACAGAGCTTGGAACAGCATGCAAATCGAAGACTTCGGCTTTGGCATCACATCAACCCGTGTGTGGCTGAAAATTAGAGAGGAGAGCACTGACGAGTGTGATGGAGCGATCATAGGCACGGCTGTCAAAGGACATGTGGCAGTCCATAGTGACTTGTCGTACTGGATTGAGAGTCGCTACAACGACACATGGAAACTTGAGAGGGCAGTCTTTGGAGAGGTCAAATCTTGCACTTGGCCAGAGACACACACCCTTTGGGGAGATGATGTTGAGGAAAGTGAACTCATCATTCCGCACACCATAGCCGGACCAAAAAGCAAGCACAATCGGAGGGAAGGGTATAAGACACAAAACCAGGGACCTTGGGATGAGAATGGCATAGTCTTGGACTTTGATTATTGCCCAGGGACAAAAGTCACCATTACAGAGGATTGTAGCAAGAGAGGCCCTTCGGTCAGAACCACTACTGACAGTGGAAAGTTGATCACTGACTGGTGCTGTCGCAGTTGCTCCCTTCCGCCCCTACGATTCCGGACAGAAAATGGCTGCTGGTACGGAATGGAAATCAGACCTGTTATGCATGATGAAACAACACTCGTCAGATCACAGGTTCATGCTTTCAAAGGTGAAATGGTTGACCCTTTTCAGCTGGGCCTTCTGGTGATGTTTCTGGCCACCCAGGAAGTCCTTCGCAAGAGGTGGACGGCCAGATTGACCATTCCTGCGGTTTTGGGGGTCCTACTTGTGCTGATGCTTGGGGGTATCACTTACACTGATTTGGCGAGGTATGTGGTGCTAGTCGCTGCTGCTTTCGCAGAGGCCAACAGTGGAGGAGACGTCCTGCACCTTGCTTTGATTGCTGTTTTTAAGATCCAAC

>wt-SA14-14-2_**Fragment-II**

GCTTTCGCAGAGGCCAACAGTGGAGGAGACGTCCTGCACCTTGCTTTGATTGCTGTTTTTAAGATCCAACCAGCATTTTTAGTGATGAACATGCTTAGCACGAGATGGACGAACCAAGAAAACGTGGTTCTGGTCCTAGGGGCTGCCTTTTTCCAATTGGCCTCAGTAGATCTGCAAATAGGAGTCCACGGAATCCTGAATGCCGCCGCTATAGCATGGATGATTGTCCGAGCGATCACCTTCCCCACAACCTCCTCCGTCACCATGCCAGTCTTAGCGCTTCTAACTCCGGGGATGAGGGCTCTATACCTAGACACTTACAGAATCATCCTCCTCGTCATAGGGATTTGCTCCCTGCTGCACGAGAGGAAAAAGACCATGGCGAAAAAGAAAGGAGCTGTACTCTTGGGCTTAGCGCTCACATCCACTGGATGGTTCTCGCCCACCACTATAGCTGCCGGACTAATGGTCTGCAACCCAAACAAGAAGAGAGGGTGGCCAGCTACTGAGTTTTTGTCGGCAGTTGGATTGATGTTTGCCATCGTAGGTGGTTTGGCCGAGTTGGATATTGAATCCATGTCAATACCCTTCATGCTGGCAGGTCTCATGGCAGTGTCCTACGTGGTGTCAGGAAAAGCAACAGATATGTGGCTTGAACGGGCCGCCGACATCAGCTGGGATATGGGTGCTGCAATCACAGGAAGCAGTCGGAGGCTGGATGTGAAACTGGATGATGACGGAGATTTTCACTTGATTGATGATCCCGGTGTTCCATGGAAGGTCTGGGTCCTGCGCATGTCTTGCATTGGCTTAGCCGCCCTCACGCCTTGGGCCATCGTTCCCGCCGCTTTCGGTTATTGGCTCACTTTAAAAACAACAAAAAGAGGGGGCGTGTTTTGGGACACGCCATCCCCAAAACCTTGCTCAAAAGGAGACACCACTACAGGAGTCTACCGAATTATGGCTAGAGGGATTCTTGGCACTTACCAGGCCGGCGTCGGAGTCATGTACGAGAATGTTTTCCACACACTATGGCACACAACTAGAGGAGCAGCCATTGTGAGTGGAGAAGGAAAATTGACGCCATACTGGGGTAGTGTGAAAGAAGACCGCATAGCTTACGGAGGCCCATGGAGGTTCGACCGAAAATGGAATGGAACAGATGACGTGCAAGTGATCGTGGTAGAACCGGGGAAGGGCGCAGTAAACATCCAGACAAAACCAGGAGTGTTTCGGACTCCCTTCGGGGAGGTTGGGGCTGTTAGTCTGGATTACCCGCGAGGAACATCCGGCTCACCCATTCTGGATTCCAATGGAGACATTATAGGCCTATACGGCAATGGAGTTGAGCTTGGCGATGGCTCATACGTCAGCGCCATCGTGCAGGGTGACCGTCAGGAGGAACCAGTCCCAGAAGCTTACACCCCAAACATGTTGAGAAAGAGACAGATGACTGTGCTAGATTTGCACCCTGGTTCAGGGAAAACCAGGAAAATTCTGCCACAAATAATTAAGGACGCTATCCAGCAGCGCCTAAGAACAGCTGTGTTGGCACCGACGCGGGTGGTAGCAGCAGAAATGGCAGAAGTTTTGAGAGGGCTCCCAGTACGATATCAAACTTCAGCAGTGCAGAGAGAGCACCAAGGGAATGAAATAGTGGATGTGATGTGCCACGCCACTCTGACCCATAGACTGATGTCACCGAACAGAGTGCCCAACTACAACCTATTTGTCATGGATGAAGCTCATTTCACCGACCCAGCCAGTATAGCCGCACGAGGATACATTGCTACCAAGGTGGAATTAGGGGAGGCAGCAGCCATCTTTATGACAGCGACCCCGCCTGGAACCACGGATCCTTTTCCTGACTCAAATGCCCCAATCCATGATTTGCAAGATGAGATACCAGACAGGGCATGGAGCAGTGGATACGAATGGATCACAGAATATGCGGGTAAAACCGTGTGGTTTGTGGCGAGCGTAAAAATGGGGAATGAGATTGCAATGTGCCTCCAAAGAGCGGGGAAAAAGGTCATCCAACTCAACCGCAAGTCCTATGACACAGAATACCCAAAATGTAAGAATGGAGACTGGGATTTTGTCATTACCACCGACATCTCTGAAATGGGGGCCAACTTCGGTGCGAGCAGGGTCATCGACTGTAGAAAGAGCGTGAAACCCACCATCTTAGAAGAGGGAGAAGGCAGAGTCATCCTCGGAAACCCATCTCCCATAACCAGTGCAAGCGCAGCTCAACGGAGGGGCAGAGTAGGCAGAAACCCCAATCAAGTTGGAGATGAATACCACTATGGAGGGGCTACCAGTGAAGATGACAGTAACCTAGCCCATTGGACAGAGGCAAAGATCATGTTAGACAACATACACATGCCCAATGGACTGGTGGCCCAGCTCTATGGACCAGAGAGGGAAAAGGCTTTCACAATGGATGGCGAATACCGTCTCAGAGGTGAAGAAAAGAAAAACTTCTTAGAGCTGCTTAGGACGGCTGACCTCCCGGTGTGGCTGGCCTACAAGGTGGCGTCCAATGGCATTCAGTACACCGACAGAAAGTGGTGTTTTGATGGGCCGCGTACGAATGCCATACTGGAGGACAACACCGAGGTAGAGATAGTCACCCGGATGGGTGAGAGGAAAATCCTCAAGCCGAGATGGCTTGATGCAAGAGTTTATGCAGATCACCAGGCCCTCAAGTGGTTCAAAGACTTTGCAGCAGGGAAGAGATCAGCCGTTAGCTTCATAGAGGTGCTCGGTCGCATGCCTGAGCATTTCATGGGAAAGACGCGGGAAGCTTTAGACACCATGTACTTGGTTGCAACGGCTGAGAAAGGTGGGAAAGCACACCGAATGGCTCTCGAAGAGCTGCCAGATGCACTGGAAACCATCACACTTATTGTCGCCATTACTGTGATGACAGGAGGATTCTTCCTACTAATGATGCAGCGAAAGGGTATAGGGAAGATGGGTCTTGGAGCTCTAGTGCTCACACTAGCTACCTTCTTCCTGTGGGCGGCAGAGGTTCCTGGAACCAAAATAGCAGGGACCCTGCTGATCGCCCTGCTGCTGATGGTGGTTCTCATCCCAGAACCGGAAAAACAGAGGTCACAGACAGATAACCAACTGGCGGTGTTTCTCATCTGTGTCTTGACCGTGGTTGGAGTGGTGGCAGCAAACGAGTACGGGATGCTAGAAAAAACCAAAGCGGATCTCAAGAGCATGTTTGGCGGAAAGACGCAGGCATCAGGACTGACTGGATTGCCAAGCATGGCACTGGACCTGCGTCCAGCCACAGCCTGGGCACTGTATGGGGGGAGCACAGTCGTGCTAACCCCTCTTCTGAAGCACCTGATCACGTCGGAATACGTCACCACATCGCTAGCTTCAATTAACTCACAAGCTGGCTCATTATTCGTCTTGCCACGAGGCGTGCCTTTTACCGACCTAGACTTGACTGTTGGCCTCGTCTTCCTTGGCTGTTGGGGTCAAGTCACCCTCACAACGTTTCTGACAGCCATGGTTCTGGCGACACTTCACTATGGGTACATGCTCCCTGGATGGCAAGCAGAAGCACTCAGGGCTGCCCAGAGAAGGACAGC

>wt-SA14-14-2_**Frgament-III**

GACACTTCACTATGGGTACATGCTCCCTGGATGGCAAGCAGAAGCACTCAGGGCTGCCCAGAGAAGGACAGCGGCTGGAATAATGAAGAATGCCGTTGTTGACGGAATGGTCGCCACTGATGTGCCTGAACTGGAAAGGACTACTCCTCTGATGCAAAAGAAAGTCGGACAGGTGCTCCTCATAGGGGTAAGCGTGGCAGCGTTCCTCGTCAACCCTAATGTCACCACTGTGAGAGAAGCAGGGGTGTTGGTGACGGCGGCTACGCTTACTTTGTGGGACAATGGAGCCAGTGCCGTTTGGAATTCCACCACAGCCACGGGACTCTGCCATGTCATGCGAGGTAGCTACCTGGCTGGAGGCTCCATTGCTTGGACTCTCATCAAGAACGCTGATAAGCCCTCCTTGAAAAGGGGAAGGCCTGGGGGCAGGACGCTAGGGGAGCAGTGGAAGGAAAAACTAAATGCCATGAGTAGAGAAGAGTTTTTTAAATACCGGAGAGAGGCCATAATCGAGGTGGACCGCACTGAAGCACGCAGGGCCAGACGTGAAAATAACATAGTGGGAGGACATCCGGTTTCGCGAGGCTCAGCAAAACTCCGTTGGCTCGTGGAGAAAGGATTTGTCTCGCCAATAGGAAAAGTCATTGATCTAGGGTGTGGGCGTGGAGGATGGAGCTACTACGCAGCAACCCTGAAGAAGGTCCAGGAAGTCAGAGGATACACGAAAGGTGGGGCGGGACATGAAGAACCGATGCTCATGCAGAGCTACGGCTGGAACCTGGTCTCCCTGAAGAGTGGAGTGGACGTGTTTTACAAACCTTCAGAGCCCAGTGATACCCTGTTCTGTGACATAGGGGAATCCTCCCCAAGTCCAGAAGTAGAAGAACAACGCACACTACGCGTCCTAGAGATGACATCTGACTGGTTGCACCGAGGACCTAGAGAGTTCTGCATTAAAGTTCTCTGCCCTTACATGCCCAAGGTTATAGAAAAAATGGAAGTTCTGCAGCGTCGCTTCGGAGGTGGGCTAGTGCGTCTCCCCCTGTCCCGAAACTCCAATCACGAGATGTATTGGGTTAGTGGAGCCGCTGGCAATGTGGTGCACGCTGTGAACATGACCAGCCAGGTATTACTGGGGCGAATGGATCGCACAGTGTGGAGAGGGCCAAAGTATGAGGAAGATGTCAACCTAGGGAGCGGAACAAGAGCCGTGGGAAAGGGAGAAGTCCATAGCAATCAGGAGAAAATCAAGAAGAGAATCCAGAAGCTTAAAGAAGAATTCGCCACAACGTGGCACAAAGACCCTGAGCATCCATACCGCACTTGGACATACCACGGAAGCTATGAAGTGAAGGCTACTGGCTCAGCCAGCTCTCTCGTCAACGGAGTGGTGAAGCTCATGAGCAAACCTTGGGACGCCATTGCCAACGTCACCACCATGGCCATGACTGACACCACCCCTTTTGGACAGCAAAGAGTTTTCAAGGAGAAAGTTGACACGAAGGCTCCTGAGCCACCAGCTGGAGCCAAGGAAGTGCTCAACGAGACCACCAACTGGCTGTGGGCCTACTTGTCACGGGAAAAAAGACCCCGCTTGTGCACCAAGGAAGAATTCATTAAGAAAGTTAACAGCAACGCGGCTCTTGGAGCAGTGTTCGCTGAACAGAATCAATGGAGCACGGCGCGTGAGGCTGTGGATGACCCGCGGTTTTGGGAGATGGTTGATGAAGAGAGGGAAAACCATCTGCGAGGAGAGTGTCACACATGTATCTACAACATGATGGGAAAAAGAGAGAAGAAGCCTGGAGAGTTTGGAAAAGCTAAAGGAAGCAGGGCCATTTGGTTCATGTGGCTTGGAGCACGGTATCTAGAGTTTGAAGCTTTGGGGTTCCTGAATGAAGACCATTGGCTGAGCCGAGAGAATTCAGGAGGTGGAGTGGAAGGCTCAGGCGTCCAAAAGCTGGGATACATCCTCCGTGACATAGCAGGAAAGCAAGGAGGGAAAATGTACGCTGATGATACCGCCGGGTGGGACACTAGAATTACCAGAACTGATTTAGAAAATGAAGCTAAGGTACTGGAGCTCCTAGACGGTGAACACCGCATGCTCGCCCGAGCCATAATTGAACTGACTTACAGGCACAAAGTGGTCAAGGTCATGAGACCTGCAGCAGAAGGAAAGACCGTGATGGACGTGATATCAAGAGAAGATCAAAGGGGGAGTGGACAGGTGGTCACTTATGCTCTTAACACTTTCACGAACATCGCTGTCCAGCTCGTCAGGCTGATGGAGGCTGAGGGGGTCATTGGACCACAACACTTGGAACATCTACCTAGGAAAAACAAGATAGCTGTCAGGACCTGGCTCTTTGAGAATGGAGAGGAGAGAGTGACCAGGATGGCGATCAGCGGAGACGACTGTGCCGTCAAACCGCTGGACGACAGATTCGCCACAGCCCTCCACTTCCTCAACGCAATGTCAAAGGTCAGAAAAGACATCCAGGAATGGAAGCCTTCGCATGGCTGGCACGATTGGCAGCAAGTTCCCTTCTGTTCTAACCATTTTCAGGAGATTGTGATGAAAGATGGAAGGAGTATAGTTGTCCCGTGCAGAGGACAGGATGAGCTGATAGGCAGGGCTCGCATCTCTCCAGGAGCTGGATGGAATGTGAAGGACACAGCTTGCCTGGCCAAAGCATATGCACAGATGTGGCTACTCCTATACTTCCATCGCAGGGACTTGCGTCTCATGGCAAATGCGATTTGCTCAGCAGTGCCAGTAGATTGGGTGCCCACAGGCAGGACATCCTGGTCAATACACTCGAAAGGAGAGTGGATGACCACGGAAGACATGCTGCAGGTCTGGAACAGAGTTTGGATTGAAGAAAATGAATGGATGATGGACAAGACTCCAATCACAAGCTGGACAGACGTTCCGTATGTGGGAAAGCGCGAGGACATCTGGTGTGGCAGCCTCATCGGAACGCGATCCAGAGCAACCTGGGCTGAGAACATCTATGCGGCGATAAACCAGGTTAGAGCTGTCATTGGGAAAGAAAATTATGTTGACTACATGACCTCACTCAGGAGATACGAAGACGTCTTGATCCAGGAAGACAGGGTCATCTAGTGTGATTTAAGGTAGAAAAGTAGACTATGTAAACAATGTAAATGAGAAAATGCATGCATATGGAGTCAGGCCAGCAAAAGCTGCCACCGGATACTGGGTAGACGGTGCTGCCTGCGTCTCAGTCCCAGGAGGACTGGGTTAACAAATCTGACAACAGAAAGTGAGAAAGCCCTCAGAACCGTCTCGGAAGTAGGTCCCTGCTCACTGGAAGTTGAAAGACCAACGTCAGGCCACAAATTTGTGCCACTCCGCTAGGGAGTGCGGCCTGCGCAGCCCCAGGAGGACTGGGTTACCAAAGCCGTTGAGGCCCCCACGGCCCAAGCCTCGTCTAGGATGCAATAGACGAGGTGTAAGGACTAGAGGTTAGAGGAGACCCCGTGGAAACAACAACATGCGGCCCAAGCCCCCTCGAAGCTGTAGAGGAGGTGGAAGGACTAGAGGTTAGAGGAGACCCCGCATTTGCATCAAACAGCATATTGACACCTGGGAATAGACTGGGAGATCTTCTGCTCTATCTCAACATCAGCTACTAGGCACAGAGCGCCGAAGTATGTAGCTGGTGGTGAGGAAGAACACAGGATCT**GGCCGGCATGGTCCCAGCCTCCTCGCTGGCGCCGGCTGGGCAACATTCCGAGGGGACCGTCCCCTCGGTAATGGCGAATGGGACTCGCGACAGACATGATAAGATACATTGATGAGTTTGGACAAACCACAACTAGAATGCAGTGAAAAAAATGCTTTATTTGTGAAATTAAGCGCTGGCATTGACCCTGAG**

To overcome the toxicity of wt-SA14-14-2_**Fragment-I**, we employed an alternative ISA strategy. The initial wt-SA14-14-2_**Fragment-I** was synthesized as **two** shorter, overlapping DNA fragments (**A** and **B**) which were produced via *synthetic clonal genes* with pTwist Amp MC medium copy plasmids.

>wt-JEV-SA14_**Fragment-I-A**

**CACCCAACTGATCTTCAGCATCTTCAATATTGGCCATTAGCCATATTATTCATTGGTTATATAGCATAAATCAATATTGGCTATTGGCCATTGCATACGTTGTATCTATATCATAATATGTACATTTATATTGGCTCATGTCCAATATGACCGCCATGTTGGCATTGATTATTGACTAGTTATTAATAGTAATCAATTACGGGGTCATTAGTTCATAGCCCATATATGGAGTTCCGCGTTACATAACTTACGGTAAATGGCCCGCCTGGCTGACCGCCCAACGACCCCCGCCCATTGACGTCAATAATGACGTATGTTCCCATAGTAACGCCAATAGGGACTTTCCATTGACGTCAATGGGTGGAGTATTTACGGTAAACTGCCCACTTGGCAGTACATCAAGTGTATCATATGCCAAGTCCGCCCCCTATTGACGTCAATGACGGTAAATGGCCCGCCTGGCATTATGCCCAGTACATGACCTTACGGGACTTTCCTACTTGGCAGTACATCTACGTATTAGTCATCGCTATTACCATGGTGATGCGGTTTTGGCAGTACACCAATGGGCGTGGATAGCGGTTTGACTCACGGGGATTTCCAAGTCTCCACCCCATTGACGTCAATGGGAGTTTGTTTTGGCACCAAAATCAACGGGACTTTCCAAAATGTCGTAATAACCCCGCCCCGTTGACGCAAATGGGCGGTAGGCGTGTACGGTGGGAGGTCTATATAAGCAGAGCTCGTTTAGTGAACCG**AGAAGTTTATCTGTGTGAACTTCTTGGCTTAGTATCGTAGAGAAGAATCGAGAGATTAATGCAGTTTAAACAGTTTTTTAGAACGGAAGATAACCATGACTAAAAAACCAGGAGGGCCCGGTAAAAACCGGGCTATCAATATGCTGAAACGCGGCCTACCCCGCGTATTCCCACTAGTGGGAGTGAAGAGGGTAGTAATGAGCTTGTTGGACGGCAGAGGGCCAGTACGTTTCGTGCTGGCTCTTATCACGTTCTTCAAGTTTACAGCATTAGCCCCGACCAAGGCGCTTTCAGGCCGATGGAAAGCAGTGGAAAAGAGTGTGGCAATGAAACATCTTACTAGTTTCAAACGAGAACTTGGAACACTCATTGACGCCGTGAACAAGCGGGGCAGAAAGCAAAACAAAAGAGGAGGAAATGAAGGCTCAATCATGTGGCTCGCGAGCTTGGCAGTTGTCATAGCTTGTGCAGGAGCCATGAAGTTGTCGAATTTCCAGGGGAAGCTTTTGATGACCATTAACAACACGGACATTGCAGACGTTATCGTGATTCCCACCTCAAAAGGAGAGAACAGATGCTGGGTCCGGGCAATCGACGTCGGCTACATGTGTGAGGACACTATCACGTACGAATGTCCTAAGCTTACCATGGGCAATGATCCAGAGGATGTGGATTGCTGGTGTGACAACCAAGAAGTCTACGTCCAATATGGACGGTGCACGCGGACCAGGCATTCCAAGCGAAGCAGGAGATCCGTGTCGGTCCAAACACATGGGGAGAGTTCACTAGTGAATAAAAAAGAGGCTTGGCTGGATTCAACGAAAGCCACACGATATCTCATGAAAACTGAGAACTGGATCATAAGGAATCCTGGCTATGCTTTTCTGGCGGCGGTACTTGGCTGGATGCTTGGCAGTAACAACGGTCAACGCGTGGTATTTACCATCCTCCTGCTGTTGGTCGCTCCGGCTTACAGTTTTAATTGTCTGGGAATGGGCAATCGTGACTTCATAGAAGGAGCCAGTGGAGCCACTTGGGTGGACTTGGTGCTAGAAGGAGACAGCTGCTTGACAATCATGGCAAACAACAAACCAACATTGGACGTCCGCATGATTAACATCGAAGCTAGCCAACTTGCTGAGGTCAGAAGTTACTGCTATCATGCTTCAGTCACTGACATCTCGACGGTGGCTCGGTGCCCCACGACTGGAGAAGCCCACAACGAGAAGCGAGCTGATAGTAGCTATGTGTGCAAACAAGGCTTCACTGACCGTGGGTGGGGCAACGGATGTGGATTTTTCGGGAAGGGAAGCATTGACACATGTGCAAAATTCTCCTGCACCAGTAAAGCGATTGGGAGAACAATCCAGCCAGAAAACATCAAATACAAAGTTGGCATTTTTGTGCATGGAACCACCACTTCGGAAAACCATGGGAATTATTCAGCGCAAGTTGGGGCGTCCCAGGCGGCAAAGTTTACAGTAACACCCAATGCTCCTTCGGTAGCCCTCAAACTTGGTGACTACGGAGAAGTCACACTGGACTGTGAGCCAAGGA

>wt-JEV-SA14_**Fragment-I-B**

TTACAGTAACACCCAATGCTCCTTCGGTAGCCCTCAAACTTGGTGACTACGGAGAAGTCACACTGGACTGTGAGCCAAGGAGTGGACTGAACACTGAAGCGTTTTACGTCATGACCGTGGGGTCAAAGTCATTTCTGGTCCATAGGGAGTGGTTTCATGACCTCGCTCTCCCCTGGACGTCCCCTTCGAGCACAGCGTGGAGAAACAGAGAACTCCTCATGGAATTTGAAGGGGCGCACGCCACAAAACAGTCCGTTGTTGCTCTTGGGTCACAGGAAGGAGGCCTCCATCATGCGTTGGCAGGAGCCATCGTGGTGGAGTACTCAAGCTCAGTGATGTTAACATCAGGCCACCTGAAATGTAGGCTGAAAATGGACAAACTGGCTCTGAAAGGCACAACCTATGGCATGTGTACAGAAAAATTCTCGTTCGCGAAAAATCCGGTGGACACTGGTCACGGAACAGTTGTCATTGAACTCTCCTACTCTGGGAGTGATGGCCCCTGCAAAATTCCGATTGTTTCCGTTGCGAGCCTCAATGACATGACCCCCGTTGGGCGGCTGGTGACAGTGAACCCCTTCGTCGCGACTTCCAGTGCCAACTCAAAGGTGCTGGTCGAGATGGAACCCCCCTTCGGAGACTCCTACATCGTAGTTGGAAGGGGAGCCAAGCAGATCAACCACCATTGGCACAAAGCTGGAAGCACGCTGGGCAAGGCCTTTTCAACAACTTTGAAGGGAGCTCAAAGACTGGCAGCGTTGGGCGACACAGCCTGGGACTTTGGCTCTATTGGAGGGGTCTTCAACTCCATAGGAAGAGCCGTTCACCAAGTGTTTGGTGGTGCCTTCAGAACACTCTTTGGGGGAATGTCTTGGATCACACAAGGGCTAATGGGTGCCCTACTGCTCTGGATGGGCGTCAACGCACGAGACCGATCAATTGCTTTGGCCTTCTTAGCCACAGGAGGTGTGCTCGTGTTCTTAGCGACCAATGTGCATGCTGACACTGGATGTGCCATTGACATCACAAGAAAAGAGATGAGATGTGGAAGTGGCATCTTCGTGCACAACGACGTGGAAGCCTGGGTGGATAGGTATAAATATTTGCCAGAAACGCCCAGATCCCTAGCGAAGATCGTCCACAAAGCGCACAAGGAAGGCGTGTGCGGAGTCAGATCTGTCACTAGACTGGAGCACCAAATGTGGGAAGCCGTAAGGGACGAATTGAACGTCCTGCTCAAAGAGAATGCAGTGGACCTCAGTGTGGTTGTGAACAAGCCCGTGGGAAGATATCGCTCAGCCCCTAAACGCCTATCCATGACGCAAGAGAAGTTTGAAATGGGCTGGAAAGCATGGGGAAAAAGCATCCTCTTTGCCCCGGAATTGGCTAACTCCACATTTGTCGTAGATGGACCTGAGACAAAGGAATGCCCTGATGAGCACAGAGCTTGGAACAGCATGCAAATCGAAGACTTCGGCTTTGGCATCACATCAACCCGTGTGTGGCTGAAAATTAGAGAGGAGAGCACTGACGAGTGTGATGGAGCGATCATAGGCACGGCTGTCAAAGGACATGTGGCAGTCCATAGTGACTTGTCGTACTGGATTGAGAGTCGCTACAACGACACATGGAAACTTGAGAGGGCAGTCTTTGGAGAGGTCAAATCTTGCACTTGGCCAGAGACACACACCCTTTGGGGAGATGATGTTGAGGAAAGTGAACTCATCATTCCGCACACCATAGCCGGACCAAAAAGCAAGCACAATCGGAGGGAAGGGTATAAGACACAAAACCAGGGACCTTGGGATGAGAATGGCATAGTCTTGGACTTTGATTATTGCCCAGGGACAAAAGTCACCATTACAGAGGATTGTAGCAAGAGAGGCCCTTCGGTCAGAACCACTACTGACAGTGGAAAGTTGATCACTGACTGGTGCTGTCGCAGTTGCTCCCTTCCGCCCCTACGATTCCGGACAGAAAATGGCTGCTGGTACGGAATGGAAATCAGACCTGTTATGCATGATGAAACAACACTCGTCAGATCACAGGTTCATGCTTTCAAAGGTGAAATGGTTGACCCTTTTCAGCTGGGCCTTCTGGTGATGTTTCTGGCCACCCAGGAAGTCCTTCGCAAGAGGTGGACGGCCAGATTGACCATTCCTGCGGTTTTGGGGGTCCTACTTGTGCTGATGCTTGGGGGTATCACTTACACTGATTTGGCGAGGTATGTGGTGCTAGTCGCTGCTGCTTTCGCAGAGGCCAACAGTGGAGGAGACGTCCTGCACCTTGCTTTGATTGCTGTTTTTAAGATCCAAC

For wt-SA14-14-2_**Fragment-II** and wt-SA14-14-2_**Frgament-III** the same sequences as above were used.

**JEV enriched for CpG (JEV-CpG) ISA Fragments**

JEV strain SA14-14-2 [GenBank: MK585066.1]

Highlighted in red – pCMV promoter sequence.

Highlighted in green – HDR/SV40pA sequence.

>E-CpG-SA14-**Fragment-1A**

CACCCAACTGATCTTCAGCATCTTCAATATTGGCCATTAGCCATATTATTCATTGGTTATATAGCATAAATCAATATTGGCTATTGGCCATTGCATACGTTGTATCTATATCATAATATGTACATTTATATTGGCTCATGTCCAATATGACCGCCATGTTGGCATTGATTATTGACTAGTTATTAATAGTAATCAATTACGGGGTCATTAGTTCATAGCCCATATATGGAGTTCCGCGTTACATAACTTACGGTAAATGGCCCGCCTGGCTGACCGCCCAACGACCCCCGCCCATTGACGTCAATAATGACGTATGTTCCCATAGTAACGCCAATAGGGACTTTCCATTGACGTCAATGGGTGGAGTATTTACGGTAAACTGCCCACTTGGCAGTACATCAAGTGTATCATATGCCAAGTCCGCCCCCTATTGACGTCAATGACGGTAAATGGCCCGCCTGGCATTATGCCCAGTACATGACCTTACGGGACTTTCCTACTTGGCAGTACATCTACGTATTAGTCATCGCTATTACCATGGTGATGCGGTTTTGGCAGTACACCAATGGGCGTGGATAGCGGTTTGACTCACGGGGATTTCCAAGTCTCCACCCCATTGACGTCAATGGGAGTTTGTTTTGGCACCAAAATCAACGGGACTTTCCAAAATGTCGTAATAACCCCGCCCCGTTGACGCAAATGGGCGGTAGGCGTGTACGGTGGGAGGTCTATATAAGCAGAGCTCGTTTAGTGAACCGagaagtttatctgtgtgaacttcttggcttagtatcgtagagaagaatcgagagattaatgcagtttaaacagttttttagaacggaagataaccatgactaaaaaaccaggagggcccggtaaaaaccgggctatcaatatgctgaaacgcggcctaccccgcgtattcccactagtgggagtgaagagggtagtaatgagcttgttggacggcagagggccagtacgtttcgtgctggctcttatcacgttcttcaagtttacagcattagccccgaccaaggcgctttcaggccgatggaaagcagtggaaaagagtgtggcaatgaaacatcttactagtttcaaacgagaacttggaacactcattgacgccgtgaacaagcggggcagaaagcaaaacaaaagaggaggaaatgaaggctcaatcatgtggctcgcgagcttggcagttgtcatagcttgtgcaggagccatgaagttgtcgaatttccaggggaagcttttgatgaccattaacaacacggacattgcagacgttatcgtgattcccacctcaaaaggagagaacagatgctgggtccgggcaatcgacgtcggctacatgtgtgaggacactatcacgtacgaatgtcctaagcttaccatgggcaatgatccagaggatgtggattgctggtgtgacaaccaagaagtctacgtccaatatggacggtgcacgcggaccaggcattccaagcgaagcaggagatccgtgtcggtccaaacacatggggagagttcactagtgaataaaaaagaggcttggctggattcaacgaaagccacacgatatctcatgaaaactgagaactggatcataaggaatcctggctatgcttttctggcggcggtacttggctggatgcttggcagtaacaacggtcaacgcgtggtatttaccatcctcctgctgttggtcgctccggcttacagtTTTAATTGTCTGGGAATGGGAAATCGCGATTTCATCGAAGGCGCAAGCGGCGCAACATGGGTCGATTTAGTTCTCGAAGGCGACAGTTGCTTAACTATAATGGCGAACAACAAACCGACGTTGGACGTCCGAATGATCAACATCGAAGCGAGTCAACTCGCGGAAGTTCGAAGTTATTGCTATCACGCGTCGGTAACGGATATCTCGACGGTCGCGCGATGTCCGACGACGGGCGAAGCGCATAACGAAAAACGCGCGGACAGCAGCTACGTATGCAAACAAGGCTTCACGGATCGCGGTTGGGGAAACGGATGCGGATTTTTCGGAAAAGGAAGCATCGACACGTGCGCAAAATTTTCTTGCACGAGCAAAGCGATCGGACGAACGATCCAACCCGAAAATATTAAATACAAAGTAGGAATTTTCGTTCACGGAACAACGACGTCGGAAAATCATGGAAATTATTCCGCACAAGTCGGCGCGTCACAAGCGGCAAAATTTACCGTCACGCCGAACGCGCCGTCCGTCGCTCTCAAACTCGGCGATTATGGCGAAGTGACGCTCGACTGCGAACCTCGAA

>E-CpG-SA14-**Fragment-1B**

TTACCGTCACGCCGAACGCGCCGTCCGTCGCTCTCAAACTCGGCGATTATGGCGAAGTGACGCTCGACTGCGAACCTCGAAGCGGACTAAACACAGAAGCGTTTTATGTAATGACCGTCGGATCGAAATCGTTTCTCGTTCATCGCGAATGGTTTCACGATCTTGCGCTCCCGTGGACTTCACCGTCAAGCACCGCGTGGCGAAATCGAGAACTCCTCATGGAATTCGAAGGCGCGCACGCAACAAAACAATCCGTCGTAGCCCTTGGATCGCAAGAAGGCGGACTTCATCACGCGTTAGCGGGCGCGATTGTCGTCGAATATTCCAGCTCTGTGATGTTGACGTCGGGACATCTGAAATGTCGACTCAAAATGGACAAACTAGCGCTAAAAGGGACGACATACGGAATGTGCACGGAAAAATTTTCGTTCGCGAAAAATCCGGTCGACACGGGTCATGGAACGGTTGTAATCGAACTATCTTATTCGGGCAGCGATGGCCCTTGCAAAATTCCGATCGTTTCGGTCGCGAGTCTGAACGACATGACGCCCGTCGGTCGTCTCGTGACAGTAAATCCGTTTGTTGCGACGTCGAGCGCAAATTCAAAAGTTCTAGTCGAAATGGAACCACCCTTCGGTGATTCGTACATCGTCGTCGGACGCGGCGCGAAACAAATCAATCATCATTGGCACAAAGCGGGAAGCACGCTCGGAAAAGCGTTTTCGACGACATTGAAAGGCGCGCAACGACTCGCCGCGTTGGGCGACACGGCGTGGGATTTCGGATCGATTGGCGGTGTTTTCAACTCGATCGGACGCGCCGTTCATCAAGTGTTCGGAGGAGCGTTCCGAACACTTTTCGGCGGAATGTCTTGGATTACGCAAGGTCTTATGGGTGCTCTCCTTCTTTGGATGGGAGTCAACGCTCGCGATCGATCGATTGCGTTGGCGTTTTTGGCGACGGGTGGCGTTCTTGTATTTTTGGCGACCAATGTGCATGCTgacactggatgtgccattgacatcacaagaaaagagatgagatgtggaagtggcatcttcgtgcacaacgacgtggaagcctgggtggataggtataaatatttgccagaaacgcccagatccctagcgaagatcgtccacaaagcgcacaaggaaggcgtgtgcggagtcagatctgtcactagactggagcaccaaatgtgggaagccgtaagggacgaattgaacgtcctgctcaaagagaatgcagtggacctcagtgtggttgtgaacaagcccgtgggaagatatcgctcagcccctaaacgcctatccatgacgcaagagaagtttgaaatgggctggaaagcatggggaaaaagcatcctctttgccccggaattggctaactccacatttgtcgtagatggacctgagacaaaggaatgccctgatgagcacagagcttggaacagcatgcaaatcgaagacttcggctttggcatcacatcaacccgtgtgtggctgaaaattagagaggagagcactgacgagtgtgatggagcgatcataggcacggctgtcaaaggacatgtggcagtccatagtgacttgtcgtactggattgagagtcgctacaacgacacatggaaacttgagagggcagtctttggagaggtcaaatcttgcacttggccagagacacacaccctttggggagatgatgttgaggaaagtgaactcatcattccgcacaccatagccggaccaaaaagcaagcacaatcggagggaagggtataagacacaaaaccagggaccttgggatgagaatggcatagtcttggactttgattattgcccagggacaaaagtcaccattacagaggattgtagcaagagaggcccttcggtcagaaccactactgacagtggaaagttgatcactgactggtgctgtcgcagttgctcccttccgcccctacgattccggacagaaaatggctgctggtacggaatggaaatcagacctgttatgcatgatgaaacaacactcgtcagatcacaggttcatgctttcaaaggtgaaatggttgacccttttcagctgggccttctggtgatgtttctggccacccaggaagtccttcgcaagaggtggacggccagattgaccattcctgcggttttgggggtcctacttgtgctgatgcttgggggtatcacttacactgatttggcgaggtatgtggtgctagtcgctgctgctttcgcagaggccaacagtggaggagacgtcctgcaccttgctttgattgctgtttttaagatccaac

>wt-SA14-14-2- **Fragment-II**

gctttcgcagaggccaacagtggaggagacgtcctgcaccttgctttgattgctgtttttaagatccaaccagcatttttagtgatgaacatgcttagcacgagatggacgaaccaagaaaacgtggttctggtcctaggggctgcctttttccaattggcctcagtagatctgcaaataggagtccacggaatcctgaatgccgccgctatagcatggatgattgtccgagcgatcaccttccccacaacctcctccgtcaccatgccagtcttagcgcttctaactccggggatgagggctctatacctagacacttacagaatcatcctcctcgtcatagggatttgctccctgctgcacgagaggaaaaagaccatggcgaaaaagaaaggagctgtactcttgggcttagcgctcacatccactggatggttctcgcccaccactatagctgccggactaatggtctgcaacccaaacaagaagagagggtggccagctactgagtttttgtcggcagttggattgatgtttgccatcgtaggtggtttggccgagttggatattgaatccatgtcaatacccttcatgctggcaggtctcatggcagtgtcctacgtggtgtcaggaaaagcaacagatatgtggcttgaacgggccgccgacatcagctgggatatgggtgctgcaatcacaggaagcagtcggaggctggatgtgaaactggatgatgacggagattttcacttgattgatgatcccggtgttccatggaaggtctgggtcctgcgcatgtcttgcattggcttagccgccctcacgccttgggccatcgttcccgccgctttcggttattggctcactttaaaaacaacaaaaagagggggcgtgttttgggacacgccatccccaaaaccttgctcaaaaggagacaccactacaggagtctaccgaattatggctagagggattcttggcacttaccaggccggcgtcggagtcatgtacgagaatgttttccacacactatggcacacaactagaggagcagccattgtgagtggagaaggaaaattgacgccatactggggtagtgtgaaagaagaccgcatagcttacggaggcccatggaggttcgaccgaaaatggaatggaacagatgacgtgcaagtgatcgtggtagaaccggggaagggcgcagtaaacatccagacaaaaccaggagtgtttcggactcccttcggggaggttggggctgttagtctggattacccgcgaggaacatccggctcacccattctggattccaatggagacattataggcctatacggcaatggagttgagcttggcgatggctcatacgtcagcgccatcgtgcagggtgaccgtcaggaggaaccagtcccagaagcttacaccccaaacatgttgagaaagagacagatgactgtgctagatttgcaccctggttcagggaaaaccaggaaaattctgccacaaataattaaggacgctatccagcagcgcctaagaacagctgtgttggcaccgacgcgggtggtagcagcagaaatggcagaagttttgagagggctcccagtacgatatcaaacttcagcagtgcagagagagcaccaagggaatgaaatagtggatgtgatgtgccacgccactctgacccatagactgatgtcaccgaacagagtgcccaactacaacctatttgtcatggatgaagctcatttcaccgacccagccagtatagccgcacgaggatacattgctaccaaggtggaattaggggaggcagcagccatctttatgacagcgaccccgcctggaaccacggatccttttcctgactcaaatgccccaatccatgatttgcaagatgagataccagacagggcatggagcagtggatacgaatggatcacagaatatgcgggtaaaaccgtgtggtttgtggcgagcgtaaaaatggggaatgagattgcaatgtgcctccaaagagcggggaaaaaggtcatccaactcaaccgcaagtcctatgacacagaatacccaaaatgtaagaatggagactgggattttgtcattaccaccgacatctctgaaatgggggccaacttcggtgcgagcagggtcatcgactgtagaaagagcgtgaaacccaccatcttagaagagggagaaggcagagtcatcctcggaaacccatctcccataaccagtgcaagcgcagctcaacggaggggcagagtaggcagaaaccccaatcaagttggagatgaataccactatggaggggctaccagtgaagatgacagtaacctagcccattggacagaggcaaagatcatgttagacaacatacacatgcccaatggactggtggcccagctctatggaccagagagggaaaaggctttcacaatggatggcgaataccgtctcagaggtgaagaaaagaaaaacttcttagagctgcttaggacggctgacctcccggtgtggctggcctacaaggtggcgtccaatggcattcagtacaccgacagaaagtggtgttttgatgggccgcgtacgaatgccatactggaggacaacaccgaggtagagatagtcacccggatgggtgagaggaaaatcctcaagccgagatggcttgatgcaagagtttatgcagatcaccaggccctcaagtggttcaaagactttgcagcagggaagagatcagccgttagcttcatagaggtgctcggtcgcatgcctgagcatttcatgggaaagacgcgggaagctttagacaccatgtacttggttgcaacggctgagaaaggtgggaaagcacaccgaatggctctcgaagagctgccagatgcactggaaaccatcacacttattgtcgccattactgtgatgacaggaggattcttcctactaatgatgcagcgaaagggtatagggaagatgggtcttggagctctagtgctcacactagctaccttcttcctgtgggcggcagaggttcctggaaccaaaatagcagggaccctgctgatcgccctgctgctgatggtggttctcatcccagaaccggaaaaacagaggtcacagacagataaccaactggcggtgtttctcatctgtgtcttgaccgtggttggagtggtggcagcaaacgagtacgggatgctagaaaaaaccaaagcggatctcaagagcatgtttggcggaaagacgcaggcatcaggactgactggattgccaagcatggcactggacctgcgtccagccacagcctgggcactgtatggggggagcacagtcgtgctaacccctcttctgaagcacctgatcacgtcggaatacgtcaccacatcgctagcttcaattaactcacaagctggctcattattcgtcttgccacgaggcgtgccttttaccgacctagacttgactgttggcctcgtcttccttggctgttggggtcaagtcaccctcacaacgtttctgacagccatggttctggcgacacttcactatgggtacatgctccctggatggcaagcagaagcactcagggctgcccagagaaggacagc

>wt-SA14-14-2- **Fragment**-**III**

gacacttcactatgggtacatgctccctggatggcaagcagaagcactcagggctgcccagagaaggacagcggctggaataatgaagaatgccgttgttgacggaatggtcgccactgatgtgcctgaactggaaaggactactcctctgatgcaaaagaaagtcggacaggtgctcctcataggggtaagcgtggcagcgttcctcgtcaaccctaatgtcaccactgtgagagaagcaggggtgttggtgacggcggctacgcttactttgtgggacaatggagccagtgccgtttggaattccaccacagccacgggactctgccatgtcatgcgaggtagctacctggctggaggctccattgcttggactctcatcaagaacgctgataagccctccttgaaaaggggaaggcctgggggcaggacgctaggggagcagtggaaggaaaaactaaatgccatgagtagagaagagttttttaaataccggagagaggccataatcgaggtggaccgcactgaagcacgcagggccagacgtgaaaataacatagtgggaggacatccggtttcgcgaggctcagcaaaactccgttggctcgtggagaaaggatttgtctcgccaataggaaaagtcattgatctagggtgtgggcgtggaggatggagctactacgcagcaaccctgaagaaggtccaggaagtcagaggatacacgaaaggtggggcgggacatgaagaaccgatgctcatgcagagctacggctggaacctggtctccctgaagagtggagtggacgtgttttacaaaccttcagagcccagtgataccctgttctgtgacataggggaatcctccccaagtccagaagtagaagaacaacgcacactacgcgtcctagagatgacatctgactggttgcaccgaggacctagagagttctgcattaaagttctctgcccttacatgcccaaggttatagaaaaaatggaagttctgcagcgtcgcttcggaggtgggctagtgcgtctccccctgtcccgaaactccaatcacgagatgtattgggttagtggagccgctggcaatgtggtgcacgctgtgaacatgaccagccaggtattactggggcgaatggatcgcacagtgtggagagggccaaagtatgaggaagatgtcaacctagggagcggaacaagagccgtgggaaagggagaagtccatagcaatcaggagaaaatcaagaagagaatccagaagcttaaagaagaattcgccacaacgtggcacaaagaccctgagcatccataccgcacttggacataccacggaagctatgaagtgaaggctactggctcagccagctctctcgtcaacggagtggtgaagctcatgagcaaaccttgggacgccattgccaacgtcaccaccatggccatgactgacaccaccccttttggacagcaaagagttttcaaggagaaagttgacacgaaggctcctgagccaccagctggagccaaggaagtgctcaacgagaccaccaactggctgtgggcctacttgtcacgggaaaaaagaccccgcttgtgcaccaaggaagaattcattaagaaagttaacagcaacgcggctcttggagcagtgttcgctgaacagaatcaatggagcacggcgcgtgaggctgtggatgacccgcggttttgggagatggttgatgaagagagggaaaaccatctgcgaggagagtgtcacacatgtatctacaacatgatgggaaaaagagagaagaagcctggagagtttggaaaagctaaaggaagcagggccatttggttcatgtggcttggagcacggtatctagagtttgaagctttggggttcctgaatgaagaccattggctgagccgagagaattcaggaggtggagtggaaggctcaggcgtccaaaagctgggatacatcctccgtgacatagcaggaaagcaaggagggaaaatgtacgctgatgataccgccgggtgggacactagaattaccagaactgatttagaaaatgaagctaaggtactggagctcctagacggtgaacaccgcatgctcgcccgagccataattgaactgacttacaggcacaaagtggtcaaggtcatgagacctgcagcagaaggaaagaccgtgatggacgtgatatcaagagaagatcaaagggggagtggacaggtggtcacttatgctcttaacactttcacgaacatcgctgtccagctcgtcaggctgatggaggctgagggggtcattggaccacaacacttggaacatctacctaggaaaaacaagatagctgtcaggacctggctctttgagaatggagaggagagagtgaccaggatggcgatcagcggagacgactgtgccgtcaaaccgctggacgacagattcgccacagccctccacttcctcaacgcaatgtcaaaggtcagaaaagacatccaggaatggaagccttcgcatggctggcacgattggcagcaagttcccttctgttctaaccattttcaggagattgtgatgaaagatggaaggagtatagttgtcccgtgcagaggacaggatgagctgataggcagggctcgcatctctccaggagctggatggaatgtgaaggacacagcttgcctggccaaagcatatgcacagatgtggctactcctatacttccatcgcagggacttgcgtctcatggcaaatgcgatttgctcagcagtgccagtagattgggtgcccacaggcaggacatcctggtcaatacactcgaaaggagagtggatgaccacggaagacatgctgcaggtctggaacagagtttggattgaagaaaatgaatggatgatggacaagactccaatcacaagctggacagacgttccgtatgtgggaaagcgcgaggacatctggtgtggcagcctcatcggaacgcgatccagagcaacctgggctgagaacatctatgcggcgataaaccaggttagagctgtcattgggaaagaaaattatgttgactacatgacctcactcaggagatacgaagacgtcttgatccaggaagacagggtcatctagtgtgatttaaggtagaaaagtagactatgtaaacaatgtaaatgagaaaatgcatgcatatggagtcaggccagcaaaagctgccaccggatactgggtagacggtgctgcctgcgtctcagtcccaggaggactgggttaacaaatctgacaacagaaagtgagaaagccctcagaaccgtctcggaagtaggtccctgctcactggaagttgaaagaccaacgtcaggccacaaatttgtgccactccgctagggagtgcggcctgcgcagccccaggaggactgggttaccaaagccgttgaggcccccacggcccaagcctcgtctaggatgcaatagacgaggtgtaaggactagaggttagaggagaccccgtggaaacaacaacatgcggcccaagccccctcgaagctgtagaggaggtggaaggactagaggttagaggagaccccgcatttgcatcaaacagcatattgacacctgggaatagactgggagatcttctgctctatctcaacatcagctactaggcacagagcgccgaagtatgtagctggtggtgaggaagaacacaggatctGGCCGGCATGGTCCCAGCCTCCTCGCTGGCGCCGGCTGGGCAACATTCCGAGGGGACCGTCCCCTCGGTAATGGCGAATGGGACTCGCGACAGACATGATAAGATACATTGATGAGTTTGGACAAACCACAACTAGAATGCAGTGAAAAAAATGCTTTATTTGTGAAATTAAGCGCTGGCATTGACCCTGAG

**WNV wild-type (WNV-WT) ISA Fragments**

Parental strain – WNV strain NY99 [GenBank: DQ211652.1].

Highlighted in red – pCMV promoter sequence.

Highlighted in green – HDR/SV40pA sequence.

>wt-NY99_**Fragment-I**

During the rescue of the wild-type WNV NY99 strain (WNV-WT), Fragment I exhibited bacterial toxicity when using a medium copy pET-28a(+) bacterial plasmid. To address this, a low copy plasmid pCC1was utilized, which enabled the successful rescue of Fragment I after several attempts over a three-month period.

**CACCCAACTGATCTTCAGCATCTTCAATATTGGCCATTAGCCATATTATTCATTGGTTATATAGCATAAATCAATATTGGCTATTGGCCATTGCATACGTTGTATCTATATCATAATATGTACATTTATATTGGCTCATGTCCAATATGACCGCCATGTTGGCATTGATTATTGACTAGTTATTAATAGTAATCAATTACGGGGTCATTAGTTCATAGCCCATATATGGAGTTCCGCGTTACATAACTTACGGTAAATGGCCCGCCTGGCTGACCGCCCAACGACCCCCGCCCATTGACGTCAATAATGACGTATGTTCCCATAGTAACGCCAATAGGGACTTTCCATTGACGTCAATGGGTGGAGTATTTACGGTAAACTGCCCACTTGGCAGTACATCAAGTGTATCATATGCCAAGTCCGCCCCCTATTGACGTCAATGACGGTAAATGGCCCGCCTGGCATTATGCCCAGTACATGACCTTACGGGACTTTCCTACTTGGCAGTACATCTACGTATTAGTCATCGCTATTACCATGGTGATGCGGTTTTGGCAGTACACCAATGGGCGTGGATAGCGGTTTGACTCACGGGGATTTCCAAGTCTCCACCCCATTGACGTCAATGGGAGTTTGTTTTGGCACCAAAATCAACGGGACTTTCCAAAATGTCGTAATAACCCCGCCCCGTTGACGCAAATGGGCGGTAGGCGTGTACGGTGGGAGGTCTATATAAGCAGAGCTCGTTTAGTGAACCG**AGTAGTTCGCCTGTGTGAGCTGACAAACTTAGTAGTGTTTGTGAGGATTAACAACAATTAACACAGTGCGAGCTGTTTCTTAGCACGAAGATCTCGATGTCTAAGAAACCAGGAGGGCCCGGCAAGAGCCGGGCTGTCAATATGCTAAAACGCGGAATGCCCCGCGTGTTGTCCTTGATTGGACTGAAGAGGGCTATGTTGAGCCTGATCGACGGCAAGGGGCCAATACGATTTGTGTTGGCTCTCTTGGCGTTCTTCAGGTTCACAGCAATTGCTCCGACCCGAGCAGTGCTGGATCGATGGAGAGGTGTGAACAAACAAACAGCGATGAAACACCTTCTGAGTTTTAAGAAGGAACTAGGGACCTTGACCAGTGCTATCAATCGGCGGAGCTCAAAACAAAAGAAAAGAGGAGGAAAGACCGGAATTGCAGTCATGATTGGCCTGATCGCCAGCGTAGGAGCAGTTACCCTCTCTAACTTCCAAGGGAAGGTGATGATGACGGTAAATGCTACTGACGTCACAGATGTCATCACGATTCCAACAGCTGCTGGAAAGAACCTATGCATTGTCAGAGCAATGGATGTGGGATACATGTGCGATGATACTATCACTTATGAATGCCCAGTACTGTCGGCTGGTAATGATCCAGAAGACATCGACTGTTGGTGCACAAAGTCAGCAGTCTACGTCAGGTATGGAAGATGCACCAAGACACGCCACTCAAGACGCAGTCGGAGGTCACTGACAGTGCAGACACACGGAGAAAGCACTCTAGCGAACAAGAAGGGGGCTTGGATGGACAGCACCAAGGCCACAAGGTATTTGGTAAAAACAGAATCATGGATCTTGAGGAACCCTGGATATGCCCTGGTGGCAGCCGTCATTGGTTGGATGCTTGGGAGCAACACCATGCAGAGAGTTGTGTTTGTCGTGCTATTGCTTTTGGTGGCCCCAGCTTACAGCTTCAACTGCCTTGGAATGAGCAACAGAGACTTCTTGGAAGGAGTGTCTGGAGCAACATGGGTGGATTTGGTTCTCGAAGGCGACAGCTGCGTGACTATCATGTCTAAGGACAAGCCTACCATCGATGTGAAGATGATGAATATGGAGGCGGCCAACCTGGCAGAGGTCCGCAGTTATTGCTATTTGGCTACCGTCAGCGATCTCTCCACCAAAGCTGCGTGCCCGACCATGGGAGAAGCTCACAATGACAAACGTGCTGACCCAGCTTTTGTGTGCAGACAAGGAGTGGTGGACAGGGGCTGGGGCAACGGCTGCGGACTATTTGGCAAAGGAAGCATTGACACATGCGCCAAATTTGCCTGCTCTACCAAGGCAATAGGAAGAACCATCTTGAAAGAGAATATCAAGTACGAAGTGGCCATTTTTGTCCATGGACCAACTACTGTGGAGTCGCACGGAAACTACTCCACACAGGTTGGAGCCACTCAGGCAGGGAGACTCAGCATCACTCCTGCGGCGCCTTCATACACACTAAAGCTTGGAGAATATGGAGAGGTGACAGTGGACTGTGAACCACGGTCAGGGATTGACACCAATGCATACTACGTGATGACTGTTGGAACAAAGACGTTCTTGGTCCATCGTGAGTGGTTCATGGACCTCAACCTCCCTTGGAGCAGTGCTGGAAGTACTGTGTGGAGGAACAGAGAGACGTTAATGGAGTTTGAGGAACCACACGCCACGAAGCAGTCTGTGATAGCATTGGGCTCACAAGAGGGAGCTCTGCATCAAGCTTTGGCTGGAGCCATTCCTGTGGAATTTTCAAGCAACACTGTCAAGTTGACGTCGGGTCATTTGAAGTGTAGAGTGAAGATGGAAAAATTGCAGTTGAAGGGAACAACCTATGGCGTCTGTTCAAAGGCTTTCAAGTTTCTTGGGACTCCCGCAGACACAGGTCACGGCACTGTGGTGTTGGAATTGCAGTACACTGGCACGGATGGACCTTGCAAAGTTCCTATCTCGTCAGTGGCTTCATTGAACGACCTAACGCCAGTGGGCAGATTGGTCACTGTCAACCCTTTTGTTTCAGTGGCCACGGCCAACGCTAAGGTCCTGATTGAATTGGAACCACCCTTTGGAGACTCATACATAGTGGTGGGCAGAGGAGAACAACAGATCAATCACCATTGGCACAAGTCTGGAAGCAGCATTGGCAAAGCCTTTACAACCACCCTCAAAGGAGCGCAGAGACTAGCCGCTCTAGGAGACACAGCTTGGGACTTTGGATCAGTTGGAGGGGTGTTCACCTCAGTTGGGAAGGCTGTCCATCAAGTGTTCGGAGGAGCATTCCGCTCACTGTTCGGAGGCATGTCCTGGATAACGCAAGGATTGCTGGGGGCTCTCCTGTTGTGGATGGGCATCAATGCTCGTGATAGGTCCATAGCTCTCACGTTTCTCGCAGTTGGAGGAGTTCTGCTCTTCCTCTCCGTGAACGTGCACGCTGACACTGGGTGTGCCATAGACATCAGCCGGCAAGAGCTGAGATGTGGAAGTGGAGTGTTCATACACAATGATGTGGAGGCTTGGATGGACCGGTACAAGTATTACCCTGAAACGCCACAAGGCCTAGCCAAGATCATTCAGAAAGCTCATAAGGAAGGAGTGTGCGGTCTACGATCAGTTTCCAGACTGGAGCATCAAATGTGGGAAGCAGTGAAGGACGAGCTGAACACTCTTTTGAAGGAGAATGGTGTGGACCTTAGTGTCGTGGTTGAGAAACAGGAGGGAATGTACAAGTCAGCACCTAAACGCCTCACCGCCACCACGGAAAAATTGGAAATTGGCTGGAAGGCCTGGGGAAAGAGTATTTTATTTGCACCAGAACTCGCCAACAACACCTTTGTGGTTGATGGTCCGGAGACCAAGGAATGTCCGACTCAGAATCGCGCTTGGAATAGCTTAGAAGTGGAGGATTTTGGATTTGGTCTCACCAGCACTCGGATGTTCCTGAAGGTCAGAGAGAGCAACACAACTGAATGTGACTCGAAGATCATTGGAACGGCTGTCAAGAACAACTTGGCGATCCACAGTGACCTGTCCTATTGGATTGAAAGCAGGCTCAATGATACGTGGAAGCTTGAAAGGGCAGTTCTGGGTGAAGTCAAATCATGTACGTGGCCTGAGACGCATACCTTGTGGGGCGATGGAATCCTTGAGAGTGACTTGATAATACCAGTCACACTGGCGGGACCACGAAGCAATCACAATCGGAGACCTGGGTACAAGACACAAAACCAGGGCCCATGGGACGAAGGCCGGGTAGAGATTGACTTCGATTACTGCCCAGGAACTACGGTCACCCTGAGTGAGAGCTGCGGACACCGTGGACCTGCCACTCGCACCACCACAGAGAGCGGAAAGTTGATAACAGATTGGTGCTGCAGGAGCTGCACCTTACCACCACTGCGCTACCAAACTGACAGCGGCTGTTGGTATGGTATGGAGATCAGACCACAGAGACATGATGAAAAGACCCTCGTGCAGTCACAAGTGAATGCTTATAATGCTGATATGATTGACCCTTTTCAGTTGGGCCTTCTGGTCGTGTTCTTGGCCACCCAGGAGGTCCTTCGC

>wt-NY99_**Fragment-II**

TATAATGCTGATATGATTGACCCTTTTCAGTTGGGCCTTCTGGTCGTGTTCTTGGCCACCCAGGAGGTCCTTCGCAAGAGGTGGACAGCCAAGATCAGCATGCCAGCTATACTGATTGCTCTGCTAGTCCTGGTGTTTGGGGGCATTACTTACACTGATGTGTTACGCTATGTCATCTTGGTGGGGGCAGCTTTCGCAGAATCTAATTCGGGAGGAGACGTGGTACACTTGGCGCTCATGGCGACCTTCAAGATACAACCAGTGTTTATGGTGGCATCGTTTCTCAAAGCGAGATGGACCAACCAGGAGAACATTTTGTTGATGTTGGCGGCTGTTTTCTTTCAAATGGCTTATCACGATGCCCGCCAAATTCTGCTCTGGGAGATCCCTGATGTGTTGAATTCACTGGCGGTAGCTTGGATGATACTGAGAGCCATAACATTCACAACGACATCAAACGTGGTTGTTCCGCTGCTAGCCCTGCTAACACCCGGGCTGAGATGCTTGAATCTGGATGTGTACAGGATACTGCTGTTGATGGTCGGAATAGGCAGCTTGATCAGGGAGAAGAGGAGTGCAGCCGCAAAAAAGAAAGGAGCAAGTCTGCTATGCTTGGCTCTAGCCTCAACAGGACTTTTCAACCCCATGATCCTTGCTGCTGGACTGATTGCATGTGATCCCAACCGTAAACGCGGATGGCCCGCAACTGAAGTGATGACAGCTGTCGGCCTAATGTTTGCCATCGTCGGAGGGCTGGCAGAGCTTGACATTGACTCCATGGCCATTCCAATGACTATCGCGGGGCTCATGTTTGCTGCTTTCGTGATTTCTGGGAAATCAACAGATATGTGGATTGAGAGAACGGCGGACATTTCCTGGGAAAGTGATGCAGAAATTACAGGCTCGAGCGAAAGAGTTGATGTGCGGCTTGATGATGATGGAAACTTCCAGCTCATGAATGATCCAGGAGCACCTTGGAAGATATGGATGCTCAGAATGGTCTGTCTCGCGATTAGTGCGTACACCCCCTGGGCAATCTTGCCCTCAGTAGTTGGATTTTGGATAACTCTCCAATACACAAAGAGAGGAGGCGTGTTGTGGGACACTCCCTCACCAAAGGAGTACAAAAAGGGGGACACGACCACCGGCGTCTACAGGATCATGACTCGTGGGCTGCTCGGCAGTTATCAAGCAGGAGCGGGCGTGATGGTTGAAGGTGTTTTCCACACCCTTTGGCATACAACAAAAGGAGCCGCTTTGATGAGCGGAGAGGGCCGCCTGGACCCATACTGGGGCAGTGTCAAGGAGGATCGACTTTGTTACGGAGGACCCTGGAAATTGCAGCACAAGTGGAACGGGCAGGATGAGGTGCAGATGATTGTGGTGGAACCTGGCAAGAACGTTAAGAACGTCCAGACGAAACCAGGGGTGTTCAAAACACCTGAAGGAGAAATCGGGGCCGTGACTTTGGACTTCCCCACTGGAACATCAGGCTCACCAATAGTGGACAAAAACGGTGATGTGATTGGGCTTTATGGCAATGGAGTCATAATGCCCAACGGCTCATACATAAGCGCGATAGTGCAGGGTGAAAGGATGGATGAGCCAATCCCAGCCGGATTCGAACCTGAGATGCTGAGGAAAAAACAGATCACTGTACTGGATCTCCATCCCGGCGCCGGTAAAACAAGGAGGATTCTGCCACAGATCATCAAAGAGGCCATAAACAGAAGACTGAGAACAGCCGTGCTAGCGCCAACCAGGGTTGTGGCTGCTGAGATGGCTGAAGCACTGAGAGGACTGCCCATCCGGTACCAGACATCCGCAGTGCCCAGAGAACATAATGGAAATGAGATTGTTGATGTCATGTGTCATGCTACCCTCACCCACAGGCTGATGTCTCCTCACAGGGTGCCGAACTACAACCTGTTCGTGATGGATGAGGCTCATTTCACCGACCCAGCTAGCATTGCAGCAAGAGGTTACATTTCCACAAAGGTCGAGCTAGGGGAGGCGGCGGCAATATTCATGACAGCCACCCCACCAGGCACTTCAGATCCATTCCCAGAGTCCAATTCACCAATTTCCGACTTACAGACTGAGATCCCGGATCGAGCTTGGAACTCTGGATACGAATGGATCACAGAATACACCGGGAAGACGGTTTGGTTTGTGCCTAGTGTCAAGATGGGGAATGAGATTGCCCTTTGCCTACAACGTGCTGGAAAGAAAGTAGTCCAATTGAACAGAAAGTCGTACGAGACGGAGTACCCAAAATGTAAGAACGATGATTGGGACTTTGTTATCACAACAGACATATCTGAAATGGGGGCTAACTTCAAGGCGAGCAGGGTGATTGACAGCCGGAAGAGTGTGAAACCAACCATCATAACAGAAGGAGAAGGGAGAGTGATCCTGGGAGAACCATCTGCAGTGACAGCAGCTAGTGCCGCCCAGAGACGTGGACGTATCGGTAGAAATCCGTCGCAAGTTGGTGATGAGTACTGTTATGGGGGGCACACGAATGAAGACGACTCGAACTTCGCCCATTGGACTGAGGCACGAATCATGCTGGACAACATCAACATGCCAAACGGACTGATCGCTCAATTCTACCAACCAGAGCGTGAGAAGGTATATACCATGGATGGGGAATACCGGCTCAGAGGAGAAGAGAGAAAAAACTTTCTGGAACTGTTGAGGACTGCAGATCTGCCAGTTTGGCTGGCTTACAAGGTTGCAGCGGCTGGAGTGTCATACCACGACCGGAGGTGGTGCTTTGATGGTCCTAGGACAAACACAATTTTAGAAGACAACAACGAAGTGGAAGTCATCACGAAGCTTGGTGAAAGGAAGATTCTGAGGCCGCGCTGGATTGATGCCAGGGTGTACTCGGATCACCAGGCACTAAAGGCGTTCAAGGACTTCGCCTCGGGAAAACGTTCTCAGATAGGGCTCATTGAGGTTCTGGGAAAGATGCCTGAGCACTTCATGGGGAAGACATGGGAAGCACTTGACACCATGTACGTTGTGGCCACTGCAGAGAAAGGAGGAAGAGCTCACAGAATGGCCCTGGAGGAACTGCCAGATGCTCTTCAGACAATTGCCTTGATTGCCTTATTGAGTGTGATGACCATGGGAGTATTCTTCCTCCTCATGCAGCGGAAGGGCATTGGAAAGATAGGTTTGGGAGGCGCTGTCTTGGGAGTCGCGACCTTTTTCTGTTGGATGGCTGAAGTTCCAGGAACGAAGATCGCCGGAATGTTGCTGCTCTCCCTTCTCTTGATGATTGTGCTAATTCCTGAGCCAGAGAAGCAACGTTCGCAGACAGACAACCAGCTAGCCGTGTTCCTGATTTGTGTCATGACCCTTGTGAGCGCAGTGGCAGCCAACGAGATGGGTTGGCTAGATAAGACCAAGAGTGACATAAGCAGTTTGTTTGGGCAAAGAATTGAGGTCAAGGAGAATTTCAGCATGGGAGAGTTTCTTCTGGACTTGAGGCCGGCAACAGCCTGGTCACTGTACGCTGTGACAACAGCGGTCCTCACTCCACTGCTAAAGCATTTGATCACGTCAGATTACATCAACACCTCATTGACCTCAATAAACGTTCAGGCAAGTGCACTATTCACACTCGCGCGAGGCTTCCCCTTCGTCGATGTTGGAGTGTCGGCTCTCCTGCTAGCAGCCGGATGCTGGGGACAAGTCACCCTCACCGTTACGGTAACAGCGGCAACACTCCTTTTTTGCCACTATGCCTACATGGTTCCCGGTTGGCAAGCTGAGGCAATGCGCTCAGCCCAGCGGCGGACAGCGGCCGGAATCATGAAGAACGCTGTAGTGGATGGCATCGTGGCCACGGACGTCCCAGAATTAGAGCGCACCACACCCATCATGCAGAAGAAAGTTGGACAGATCATGCTGATCTTGGTGTCTCTAGCTGCAGTAGTAGTGAACCCGTCTGTGAAGACAGTACGAGAAGCCGGAATTTTGATCACGGCCGCAGCGGTGACGCTTTGGGAGAATGGAGCAAGCTCTGTTTGGAACGCAACAACTGCCATCGGACTCTGCCACATCATGCGTGGGGGTTGGTTGTCATGTCTATCCATAACATGGACACTCATAAAGAACATGGAAAAACCAGGACTAAAAAGAGGTGGGGCAAAAGGACGCACCTTGGGAGAGGTTTGGAAAGAAAGACTCAACCAGATGACAAAAGAAGAGTTCACT

>wt-NY99_**Fragment-III**

GGTGGGGCAAAAGGACGCACCTTGGGAGAGGTTTGGAAAGAAAGACTCAACCAGATGACAAAAGAAGAGTTCACTAGGTACCGCAAAGAGGCCATCATCGAAGTCGATCGCTCAGCGGCAAAACACGCCAGGAAAGAAGGCAATGTCACTGGAGGGCATCCAGTCTCTAGGGGCACAGCAAAACTGAGATGGCTGGTCGAACGGAGGTTTCTCGAACCGGTCGGAAAAGTGATTGACCTTGGATGTGGAAGAGGCGGTTGGTGTTACTATATGGCAACCCAAAAAAGAGTCCAAGAAGTCAGAGGGTACACAAAGGGCGGTCCCGGACATGAAGAGCCCCAACTAGTGCAAAGTTATGGATGGAACATTGTCACCATGAAGAGTGGAGTGGATGTGTTCTACAGACCTTCTGAGTGTTGTGACACCCTCCTTTGTGACATCGGAGAGTCCTCGTCAAGTGCTGAGGTTGAAGAGCATAGGACGATTCGGGTCCTTGAAATGGTTGAGGACTGGCTGCACCGAGGGCCAAGGGAATTTTGCGTGAAGGTGCTCTGTCCCTACATGCCGAAAGTCATAGAGAAGATGGAGCTGCTCCAACGCCGGTATGGGGGGGGACTGGTCAGAAACCCACTCTCACGGAATTCCACGCACGAGATGTATTGGGTGAGTCGAGCTTCAGGCAATGTGGTACATTCAGTGAATATGACCAGCCAGGTGCTCCTAGGAAGAATGGAAAAAAGGACCTGGAAGGGACCCCAATACGAGGAAGATGTAAACTTGGGAAGTGGAACCAGGGCGGTGGGAAAACCCCTGCTCAACTCAGACACCAGTAAAATCAAGAACAGGATTGAACGACTCAGGCGTGAGTACAGTTCGACGTGGCACCACGATGAGAACCACCCATATAGAACCTGGAACTATCACGGCAGTTATGATGTGAAGCCCACAGGCTCCGCCAGTTCGCTGGTCAATGGAGTGGTCAGGCTCCTCTCAAAACCATGGGACACCATCACGAATGTTACCACCATGGCCATGACTGACACTACTCCCTTCGGGCAGCAGCGAGTGTTCAAAGAGAAGGTGGACACGAAAGCTCCTGAACCGCCAGAAGGAGTGAAGTACGTGCTCAACGAGACCACCAACTGGTTGTGGGCGTTTTTGGCCAGAGAAAAACGTCCCAGAATGTGCTCTCGAGAGGAATTCATAAGAAAGGTCAACAGCAATGCAGCTTTGGGTGCCATGTTTGAAGAGCAGAATCAATGGAGGAGCGCCAGAGAAGCAGTTGAAGATCCAAAATTTTGGGAGATGGTGGATGAGGAGCGCGAGGCACATCTGCGGGGGGAATGTCACACTTGCATTTACAACATGATGGGAAAGAGAGAGAAAAAACCCGGAGAGTTCGGAAAGGCCAAGGGAAGCAGAGCCATTTGGTTCATGTGGCTCGGAGCTCGCTTTCTGGAGTTCGAGGCTCTGGGTTTTCTCAATGAAGACCACTGGCTTGGAAGAAAGAACTCAGGAGGAGGTGTCGAGGGCTTGGGCCTCCAAAAACTGGGTTACATCCTGCGTGAAGTTGGCACCCGGCCTGGGGGCAAGATCTATGCTGATGACACAGCTGGCTGGGACACCCGCATCACGAGAGCTGACTTGGAAAATGAAGCTAAGGTGCTTGAGCTGCTTGATGGGGAACATCGGCGTCTTGCCAGGGCCATCATTGAGCTCACCTATCGTCACAAAGTTGTGAAAGTGATGCGCCCGGCTGCTGATGGAAGAACCGTCATGGATGTTATCTCCAGAGAAGATCAGAGGGGGAGTGGACAAGTTGTCACCTACGCCCTAAACACTTTCACCAACCTGGCCGTCCAGCTGGTGAGGATGATGGAAGGGGAAGGAGTGATTGGCCCAGATGATGTGGAGAAACTCACAAAAGGGAAAGGACCCAAAGTCAGGACCTGGCTGTTTGAGAATGGGGAAGAAAGACTCAGCCGCATGGCTGTCAGTGGAGATGACTGTGTGGTAAAGCCCCTGGACGATCGCTTTGCCACCTCGCTCCACTTCCTCAATGCTATGTCAAAGGTTCGCAAAGACATCCAAGAGTGGAAACCGTCAACTGGATGGTATGATTGGCAGCAGGTTCCATTTTGCTCAAACCATTTCACTGAATTGATCATGAAAGATGGAAGAACACTGGTGGTTCCATGCCGAGGACAGGATGAATTGGTAGGCAGAGCTCGCATATCTCCAGGGGCCGGATGGAACGTCCGCGACACTGCTTGTCTGGCTAAGTCTTATGCCCAGATGTGGCTGCTTCTGTACTTCCACAGAAGAGACCTGCGGCTCATGGCCAACGCCATTTGCTCCGCTGTCCCTGTGAATTGGGTCCCTACCGGAAGAACCACGTGGTCCATCCATGCAGGAGGAGAGTGGATGACAACAGAGGACATGTTGGAGGTCTGGAACCGTGTTTGGATAGAGGAGAATGAATGGATGGAAGACAAAACCCCAGTGGAGAAATGGAGTGACGTCCCATATTCAGGAAAACGAGAGGACATCTGGTGTGGCAGCCTGATTGGCACAAGAGCCCGAGCCACGTGGGCAGAAAACATCCAGGTGGCTATCAACCAAGTCAGAGCAATCATCGGAGATGAGAAGTATGTGGACTACATGAGTTCACTAAAGAGATATGAAGACACAACTTTGGTTGAGGACACAGTACTGTAGATATTTAATCAATTGTAAATAGACAATATAAGTATGCATAAAAGTGTAGTTTTATAGTAGTATTTAGTGGTGTTAGTGTAAATAGTTAAGAAAATTTTGAGGAGAAAGTCAGGCCGGGAAGTTCCCGCCACCGGAAGTTGAGTAGACGGTGCTGCCTGCGACTCAACCCCAGGAGGACTGGGTGAACAAAGCCGCGAAGTGATCCATGTAAGCCCTCAGAACCGTCTCGGAAGGAGGACCCCACATGTTGTAACTTCAAAGCCCAATGTCAGACCACGCTACGGCGTGCTACTCTGCGGAGAGTGCAGTCTGCGATAGTGCCCCAGGAGGACTGGGTTAACAAAGGCAAACCAACGCCCCACGCGGCCCTAGCCCCGGTAATGGTGTTAACCAGGGCGAAAGGACTAGAGGTTAGAGGAGACCCCGCGGTTTAAAGTGCACGGCCCAGCCTGGCTGAAGCTGTAGGTCAGGGGAAGGACTAGAGGTTAGTGGAGACCCCGTGCCACAAAACACCACAACAAAACAGCATATTGACACCTGGGATAGACTAGGAGATCTTCTGCTCTGCACAACCAGCCACACGGCACAGTGCGCCGACAATGGTGGCTGGTGGTGCGAGAACACAGGATCT**GGCCGGCATGGTCCCAGCCTCCTCGCTGGCGCCGGCTGGGCAACATTCCGAGGGGACCGTCCCCTCGGTAATGGCGAATGGGACTCGCGACAGACATGATAAGATACATTGATGAGTTTGGACAAACCACAACTAGAATGCAGTGAAAAAAATGCTTTATTTGTGAAATTAAGCGCTGGCATTGACCCTGAG**

**WNV permuted (WNV-Per) ISA Fragments**

Parental strain – WNV strain NY99 [GenBank: DQ211652.1].

Highlighted in red – pCMV promoter sequence.

Highlighted in green – HDR/SV40pA sequence.

>NY99-Pr-E-NS1_**Fragment-I**

The rescue of this fragment failed due to its toxicity to bacteria. The alternative ISA strategy is described below.

CACCCAACTGATCTTCAGCATCTTCAATATTGGCCATTAGCCATATTATTCATTGGTTATATAGCATAAATCAATATTGGCTATTGGCCATTGCATACGTTGTATCTATATCATAATATGTACATTTATATTGGCTCATGTCCAATATGACCGCCATGTTGGCATTGATTATTGACTAGTTATTAATAGTAATCAATTACGGGGTCATTAGTTCATAGCCCATATATGGAGTTCCGCGTTACATAACTTACGGTAAATGGCCCGCCTGGCTGACCGCCCAACGACCCCCGCCCATTGACGTCAATAATGACGTATGTTCCCATAGTAACGCCAATAGGGACTTTCCATTGACGTCAATGGGTGGAGTATTTACGGTAAACTGCCCACTTGGCAGTACATCAAGTGTATCATATGCCAAGTCCGCCCCCTATTGACGTCAATGACGGTAAATGGCCCGCCTGGCATTATGCCCAGTACATGACCTTACGGGACTTTCCTACTTGGCAGTACATCTACGTATTAGTCATCGCTATTACCATGGTGATGCGGTTTTGGCAGTACACCAATGGGCGTGGATAGCGGTTTGACTCACGGGGATTTCCAAGTCTCCACCCCATTGACGTCAATGGGAGTTTGTTTTGGCACCAAAATCAACGGGACTTTCCAAAATGTCGTAATAACCCCGCCCCGTTGACGCAAATGGGCGGTAGGCGTGTACGGTGGGAGGTCTATATAAGCAGAGCTCGTTTAGTGAACCGAGTAGTTCGCCTGTGTGAGCTGACAAACTTAGTAGTGTTTGTGAGGATTAACAACAATTAACACAGTGCGAGCTGTTTCTTAGCACGAAGATCTCGATGTCTAAGAAACCAGGAGGGCCCGGCAAGAGCCGGGCTGTCAATATGCTAAAACGCGGAATGCCCCGCGTGTTGTCCTTGATTGGACTGAAGAGGGCTATGTTGAGCCTGATCGACGGCAAGGGGCCAATACGATTTGTGTTGGCTCTCTTGGCGTTCTTCAGGTTCACAGCAATTGCTCCGACCCGAGCAGTGCTGGATCGATGGAGAGGTGTGAACAAACAAACAGCGATGAAACACCTTCTGAGTTTTAAGAAGGAACTAGGGACCTTGACCAGTGCTATCAATCGGCGGAGCTCAAAACAAAAGAAAAGAGGAGGAAAGACCGGAATTGCAGTCATGATTGGCCTGATCGCCAGCGTAGGAGCAGTTACCCTCTCTAACTTCCAAGGGAAGGTGATGATGACGGTAAATGCTACTGACGTCACAGATGTCATCACGATTCCAACAGCTGCTGGAAAGAACCTATGCATTGTCAGAGCAATGGATGTGGGATACATGTGCGATGATACTATCACTTATGAATGCCCAGTACTGTCGGCTGGTAATGATCCAGAAGACATCGACTGTTGGTGCACAAAGTCAGCAGTCTACGTCAGGTATGGAAGATGCACCAAGACACGCCACTCAAGACGCAGTCGGAGGTCACTGACAGTGCAGACACACGGAGAAAGCACTCTAGCGAACAAGAAGGGGGCTTGGATGGACAGCACCAAGGCCACAAGGTATTTGGTAAAAACAGAATCATGGATCTTGAGGAACCCTGGATATGCCCTGGTGGCAGCCGTCATTGGTTGGATGCTTGGGAGCAACACCATGCAGAGAGTTGTGTTTGTCGTGCTATTGCTTTTGGTGGCCCCAGCTTACAGCTTCAACTGTCTGGGCATGAGCAATAGAGACTTTTTGGAAGGAGTCTCCGGAGCAACCTGGGTGGATCTGGTCCTCGAAGGCGACAGCTGCGTGACTATCATGTCCAAGGATAAGCCCACCATCGATGTGAAGATGATGAACATGGAGGCTGCCAATCTGGCGGAGGTTCGAAGCTATTGTTATTTGGCAACCGTCAGTGATCTCTCCACTAAAGCTGCCTGCCCGACCATGGGAGAAGCGCACAACGACAAACGTGCTGACCCTGCATTTGTGTGCAGACAAGGAGTGGTGGACAGGGGCTGGGGCAACGGATGCGGCCTTTTTGGGAAAGGCAGCATTGACACATGCGCCAAATTTGCTTGTTCCACTAAAGCAATAGGGAGAACCATTTTGAAAGAGAACATCAAGTACGAAGTGGCTATCTTTGTCCATGGACCAACTACAGTGGAGTCTCACGGCAACTACTCCACCCAGGTTGGAGCCACACAGGCAGGAAGACTCAGCATCACACCAGCAGCTCCATCATACACACTAAAACTTGGAGAATATGGCGAGGTCACAGTTGACTGTGAACCTCGGTCAGGAATTGACACCAATGCCTACTACGTGATGACTGTGGGAACTAAGACGTTCTTGGTTCACCGTGAGTGGTTCATGGACCTGAACCTGCCTTGGAGCAGCGCGGGAAGCACAGTGTGGAGGAACCGAGAGACTTTGATGGAGTTCGAGGAGCCGCACGCCACTAAGCAGTCTGTGATAGCCTTGGGCTCTCAGGAGGGAGCGCTGCACCAAGCATTGGCTGGAGCCATCCCAGTTGAATTCTCCAGTAACACAGTCAAGTTGACGTCGGGTCATTTAAAATGTAGAGTGAAGATGGAAAAATTGCAGTTGAAAGGAACAACTTATGGAGTATGCTCAAAAGCGTTCAAGTTCCTTGGGACACCCGCTGACACTGGACACGGCACGGTGGTCTTGGAATTGCAGTACACTGGCACGGATGGACCTTGCAAGGTTCCGATCTCGTCTGTGGCCTCGTTAAACGATTTGACGCCTGTGGGCAGATTGGTGACTGTCAATCCTTTTGTGTCAGTGGCCACTGCCAATGCTAAGGTCCTCATTGAGTTGGAACCTCCATTTGGAGACTCATACATAGTGGTGGGCAGAGGAGAACAACAGATCAACCATCATTGGCACAAGTCTGGCAGCAGCATTGGCAAAGCTTTCACAACCACTCTCAAAGGAGCACAGAGACTAGCCGCTCTAGGAGACACAGCCTGGGACTTTGGCTCAGTTGGAGGGGTCTTTACATCAGTTGGGAAAGCTGTCCACCAGGTGTTTGGAGGAGCTTTCAGATCACTGTTCGGAGGAATGTCTTGGATAACTCAGGGCTTGCTGGGGGCTCTTCTGTTGTGGATGGGAATCAATGCGCGTGATAGGTCAATAGCACTCACGTTCCTCGCAGTTGGAGGAGTTCTTCTGTTCCTGTCTGTGAACGTGCACGCTGACACGGGGTGTGCCATTGATATCAGCCGCCAAGAGCTGAGATGTGGAAGTGGGGTGTTCATACACAACGATGTGGAGGCTTGGATGGATCGGTACAAGTATTACCCAGAAACGCCACAAGGACTAGCCAAGATCATACAGAAGGCACATAAAGAAGGCGTGTGCGGTCTGCGATCTGTTTCCAGGCTGGAACACCAAATGTGGGAGGCTGTGAAGGATGAACTCAATACACTTTTGAAGGAAAACGGTGTGGATCTCAGTGTCGTGGTTGAGAAGCAAGAGGGAATGTACAAGTCCGCTCCCAAGCGCCTGACGGCCACGACAGAAAAATTGGAAATTGGCTGGAAGGCCTGGGGTAAGAGCATTTTATTTGCACCGGAACTAGCCAACAACACCTTTGTTGTGGATGGTCCTGAGACCAAAGAATGTCCGACTCAGAATCGTGCTTGGAACAGTTTGGAAGTAGAGGATTTTGGATTCGGACTCACCAGTACTCGGATGTTCCTCAAGGTCAGAGAGAGCAACACAACAGAATGTGACTCTAAGATCATTGGAACTGCTGTGAAGAACAACTTGGCCATCCACAGTGACCTGTCATATTGGATTGAGAGCAGGCTAAATGACACCTGGAAGCTGGAGAGGGCAGTACTTGGTGAGGTTAAATCCTGTACGTGGCCTGAGACCCATACATTGTGGGGCGATGGAATCCTCGAGAGCGACTTGATAATTCCAGTCACACTGGCTGGACCTCGAAGCAACCACAATCGGAGACCGGGGTACAAGACACAAAACCAGGGTCCCTGGGATGAAGGCCGAGTAGAAATTGACTTTGATTATTGCCCAGGAACCACAGTCACGCTGAGCGAAAGCTGTGGACACCGAGGCCCGGCCACTCGCACTACGACAGAGAGTGGGAAGTTGATCACAGACTGGTGCTGCAGAAGCTGCACGTTACCACCACTTCGCTACCAAACTGACAGCGGCTGCTGGTATGGAATGGAGATCAGACCACAGAGACATGATGAGAAGACCCTCGTGCAGTCACAAGTGAATGCTTATAATGCTGATATGATTGACCCTTTTCAGTTGGGCCTTCTGGTCGTGTTCTTGGCCACCCAGGAGGTCCTTCGC

>wt-NY99_**Fragment-II**

TATAATGCTGATATGATTGACCCTTTTCAGTTGGGCCTTCTGGTCGTGTTCTTGGCCACCCAGGAGGTCCTTCGCAAGAGGTGGACAGCCAAGATCAGCATGCCAGCTATACTGATTGCTCTGCTAGTCCTGGTGTTTGGGGGCATTACTTACACTGATGTGTTACGCTATGTCATCTTGGTGGGGGCAGCTTTCGCAGAATCTAATTCGGGAGGAGACGTGGTACACTTGGCGCTCATGGCGACCTTCAAGATACAACCAGTGTTTATGGTGGCATCGTTTCTCAAAGCGAGATGGACCAACCAGGAGAACATTTTGTTGATGTTGGCGGCTGTTTTCTTTCAAATGGCTTATCACGATGCCCGCCAAATTCTGCTCTGGGAGATCCCTGATGTGTTGAATTCACTGGCGGTAGCTTGGATGATACTGAGAGCCATAACATTCACAACGACATCAAACGTGGTTGTTCCGCTGCTAGCCCTGCTAACACCCGGGCTGAGATGCTTGAATCTGGATGTGTACAGGATACTGCTGTTGATGGTCGGAATAGGCAGCTTGATCAGGGAGAAGAGGAGTGCAGCCGCAAAAAAGAAAGGAGCAAGTCTGCTATGCTTGGCTCTAGCCTCAACAGGACTTTTCAACCCCATGATCCTTGCTGCTGGACTGATTGCATGTGATCCCAACCGTAAACGCGGATGGCCCGCAACTGAAGTGATGACAGCTGTCGGCCTAATGTTTGCCATCGTCGGAGGGCTGGCAGAGCTTGACATTGACTCCATGGCCATTCCAATGACTATCGCGGGGCTCATGTTTGCTGCTTTCGTGATTTCTGGGAAATCAACAGATATGTGGATTGAGAGAACGGCGGACATTTCCTGGGAAAGTGATGCAGAAATTACAGGCTCGAGCGAAAGAGTTGATGTGCGGCTTGATGATGATGGAAACTTCCAGCTCATGAATGATCCAGGAGCACCTTGGAAGATATGGATGCTCAGAATGGTCTGTCTCGCGATTAGTGCGTACACCCCCTGGGCAATCTTGCCCTCAGTAGTTGGATTTTGGATAACTCTCCAATACACAAAGAGAGGAGGCGTGTTGTGGGACACTCCCTCACCAAAGGAGTACAAAAAGGGGGACACGACCACCGGCGTCTACAGGATCATGACTCGTGGGCTGCTCGGCAGTTATCAAGCAGGAGCGGGCGTGATGGTTGAAGGTGTTTTCCACACCCTTTGGCATACAACAAAAGGAGCCGCTTTGATGAGCGGAGAGGGCCGCCTGGACCCATACTGGGGCAGTGTCAAGGAGGATCGACTTTGTTACGGAGGACCCTGGAAATTGCAGCACAAGTGGAACGGGCAGGATGAGGTGCAGATGATTGTGGTGGAACCTGGCAAGAACGTTAAGAACGTCCAGACGAAACCAGGGGTGTTCAAAACACCTGAAGGAGAAATCGGGGCCGTGACTTTGGACTTCCCCACTGGAACATCAGGCTCACCAATAGTGGACAAAAACGGTGATGTGATTGGGCTTTATGGCAATGGAGTCATAATGCCCAACGGCTCATACATAAGCGCGATAGTGCAGGGTGAAAGGATGGATGAGCCAATCCCAGCCGGATTCGAACCTGAGATGCTGAGGAAAAAACAGATCACTGTACTGGATCTCCATCCCGGCGCCGGTAAAACAAGGAGGATTCTGCCACAGATCATCAAAGAGGCCATAAACAGAAGACTGAGAACAGCCGTGCTAGCGCCAACCAGGGTTGTGGCTGCTGAGATGGCTGAAGCACTGAGAGGACTGCCCATCCGGTACCAGACATCCGCAGTGCCCAGAGAACATAATGGAAATGAGATTGTTGATGTCATGTGTCATGCTACCCTCACCCACAGGCTGATGTCTCCTCACAGGGTGCCGAACTACAACCTGTTCGTGATGGATGAGGCTCATTTCACCGACCCAGCTAGCATTGCAGCAAGAGGTTACATTTCCACAAAGGTCGAGCTAGGGGAGGCGGCGGCAATATTCATGACAGCCACCCCACCAGGCACTTCAGATCCATTCCCAGAGTCCAATTCACCAATTTCCGACTTACAGACTGAGATCCCGGATCGAGCTTGGAACTCTGGATACGAATGGATCACAGAATACACCGGGAAGACGGTTTGGTTTGTGCCTAGTGTCAAGATGGGGAATGAGATTGCCCTTTGCCTACAACGTGCTGGAAAGAAAGTAGTCCAATTGAACAGAAAGTCGTACGAGACGGAGTACCCAAAATGTAAGAACGATGATTGGGACTTTGTTATCACAACAGACATATCTGAAATGGGGGCTAACTTCAAGGCGAGCAGGGTGATTGACAGCCGGAAGAGTGTGAAACCAACCATCATAACAGAAGGAGAAGGGAGAGTGATCCTGGGAGAACCATCTGCAGTGACAGCAGCTAGTGCCGCCCAGAGACGTGGACGTATCGGTAGAAATCCGTCGCAAGTTGGTGATGAGTACTGTTATGGGGGGCACACGAATGAAGACGACTCGAACTTCGCCCATTGGACTGAGGCACGAATCATGCTGGACAACATCAACATGCCAAACGGACTGATCGCTCAATTCTACCAACCAGAGCGTGAGAAGGTATATACCATGGATGGGGAATACCGGCTCAGAGGAGAAGAGAGAAAAAACTTTCTGGAACTGTTGAGGACTGCAGATCTGCCAGTTTGGCTGGCTTACAAGGTTGCAGCGGCTGGAGTGTCATACCACGACCGGAGGTGGTGCTTTGATGGTCCTAGGACAAACACAATTTTAGAAGACAACAACGAAGTGGAAGTCATCACGAAGCTTGGTGAAAGGAAGATTCTGAGGCCGCGCTGGATTGATGCCAGGGTGTACTCGGATCACCAGGCACTAAAGGCGTTCAAGGACTTCGCCTCGGGAAAACGTTCTCAGATAGGGCTCATTGAGGTTCTGGGAAAGATGCCTGAGCACTTCATGGGGAAGACATGGGAAGCACTTGACACCATGTACGTTGTGGCCACTGCAGAGAAAGGAGGAAGAGCTCACAGAATGGCCCTGGAGGAACTGCCAGATGCTCTTCAGACAATTGCCTTGATTGCCTTATTGAGTGTGATGACCATGGGAGTATTCTTCCTCCTCATGCAGCGGAAGGGCATTGGAAAGATAGGTTTGGGAGGCGCTGTCTTGGGAGTCGCGACCTTTTTCTGTTGGATGGCTGAAGTTCCAGGAACGAAGATCGCCGGAATGTTGCTGCTCTCCCTTCTCTTGATGATTGTGCTAATTCCTGAGCCAGAGAAGCAACGTTCGCAGACAGACAACCAGCTAGCCGTGTTCCTGATTTGTGTCATGACCCTTGTGAGCGCAGTGGCAGCCAACGAGATGGGTTGGCTAGATAAGACCAAGAGTGACATAAGCAGTTTGTTTGGGCAAAGAATTGAGGTCAAGGAGAATTTCAGCATGGGAGAGTTTCTTCTGGACTTGAGGCCGGCAACAGCCTGGTCACTGTACGCTGTGACAACAGCGGTCCTCACTCCACTGCTAAAGCATTTGATCACGTCAGATTACATCAACACCTCATTGACCTCAATAAACGTTCAGGCAAGTGCACTATTCACACTCGCGCGAGGCTTCCCCTTCGTCGATGTTGGAGTGTCGGCTCTCCTGCTAGCAGCCGGATGCTGGGGACAAGTCACCCTCACCGTTACGGTAACAGCGGCAACACTCCTTTTTTGCCACTATGCCTACATGGTTCCCGGTTGGCAAGCTGAGGCAATGCGCTCAGCCCAGCGGCGGACAGCGGCCGGAATCATGAAGAACGCTGTAGTGGATGGCATCGTGGCCACGGACGTCCCAGAATTAGAGCGCACCACACCCATCATGCAGAAGAAAGTTGGACAGATCATGCTGATCTTGGTGTCTCTAGCTGCAGTAGTAGTGAACCCGTCTGTGAAGACAGTACGAGAAGCCGGAATTTTGATCACGGCCGCAGCGGTGACGCTTTGGGAGAATGGAGCAAGCTCTGTTTGGAACGCAACAACTGCCATCGGACTCTGCCACATCATGCGTGGGGGTTGGTTGTCATGTCTATCCATAACATGGACACTCATAAAGAACATGGAAAAACCAGGACTAAAAAGAGGTGGGGCAAAAGGACGCACCTTGGGAGAGGTTTGGAAAGAAAGACTCAACCAGATGACAAAAGAAGAGTTCACT

>wt-NY99_**Fragment-III**

GGTGGGGCAAAAGGACGCACCTTGGGAGAGGTTTGGAAAGAAAGACTCAACCAGATGACAAAAGAAGAGTTCACTAGGTACCGCAAAGAGGCCATCATCGAAGTCGATCGCTCAGCGGCAAAACACGCCAGGAAAGAAGGCAATGTCACTGGAGGGCATCCAGTCTCTAGGGGCACAGCAAAACTGAGATGGCTGGTCGAACGGAGGTTTCTCGAACCGGTCGGAAAAGTGATTGACCTTGGATGTGGAAGAGGCGGTTGGTGTTACTATATGGCAACCCAAAAAAGAGTCCAAGAAGTCAGAGGGTACACAAAGGGCGGTCCCGGACATGAAGAGCCCCAACTAGTGCAAAGTTATGGATGGAACATTGTCACCATGAAGAGTGGAGTGGATGTGTTCTACAGACCTTCTGAGTGTTGTGACACCCTCCTTTGTGACATCGGAGAGTCCTCGTCAAGTGCTGAGGTTGAAGAGCATAGGACGATTCGGGTCCTTGAAATGGTTGAGGACTGGCTGCACCGAGGGCCAAGGGAATTTTGCGTGAAGGTGCTCTGTCCCTACATGCCGAAAGTCATAGAGAAGATGGAGCTGCTCCAACGCCGGTATGGGGGGGGACTGGTCAGAAACCCACTCTCACGGAATTCCACGCACGAGATGTATTGGGTGAGTCGAGCTTCAGGCAATGTGGTACATTCAGTGAATATGACCAGCCAGGTGCTCCTAGGAAGAATGGAAAAAAGGACCTGGAAGGGACCCCAATACGAGGAAGATGTAAACTTGGGAAGTGGAACCAGGGCGGTGGGAAAACCCCTGCTCAACTCAGACACCAGTAAAATCAAGAACAGGATTGAACGACTCAGGCGTGAGTACAGTTCGACGTGGCACCACGATGAGAACCACCCATATAGAACCTGGAACTATCACGGCAGTTATGATGTGAAGCCCACAGGCTCCGCCAGTTCGCTGGTCAATGGAGTGGTCAGGCTCCTCTCAAAACCATGGGACACCATCACGAATGTTACCACCATGGCCATGACTGACACTACTCCCTTCGGGCAGCAGCGAGTGTTCAAAGAGAAGGTGGACACGAAAGCTCCTGAACCGCCAGAAGGAGTGAAGTACGTGCTCAACGAGACCACCAACTGGTTGTGGGCGTTTTTGGCCAGAGAAAAACGTCCCAGAATGTGCTCTCGAGAGGAATTCATAAGAAAGGTCAACAGCAATGCAGCTTTGGGTGCCATGTTTGAAGAGCAGAATCAATGGAGGAGCGCCAGAGAAGCAGTTGAAGATCCAAAATTTTGGGAGATGGTGGATGAGGAGCGCGAGGCACATCTGCGGGGGGAATGTCACACTTGCATTTACAACATGATGGGAAAGAGAGAGAAAAAACCCGGAGAGTTCGGAAAGGCCAAGGGAAGCAGAGCCATTTGGTTCATGTGGCTCGGAGCTCGCTTTCTGGAGTTCGAGGCTCTGGGTTTTCTCAATGAAGACCACTGGCTTGGAAGAAAGAACTCAGGAGGAGGTGTCGAGGGCTTGGGCCTCCAAAAACTGGGTTACATCCTGCGTGAAGTTGGCACCCGGCCTGGGGGCAAGATCTATGCTGATGACACAGCTGGCTGGGACACCCGCATCACGAGAGCTGACTTGGAAAATGAAGCTAAGGTGCTTGAGCTGCTTGATGGGGAACATCGGCGTCTTGCCAGGGCCATCATTGAGCTCACCTATCGTCACAAAGTTGTGAAAGTGATGCGCCCGGCTGCTGATGGAAGAACCGTCATGGATGTTATCTCCAGAGAAGATCAGAGGGGGAGTGGACAAGTTGTCACCTACGCCCTAAACACTTTCACCAACCTGGCCGTCCAGCTGGTGAGGATGATGGAAGGGGAAGGAGTGATTGGCCCAGATGATGTGGAGAAACTCACAAAAGGGAAAGGACCCAAAGTCAGGACCTGGCTGTTTGAGAATGGGGAAGAAAGACTCAGCCGCATGGCTGTCAGTGGAGATGACTGTGTGGTAAAGCCCCTGGACGATCGCTTTGCCACCTCGCTCCACTTCCTCAATGCTATGTCAAAGGTTCGCAAAGACATCCAAGAGTGGAAACCGTCAACTGGATGGTATGATTGGCAGCAGGTTCCATTTTGCTCAAACCATTTCACTGAATTGATCATGAAAGATGGAAGAACACTGGTGGTTCCATGCCGAGGACAGGATGAATTGGTAGGCAGAGCTCGCATATCTCCAGGGGCCGGATGGAACGTCCGCGACACTGCTTGTCTGGCTAAGTCTTATGCCCAGATGTGGCTGCTTCTGTACTTCCACAGAAGAGACCTGCGGCTCATGGCCAACGCCATTTGCTCCGCTGTCCCTGTGAATTGGGTCCCTACCGGAAGAACCACGTGGTCCATCCATGCAGGAGGAGAGTGGATGACAACAGAGGACATGTTGGAGGTCTGGAACCGTGTTTGGATAGAGGAGAATGAATGGATGGAAGACAAAACCCCAGTGGAGAAATGGAGTGACGTCCCATATTCAGGAAAACGAGAGGACATCTGGTGTGGCAGCCTGATTGGCACAAGAGCCCGAGCCACGTGGGCAGAAAACATCCAGGTGGCTATCAACCAAGTCAGAGCAATCATCGGAGATGAGAAGTATGTGGACTACATGAGTTCACTAAAGAGATATGAAGACACAACTTTGGTTGAGGACACAGTACTGTAGATATTTAATCAATTGTAAATAGACAATATAAGTATGCATAAAAGTGTAGTTTTATAGTAGTATTTAGTGGTGTTAGTGTAAATAGTTAAGAAAATTTTGAGGAGAAAGTCAGGCCGGGAAGTTCCCGCCACCGGAAGTTGAGTAGACGGTGCTGCCTGCGACTCAACCCCAGGAGGACTGGGTGAACAAAGCCGCGAAGTGATCCATGTAAGCCCTCAGAACCGTCTCGGAAGGAGGACCCCACATGTTGTAACTTCAAAGCCCAATGTCAGACCACGCTACGGCGTGCTACTCTGCGGAGAGTGCAGTCTGCGATAGTGCCCCAGGAGGACTGGGTTAACAAAGGCAAACCAACGCCCCACGCGGCCCTAGCCCCGGTAATGGTGTTAACCAGGGCGAAAGGACTAGAGGTTAGAGGAGACCCCGCGGTTTAAAGTGCACGGCCCAGCCTGGCTGAAGCTGTAGGTCAGGGGAAGGACTAGAGGTTAGTGGAGACCCCGTGCCACAAAACACCACAACAAAACAGCATATTGACACCTGGGATAGACTAGGAGATCTTCTGCTCTGCACAACCAGCCACACGGCACAGTGCGCCGACAATGGTGGCTGGTGGTGCGAGAACACAGGATCT**GGCCGGCATGGTCCCAGCCTCCTCGCTGGCGCCGGCTGGGCAACATTCCGAGGGGACCGTCCCCTCGGTAATGGCGAATGGGACTCGCGACAGACATGATAAGATACATTGATGAGTTTGGACAAACCACAACTAGAATGCAGTGAAAAAAATGCTTTATTTGTGAAATTAAGCGCTGGCATTGACCCTGAG**

To overcome the toxicity of NY99-Pr-E-NS1_**Fragment-I (see above)**, we employed an alternative ISA strategy. The initial NY99-Pr-E-NS1_**Fragment-I** was synthesized as **three** shorter, overlapping DNA fragments (**A**, **B,** and **c**). **Fragment-I-A** and **Fragment-I-B** DNA fragments were synthesized *de novo* without insertion into bacterial plasmids or preamplification in bacteria, using the service provided by Twist Biosciences. For the **Fragment-I-c** we used *synthetic clonal genes* with the pTwist Amp MC medium copy plasmid. The rescue of **Fragment-I-c** fragment failed due to its toxicity to bacteria. The alternative ISA strategy is described below.

***In brow are 22 nt adapters applied during synthesis of DNA without using bacterial plasmids and bacteria.***

>NY99-Pr-E-NS1_**Fragment-I-A**

***caatccgccctcactacaaccg*CACCCAACTGATCTTCAGCATCTTCAATATTGGCCATTAGCCATATTATTCATTGGTTATATAGCATAAATCAATATTGGCTATTGGCCATTGCATACGTTGTATCTATATCATAATATGTACATTTATATTGGCTCATGTCCAATATGACCGCCATGTTGGCATTGATTATTGACTAGTTATTAATAGTAATCAATTACGGGGTCATTAGTTCATAGCCCATATATGGAGTTCCGCGTTACATAACTTACGGTAAATGGCCCGCCTGGCTGACCGCCCAACGACCCCCGCCCATTGACGTCAATAATGACGTATGTTCCCATAGTAACGCCAATAGGGACTTTCCATTGACGTCAATGGGTGGAGTATTTACGGTAAACTGCCCACTTGGCAGTACATCAAGTGTATCATATGCCAAGTCCGCCCCCTATTGACGTCAATGACGGTAAATGGCCCGCCTGGCATTATGCCCAGTACATGACCTTACGGGACTTTCCTACTTGGCAGTACATCTACGTATTAGTCATCGCTATTACCATGGTGATGCGGTTTTGGCAGTACACCAATGGGCGTGGATAGCGGTTTGACTCACGGGGATTTCCAAGTCTCCACCCCATTGACGTCAATGGGAGTTTGTTTTGGCACCAAAATCAACGGGACTTTCCAAAATGTCGTAATAACCCCGCCCCGTTGACGCAAATGGGCGGTAGGCGTGTACGGTGGGAGGTCTATATAAGCAGAGCTCGTTTAGTGAACCG**AGTAGTTCGCCTGTGTGAGCTGACAAACTTAGTAGTGTTTGTGAGGATTAACAACAATTAACACAGTGCGAGCTGTTTCTTAGCACGAAGATCTCGATGTCTAAGAAACCAGGAGGGCCCGGCAAGAGCCGGGCTGTCAATATGCTAAAACGCGGAATGCCCCGCGTGTTGTCCTTGATTGGACTGAAGAGGGCTATGTTGAGCCTGATCGACGGCAAGGGGCCAATACGATTTGTGTTGGCTCTCTTGGCGTTCTTCAGGTTCACAGCAATTGCTCCGACCCGAGCAGTGCTGGATCGATGGAGAGGTGTGAACAAACAAACAGCGATGAAACACCTTCTGAGTTTTAAGAAGGAACTAGGGACCTTGACCAGTGCTATCAATCGGCGGAGCTCAAAACAAAAGAAAAGAGGAGGAAAGACCGGAATTGCAGTCATGAT***ctactctggcgtcgatgaggga***

>NY99-Pr-E-NS1_**Fragment-I-B**

***caatccgccctcactacaaccg***TGACCAGTGCTATCAATCGGCGGAGCTCAAAACAAAAGAAAAGAGGAGGAAAGACCGGAATTGCAGTCATGATTGGCCTGATCGCCAGCGTAGGAGCAGTTACCCTCTCTAACTTCCAAGGGAAGGTGATGATGACGGTAAATGCTACTGACGTCACAGATGTCATCACGATTCCAACAGCTGCTGGAAAGAACCTATGCATTGTCAGAGCAATGGATGTGGGATACATGTGCGATGATACTATCACTTATGAATGCCCAGTACTGTCGGCTGGTAATGATCCAGAAGACATCGACTGTTGGTGCACAAAGTCAGCAGTCTACGTCAGGTATGGAAGATGCACCAAGACACGCCACTCAAGACGCAGTCGGAGGTCACTGACAGTGCAGACACACGGAGAAAGCACTCTAGCGAACAAGAAGGGGGCTTGGATGGACAGCACCAAGGCCACAAGGTATTTGGTAAAAACAGAATCATGGATCTTGAGGAACCCTGGATATGCCCTGGTGGCAGCCGTCATTGGTTGGATGCTTGGGAGCAACACCATGCAGAGAGTTGTGTTTGTCGTGCTATTGCTTTTGGTGGCCCCAGCTTACAGCTTCAACTGTCTGGGCATGAGCAATAGAGACTTTTTGGAAGGAGTCTCCGGAGCAACCTGGGTGGATCTGGTCCTCGAAGGCGACAGCTGCGTGACTATCATGTCCAAGGATAAGCCCACCATCGATGTGAAGATGATGAACATGGAGGCTGCCAATCTGGCGGAGGTTCGAAGCTATTGTTATTTGGCAACCGTCAGTGATCTCTCCACTAAAGCTGCCTGCCCGACCATGGGAGAAGCGCACAACGACAAACGTGCTGACCCTGCATTTGTGTGCAGACAAGGAGTGGTGGACAGGGGCTGGGGCAACGGATGCGGCCTTTTTGGGAAAGGCAGCATTGACACATGCGCCAAATTTGCTTGTTCCACTAAAGCAATAGGGAGAACCATTTTGAAAGAGAACATCAAGTACGAAGTGGCTATCTTTGTCCATGGACCAACTACAGTGGAGTCTCACGGCAACTACTCCACCCAGGTTGGAGCCACACAGGCAGGAAGACTCAGCATCACACCAGCAGCTCCATCATA***ctactctggcgtcgatgaggga***

> NY99-Pr-E-NS1_**Fragment-I-c**

The rescue of this fragment failed due to its toxicity to bacteria. The alternative ISA strategy is described below.

CGGCAACTACTCCACCCAGGTTGGAGCCACACAGGCAGGAAGACTCAGCATCACACCAGCAGCTCCATCATACACACTAAAACTTGGAGAATATGGCGAGGTCACAGTTGACTGTGAACCTCGGTCAGGAATTGACACCAATGCCTACTACGTGATGACTGTGGGAACTAAGACGTTCTTGGTTCACCGTGAGTGGTTCATGGACCTGAACCTGCCTTGGAGCAGCGCGGGAAGCACAGTGTGGAGGAACCGAGAGACTTTGATGGAGTTCGAGGAGCCGCACGCCACTAAGCAGTCTGTGATAGCCTTGGGCTCTCAGGAGGGAGCGCTGCACCAAGCATTGGCTGGAGCCATCCCAGTTGAATTCTCCAGTAACACAGTCAAGTTGACGTCGGGTCATTTAAAATGTAGAGTGAAGATGGAAAAATTGCAGTTGAAAGGAACAACTTATGGAGTATGCTCAAAAGCGTTCAAGTTCCTTGGGACACCCGCTGACACTGGACACGGCACGGTGGTCTTGGAATTGCAGTACACTGGCACGGATGGACCTTGCAAGGTTCCGATCTCGTCTGTGGCCTCGTTAAACGATTTGACGCCTGTGGGCAGATTGGTGACTGTCAATCCTTTTGTGTCAGTGGCCACTGCCAATGCTAAGGTCCTCATTGAGTTGGAACCTCCATTTGGAGACTCATACATAGTGGTGGGCAGAGGAGAACAACAGATCAACCATCATTGGCACAAGTCTGGCAGCAGCATTGGCAAAGCTTTCACAACCACTCTCAAAGGAGCACAGAGACTAGCCGCTCTAGGAGACACAGCCTGGGACTTTGGCTCAGTTGGAGGGGTCTTTACATCAGTTGGGAAAGCTGTCCACCAGGTGTTTGGAGGAGCTTTCAGATCACTGTTCGGAGGAATGTCTTGGATAACTCAGGGCTTGCTGGGGGCTCTTCTGTTGTGGATGGGAATCAATGCGCGTGATAGGTCAATAGCACTCACGTTCCTCGCAGTTGGAGGAGTTCTTCTGTTCCTGTCTGTGAACGTGCACGCTGACACGGGGTGTGCCATTGATATCAGCCGCCAAGAGCTGAGATGTGGAAGTGGGGTGTTCATACACAACGATGTGGAGGCTTGGATGGATCGGTACAAGTATTACCCAGAAACGCCACAAGGACTAGCCAAGATCATACAGAAGGCACATAAAGAAGGCGTGTGCGGTCTGCGATCTGTTTCCAGGCTGGAACACCAAATGTGGGAGGCTGTGAAGGATGAACTCAATACACTTTTGAAGGAAAACGGTGTGGATCTCAGTGTCGTGGTTGAGAAGCAAGAGGGAATGTACAAGTCCGCTCCCAAGCGCCTGACGGCCACGACAGAAAAATTGGAAATTGGCTGGAAGGCCTGGGGTAAGAGCATTTTATTTGCACCGGAACTAGCCAACAACACCTTTGTTGTGGATGGTCCTGAGACCAAAGAATGTCCGACTCAGAATCGTGCTTGGAACAGTTTGGAAGTAGAGGATTTTGGATTCGGACTCACCAGTACTCGGATGTTCCTCAAGGTCAGAGAGAGCAACACAACAGAATGTGACTCTAAGATCATTGGAACTGCTGTGAAGAACAACTTGGCCATCCACAGTGACCTGTCATATTGGATTGAGAGCAGGCTAAATGACACCTGGAAGCTGGAGAGGGCAGTACTTGGTGAGGTTAAATCCTGTACGTGGCCTGAGACCCATACATTGTGGGGCGATGGAATCCTCGAGAGCGACTTGATAATTCCAGTCACACTGGCTGGACCTCGAAGCAACCACAATCGGAGACCGGGGTACAAGACACAAAACCAGGGTCCCTGGGATGAAGGCCGAGTAGAAATTGACTTTGATTATTGCCCAGGAACCACAGTCACGCTGAGCGAAAGCTGTGGACACCGAGGCCCGGCCACTCGCACTACGACAGAGAGTGGGAAGTTGATCACAGACTGGTGCTGCAGAAGCTGCACGTTACCACCACTTCGCTACCAAACTGACAGCGGCTGCTGGTATGGAATGGAGATCAGACCACAGAGACATGATGAGAAGACCCTCGTGCAGTCACAAGTGAATGCTTATAATGCTGATATGATTGACCCTTTTCAGTTGGGCCTTCTGGTCGTGTTCTTGGCCACCCAGGAGGTCCTTCGC

To overcome the toxicity of NY99-Pr-E-NS1_**Fragment-I-c (see above)**, we employed an alternative ISA strategy. The initial toxic NY99-Pr-E-NS1_**Fragment-Ic** was synthesized as **two** shorter, overlapping DNA fragments (**C** and **D**). These DNA fragments were synthesized *de novo* without insertion into bacterial plasmids or preamplification in bacteria, using the service provided by Twist Biosciences.

> NY99-Pr-E-NS1_**Fragment-I-C**

***caatccgccctcactacaaccg***CGGCAACTACTCCACCCAGGTTGGAGCCACACAGGCAGGAAGACTCAGCATCACACCAGCAGCTCCATCATACACACTAAAACTTGGAGAATATGGCGAGGTCACAGTTGACTGTGAACCTCGGTCAGGAATTGACACCAATGCCTACTACGTGATGACTGTGGGAACTAAGACGTTCTTGGTTCACCGTGAGTGGTTCATGGACCTGAACCTGCCTTGGAGCAGCGCGGGAAGCACAGTGTGGAGGAACCGAGAGACTTTGATGGAGTTCGAGGAGCCGCACGCCACTAAGCAGTCTGTGATAGCCTTGGGCTCTCAGGAGGGAGCGCTGCACCAAGCATTGGCTGGAGCCATCCCAGTTGAATTCTCCAGTAACACAGTCAAGTTGACGTCGGGTCATTTAAAATGTAGAGTGAAGATGGAAAAATTGCAGTTGAAAGGAACAACTTATGGAGTATGCTCAAAAGCGTTCAAGTTCCTTGGGACACCCGCTGACACTGGACACGGCACGGTGGTCTTGGAATTGCAGTACACTGGCACGGATGGACCTTGCAAGGTTCCGATCTCGTCTGTGGCCTCGTTAAACGATTTGACGCCTGTGGGCAGATTGGTGACTGTCAATCCTTTTGTGTCAGTGGCCACTGCCAATGCTAAGGTCCTCATTGAGTTGGAACCTCCATTTGGAGACTCATACATAGTGGTGGGCAGAGGAGAACAACAGATCAACCATCATTGGCACAAGTCTGGCAGCAGCATTGGCAAAGCTTTCACAACCACTCTCAAAGGAGCACAGAGACTAGCCGCTCTAGGAGACACAGCCTGGGACTTTGGCTCAGTTGGAGGGGTCTTTACATCAGTTGGGAAAGCTGTCCACCAGGTGTTTGGAGGAGCTTTCAGATCACTGTTCGGAGGAATGTCTTGGATAACTCAGGGCTTGCTGGGGGCTCTTCTGTTGTGGATGGGAATCAATGCGCGTGATAGGTCAATAGCACTCACGTTCCTCGCAGTTGGAGGAGTTCTTCTGTTCCTGTCTGTGAACGTGCACGCTGACACGGGGTGTGCCATTGATATCAGCCGCCAAGAGCTGAGATGTGGAAGTGGGGTGTTCATACACAACGATGTGGAGGCTTGGATGGATCGGTACAAGTATTACCCAGAAACG***ctactctggcgtcgatgaggga***

> NY99-Pr-E-NS1_**Fragment-I-D**

***caatccgccctcactacaaccg***TGTGGAAGTGGGGTGTTCATACACAACGATGTGGAGGCTTGGATGGATCGGTACAAGTATTACCCAGAAACGCCACAAGGACTAGCCAAGATCATACAGAAGGCACATAAAGAAGGCGTGTGCGGTCTGCGATCTGTTTCCAGGCTGGAACACCAAATGTGGGAGGCTGTGAAGGATGAACTCAATACACTTTTGAAGGAAAACGGTGTGGATCTCAGTGTCGTGGTTGAGAAGCAAGAGGGAATGTACAAGTCCGCTCCCAAGCGCCTGACGGCCACGACAGAAAAATTGGAAATTGGCTGGAAGGCCTGGGGTAAGAGCATTTTATTTGCACCGGAACTAGCCAACAACACCTTTGTTGTGGATGGTCCTGAGACCAAAGAATGTCCGACTCAGAATCGTGCTTGGAACAGTTTGGAAGTAGAGGATTTTGGATTCGGACTCACCAGTACTCGGATGTTCCTCAAGGTCAGAGAGAGCAACACAACAGAATGTGACTCTAAGATCATTGGAACTGCTGTGAAGAACAACTTGGCCATCCACAGTGACCTGTCATATTGGATTGAGAGCAGGCTAAATGACACCTGGAAGCTGGAGAGGGCAGTACTTGGTGAGGTTAAATCCTGTACGTGGCCTGAGACCCATACATTGTGGGGCGATGGAATCCTCGAGAGCGACTTGATAATTCCAGTCACACTGGCTGGACCTCGAAGCAACCACAATCGGAGACCGGGGTACAAGACACAAAACCAGGGTCCCTGGGATGAAGGCCGAGTAGAAATTGACTTTGATTATTGCCCAGGAACCACAGTCACGCTGAGCGAAAGCTGTGGACACCGAGGCCCGGCCACTCGCACTACGACAGAGAGTGGGAAGTTGATCACAGACTGGTGCTGCAGAAGCTGCACGTTACCACCACTTCGCTACCAAACTGACAGCGGCTGCTGGTATGGAATGGAGATCAGACCACAGAGACATGATGAGAAGACCCTCGTGCAGTCACAAGTGAATGCTTATAATGCTGATATGATTGACCCTTTTCAGTTGGGCCTTCTGGTCGTGTTCTTGGCCACCCAGGAGGTCCTTCGC***ctactctggcgtcgatgaggga***

For NY99-Pr-E-NS1_**Fragment-I-A**, NY99-Pr-E-NS1_**Fragment-I-B**, wt-NY99_**Fragment-II** and wt-NY99_**Fragment-III** the same sequences as above were used.
