## supplemental tables and files for "Infectious subgenomic amplicon strategies for Japanese encephalitis and West Nile viruses": Table S1 ISA fragments and primers.docx

**Synthetic clonal genes**—DNA fragments are synthesized, inserted into the bacterial plasmids, amplified in bacteria, extracted, and plasmids with inserted fragments are delivered as lyophilized DNA with known concentration.

***Synthesis of DNA***—DNA fragments, flanked with 22 nt adapters, are rapidly (2-4 days) synthesized without using bacterial plasmids and bacteria.

*****This fragment was ordered from three companies—GenScript, GeneArt, and Twist Biosciences.

For sequences of each fragment, see **File S1**.

| **Virus**  **(*ISA troubleshooting*)** | **Fragment ID** | **Plasmids** | **Fragment origin** | **Primers** |
| --- | --- | --- | --- | --- |
| **DENV**  **wild-type** |  |  |  |  |
|  | DENV2_16681_**Fragment-I** | pTwist Amp MC | synthetic clonal genes | DENV2_16681-I-Forward: CACCCAACTGATCTTCAGCATCT  DENV2_16681-I-Reverse: TTGCATGTTTCGTTCCTACTCG |
|  | DENV2_16681_**Fragment-II** | pTwist Amp MC | synthetic clonal genes | DENV2_16681-II-Forward: AACTTTTCACTAGGAGTCTTGGG  DENV2_16681-II-Reverse: CTGAGCTTCTCTGGTTGCT |
|  | DENV2_16681_**Fragment-III** | pTwist Amp MC | synthetic clonal genes | DENV2_16681-III-Forward: CAGCTCTTTTCTTATTGGTAGCACA  DENV2_16681-III-Reverse: CTCAGGGTCAATGCCAGCGCTT |
| **JEV-WT**  **wild-type** |  |  |  |  |
|  | wt-JEV-SA14_**Fragment-I** | pET-28a(+)  pCC1  pTwist Amp MC | synthetic clonal genes  *Failed**  *Failed*  *Failed* | NA |
| *Divide into 2 fragments and use synthesis of DNA* | wt-JEV-SA14_**Fragment-I-A**  wt-JEV-SA14_**Fragment-I-B** | pTwist Amp MC  pTwist Amp MC | synthetic clonal genes  synthetic clonal genes | JEV-FI-ISA-SA14-14-2-Fr: CACCCAACTGATCTTCAGCATCT  wt-SA14-Fragment-1-A-Rev: TCCTTGGCTCACAGTCCAGTGTG  wt-SA14-Fragment-1-B-Fr: GTTGGATCTTAAAAACAGCAATCAA  wt-SA14-Fragment-1-B-Rev: TTACAGTAACACCCAATGCTCCT |
|  | wt-SA14-14-2_**Fragment-II** | pET-28a(+) | synthetic clonal genes | JEV-FII-ISA-SA14-14-2-Fr: GCTTTCGCAGAGGCCAACAGTGG  JEV-FII-ISA-SA14-14-2-Rev: GCTGTCCTTCTCTGGGCAGCCC |
|  | wt-SA14-14-2_**Frgament-III** | pET-28a(+) | synthetic clonal genes | JEV-FIII-ISA-SA14-14-2-Fr: GACACTTCACTATGGGTACATGCTCCC  JEV-FIII-ISA-SA14-14-2-Rev: CTCAGGGTCAATGCCAGCGCTT |
| **JEV-CpG**  **CpG-enriched** |  |  |  |  |
|  | E-CpG-SA14-**Fragment-1A**  E-CpG-SA14-**Fragment-1B** | pTwist Amp MC  pTwist Amp MC | synthetic clonal genes  synthetic clonal genes | E-CG-UAnorm-SA14-F1A-Fr:  CACCCAACTGATCTTCAGCATCT  E-CG-UAnorm-SA14-F1A-Rv:  TTCGAGGTTCGCAGTCGAGCGTC  E-CG-UAnorm-SA14-F1B-Fr:  TTACCGTCACGCCGAACGCGCCG  E-CG-UAnorm-SA14-F1B-Rv:  GTTGGATCTTAAAAACAGCAATCAA |
|  | wt-SA14-14-2- Fragment-II | pET-28a(+) | synthetic clonal genes | JEV-FII-ISA-SA14-14-2-Fr: GCTTTCGCAGAGGCCAACAGTGG  JEV-FII-ISA-SA14-14-2-Rev: GCTGTCCTTCTCTGGGCAGCCC |
|  | wt-SA14-14-2- Fragment-III | pET-28a(+) | synthetic clonal genes | JEV-FIII-ISA-SA14-14-2-Fr: GACACTTCACTATGGGTACATGCTCCC  JEV-FIII-ISA-SA14-14-2-Rev: CTCAGGGTCAATGCCAGCGCTT |
| **WNV-WT**  **wild-type** |  |  |  |  |
|  | wt-NY99_**Fragment-I** | pET-28a(+) | synthetic clonal genes  *Failed** | NA |
| *Low copy number plasmid* | wt-NY99_**Fragment-I** | **pCC1** | synthetic clonal genes | WNV-wt-NY99-I-Forward: CACCCAACTGATCTTCAGCATCT  WNV-wt-NY99-I-Reverse: GCGAAGGACCTCCTGGGTGGC |
|  | wt-NY99_**Fragment-II** | pET-28a(+) | synthetic clonal genes | WNV-wt-NY99-II-Forward: TATAATGCTGATATGATTGACC  WNV-wt-NY99-II-Reverse: AGTGAACTCTTCTTTTGTCATC |
|  | wt-NY99_**Fragment-III** | pET-28a(+) | synthetic clonal genes | WNV-wt-NY99-III-Forward: GGTGGGGCAAAAGGACGCACCT  WNV-wt-NY99-III-Reverse: CTCAGGGTCAATGCCAGCGCTT |
| **WNV-Per**  **permuted** |  |  |  |  |
|  | NY99-Pr-E-NS1_**Fragment-I** | pET-28a(+)  pCC1  pTwist Amp MC | synthetic clonal genes  *Failed**  *Failed*  *Failed* | NA |
| *Divide into 3 fragments* | NY99-Pr-E-NS1_**Fragment-I-A**  NY99-Pr-E-NS1_**Fragment-I-B**  NY99-Pr-E-NS1_**Fragment-I-c** | NA  NA  pTwist Amp MC | *synthesis of DNA*  *synthesis of DNA*  *Failed** | See below |
| *Divide into 4 fragments and use synthesis of DNA* | NY99-Pr-E-NS1_**Fragment-I-A**  NY99-Pr-E-NS1_**Fragment-I-B**  NY99-Pr-E-NS1_**Fragment-I-C**  NY99-Pr-E-**NS1_Fragment-I-D** | NA  NA  NA  NA | *synthesis of DNA*  *synthesis of DNA*  *synthesis of DNA*  *synthesis of DNA* | Fr-NY99-Per-FI-A: CACCCAACTGATCTTCAGCATCT  Rv-NY99-Per-FI-A: ATCATGACTGCAATTCCGGTC  Fr-NY99-Per-FI-B: TGACCAGTGCTATCAATCGGC  Rv-NY99-Per-FI-B: TATGATGGAGCTGCTGGTGTG  Fr-NY99-Per-FI-C: CGGCAACTACTCCACCCAG  Rv-NY99-Per-FI-C: CGTTTCTGGGTAATACTTGTACCGAT  Fr-NY99-Per-FI-D: TGTGGAAGTGGGGTGTTCATAC  Rv-NY99-Per-FI-D: GCGAAGGACCTCCTGGGT |
|  | wt-NY99_**Fragment-II** | pET-28a(+) | synthetic clonal genes | WNV-wt-NY99-II-Forward: TATAATGCTGATATGATTGACC  WNV-wt-NY99-II-Reverse: AGTGAACTCTTCTTTTGTCATC |
|  | wt-NY99_**Fragment-III** | pET-28a(+) | synthetic clonal genes | WNV-wt-NY99-III-Forward: GGTGGGGCAAAAGGACGCACCT  WNV-wt-NY99-III-Reverse: CTCAGGGTCAATGCCAGCGCTT |
