## supplemental tables and files for "Infectious subgenomic amplicon strategies for Japanese encephalitis and West Nile viruses": Table S2 JEV and WNV genomic parameters.docx

**Japanese encephalitis virus genomic parameters**

|  | **GC3** | **Obs_ENc** | **Exp_ENc** | **Ratio_ENc** | **CP_Bias** | **Total CpG** | **Total UpA** |
| --- | --- | --- | --- | --- | --- | --- | --- |
| **JEV-WT** | 0.552 | 54.5031 | 59.931 | 0.909 | 0.005 | 58 | 33 |
| **JEV-CpG** | 0.53 | 40.5754 | 60.322 | 0.673 | -0.266 | 193 | 37 |

**The genomic region encoding envelope (E) protein**

**West Nile virus genomic parameters**

|  | **GC3** | **Obs_ENc** | **Exp_ENc** | **Ratio_ENc** | **CP_Bias** | **Total CpG** | **Total UpA** |
| --- | --- | --- | --- | --- | --- | --- | --- |
| **WNV-WT** | 0.556 | 50.1376 | 59.837 | 0.838 | 0.038 | 49 | 36 |
| **WNV-Per** | 0.556 | 51.8292 | 59.837 | 0.866 | 0.031 | 48 | 36 |

**The genomic region encoding envelope (E) protein**

|  | **GC3** | **Obs_ENc** | **Exp_ENc** | **Ratio_ENc** | **CP_Bias** | **Total CpG** | **Total UpA** |
| --- | --- | --- | --- | --- | --- | --- | --- |
| **WNV-WT** | 0.556 | 55.7047 | 59.848 | 0.931 | 0.012 | 37 | 30 |
| **WNV-Per** | 0.556 | 56.926 | 59.848 | 0.951 | 0.038 | 37 | 30 |

**The genomic region encoding non-structural 1 (NS1) protein**

**GC3** - Guanine and cytosine content at the third codon position.

**Obs_Enc** – Effective number of codons, observed.

**Exp_Enc** – Effective number of codons, expected.

**Ratio_Enc** – Effective number of codons, ratio.

**CP_Bias** – Codon pair bias.

**Total UpA** – Total number of UpA dinucleotides.

**Total CpG** – Total number of CpG dinucleotides.
